# Malaria parasite evades mosquito immunity by glutaminyl cyclase mediated protein modification

**DOI:** 10.1101/2021.07.23.453408

**Authors:** Surendra Kumar Kolli, Alvaro Molina-Cruz, Tamasa Araki, Fiona J.A. van Geurten, Jai Ramesar, Severine Chevalley-Maurel, Hans J Kroeze, Sacha Bezemer, Clarize de Korne, Roxanne Withers, Nadia Raytselis, Angela F. El Hebieshy, Robert Q Kim, Matthew Child, Soichiro Kakuta, Hajime Hisaeda, Hirotaka Kobayashi, Takeshi Annoura, Paul J. Hensbergen, Blandine M. Franke-Fayard, Carolina Barillas-Mury, Ferenc A. Scheeren, Chris J. Janse

**Author notes:** Present address: Center for Global Health and Infectious Diseases Research, College of Public Health, University of South Florida; Tampa, Florida, USA.

## Abstract

Glutaminyl cyclase (QC) modifies N-terminal glutamine or glutamic acid residues of target proteins into cyclic pyroglutamic acid (pGlu). Here, we report the biochemical and functional analysis of *Plasmodium* QC. We show that *Plasmodium* sporozoites of QC-null mutants are recognized by the mosquito immune system and melanized when they reach the hemocoel. Sporozoite numbers in salivary glands are also reduced in mosquitoes infected with QC-null or QC catalytically-dead mutants. This phenotype can be rescued by genetic complementation or by disrupting mosquito hemocytes or melanization immune responses. Mutation of a single QC-target glutamine of the major sporozoite surface protein (CSP) also results in immune recognition of sporozoites. These findings reveal QC-mediated post-translational modification of surface proteins as a major mechanism of mosquito immune evasion by *Plasmodium* sporozoites.

## Introduction

N-terminal modification of glutamine or glutamic acid residues to pyroglutamic acid (pGlu; 5-oxo-L-proline) is a posttranslational modification (PTM), catalyzed by glutaminyl cyclases (QCs) found in eukaryotes and prokaryotes (*1, 2*). Two evolutionary unrelated classes exist; mammalian QCs and QCs of bacteria, plants and parasites, which share no sequence homology, supporting a different evolutionary origin(*3*). Mammalian cells can express two forms, the secreted glutaminyl‐peptide cyclotransferase (QPCT) or its iso-enzyme (QPCTL), localized in the Golgi complex. pGlu is implicated in maturation and stabilization of mammalian proteins such has neuropeptides and cytokines (*3, 4*). QC activity has been associated in humans with pathological processes such as amyloidotic diseases (*3–6*) and QPCTL is critical for pGLu formation on CD47, facilitating myeloid immune evasion (*7, 8*). The physiological function of QCs in plants, bacteria and parasites remains poorly characterized. Several pGlu-containing proteins of plants are involved in defense reactions against pathogens (*3*). Here we report the biochemical and functional analysis of QC of malaria parasites.

## Results

### *Plasmodium* QC shows sequence homology to bacterial and plant QCs, has cyclase activity and is expressed in transmission stages

A single gene encoding a glutaminyl cyclase (QC), named glutaminyl-peptide cyclotransferase, has been identified by electronic annotation in all sequenced *Plasmodium* genomes. *Plasmodium* QCs share 70-76% sequence similarity and 50-54% identity and contain a transmembrane domain (**Fig. S1**). QC of the human malaria parasite *P. falciparum* (*Pf*QC) shows 21-27% identity to QCs of various bacteria and the plant *Carica papaya* (*Cp*QC)(*1, 9–11*) (**Fig. S2**). All 9 amino acids of the catalytic site of bacterial and plant QCs(*1*) are conserved in *Plasmodium* QC and the secondary structure of *Pf*QC, threaded against *Cp*QC and bacterial QCs, reveal 22 conserved β-sheets (**Fig. S2**). A *Pf*QC 3D-homology model, built against *Cp*QC and bacteria QCs, predicted 5 conserved blades of antiparallel β-sheets and a highly similar tertiary structure of *Pf*QC and *Cp*QC **(Fig. 1A,B; Fig. S2**). Recombinant wild type *Pf*QC has cyclase activity in a two-step enzyme-activity assay that is not present in a catalytically-dead *Pf*QC (*Pf*QC^CD^), in which two mutated, conserved amino acid residues in the catalytic site were mutated (**Fig. 1C**).

**Fig. 1.**
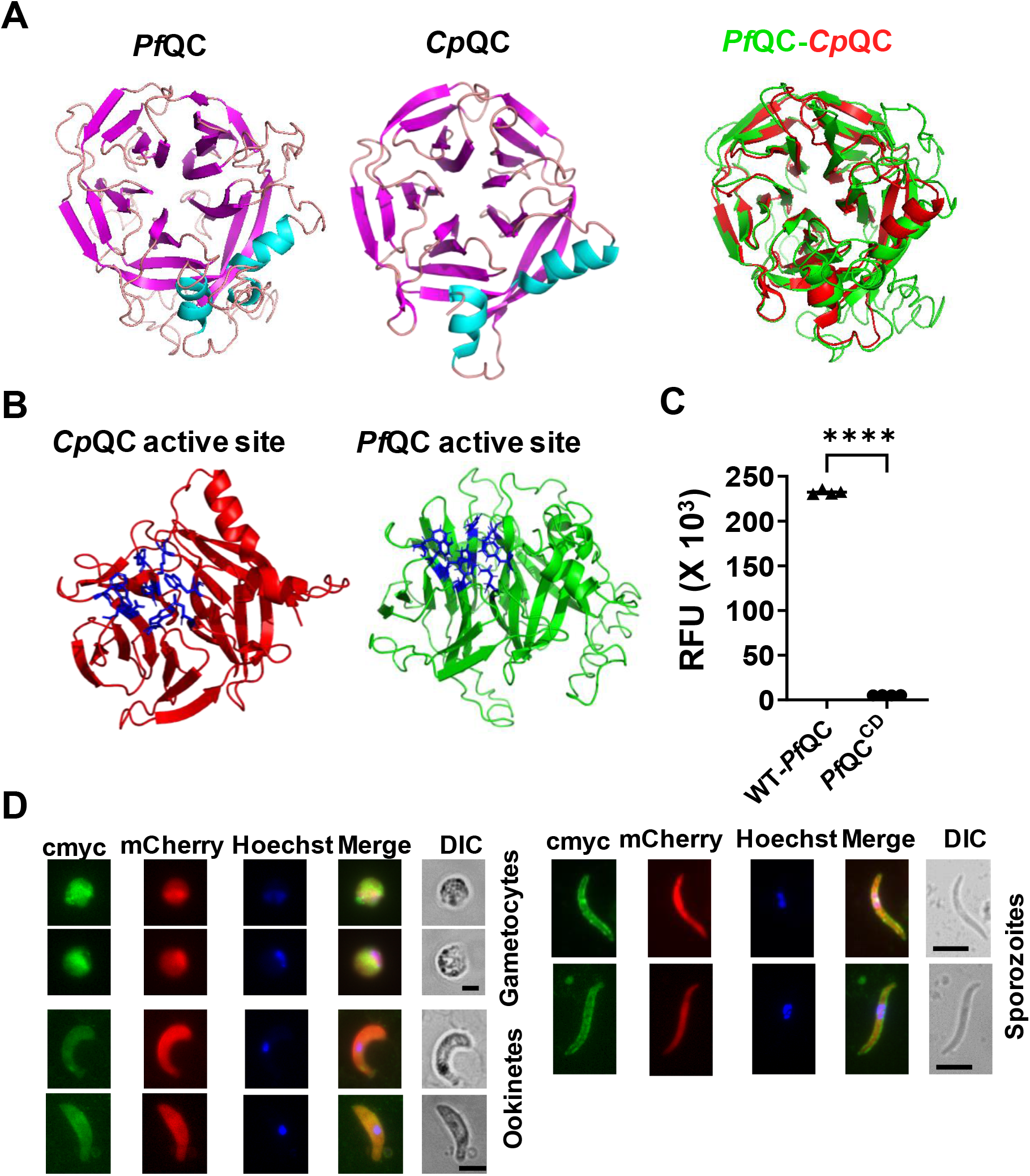
3D-homology model, cyclase activity and expression of *P. falciparum* glutaminyl cyclase (OC) **A**. Three dimensional (3D) homology model of *P. falciparum* QC (*Pf*QC), generated against the resolved QC structures from *C. papaya* and *Z. mobilis*, and the 3D template structure of *C. papaya* QC (*Cp*QC). Structures are visualized using Pymol. Helices – cyan; sheets -magenta; loops - brown. An overlay is shown of *PfQC* (green) and *Cp*QC (red). **B**. Visualization of the *Cp*QC catalytic site (blue), composed of active site residues F22, Q24, F67, E69, W83, W110, N155, W169 and K225 and the predicted *Pf*QC (blue) catalytic site, based on the catalytic site residues conserved with the *Cp*QC catalytic site (F107, Q109, F152, E154, Y174, Y200, N269, F285 and K349; **Fig. S2**). **C**. Cyclase activity of *Pf*QC (relative fluorescence units, RFU), in an enzyme activity assay using recombinant wild type (QC^WT^; n=4) and cyclase-dead *Pf*QC (QC^CD^; n=4), containing two point mutations in the active site, F107A and Q109A. **D**. QC expression in gametocytes, ookinetes and sporozoites, visualized by staining with anti-cmyc antibodies of fixed *Pbqc::cmyc* parasites that express cmyc-tagged QC and mCherry. Nuclei stained with Hoechst 33342. Scale bar: 5 μm.

Published expression data indicate that QC is expressed in gametocytes, ookinetes, oocysts, and sporozoites, *Plasmodium* mosquito stages critical for disease transmission (**Fig. S3**). Gametocytes, formed in the blood of the vertebrate host, develop into gametes when taken up by a mosquito and fuse to form zygotes. Zygotes transform into ookinetes that cross the midgut epithelial cells and develop into oocysts in the hemocoel (*12*). Sporozoites, formed inside oocysts, are released into the hemocoel and invade salivary glands for transmission to a new vertebrate host (*13, 14*). QC expression in transmission stages was confirmed by analyzing a transgenic rodent malaria parasite (*P. berghei; Pbqc::cmyc*), expressing a cmyc-tagged QC (**Fig. 1D, Fig. S4; Table S2**).

### Absence of QC leads to melanization of oocysts and sporozoites and reduced sporozoite numbers

To analyze the QC function *in vivo*, QC-null mutants of the rodent parasite *P. berghei* and the human parasite *P. falciparum* were generated. QC-null mutants were created by deleting the *qc* gene from the parasite genome using standard methods of genetic modification. Blood-stages of three *P. berghei* QC-null mutants (*Δqc1; Δqc2; Δqc-G)* showed wild-type (WT) asexual growth rate and gametocyte production (**Fig. S5, Table S2**). When fed to *A. stephensi*, QC-null parasites produced similar numbers of oocysts as WT parasites (**Fig. 2A; Table S2)**. In contrast, the number of QC-null salivary gland sporozoites (sg-sporozoites) was significantly reduced compared to WT (Mann-Whitney test *Δqc1* p= 0.0095, *Δqc2* p= 0.0043 and *Δqc-G* p=0.0238; **Fig. 2B**). Closer examination of maturing oocysts, revealed the presence of aberrant, dark-colored oocysts in 55-65% of QC-null-infected mosquitoes (1-85 dark-colored oocysts/mosquito), while such oocysts were absent in WT-infected mosquitoes (**Fig. 2A,C,D,E**). No dark-colored oocysts were detected before day 10 post-infection (p.i.). Light-microscopy analysis of oocysts between day 14 and 21 showed the presence of aberrant, enlarged, dark-colored sporozoites, either still inside or during release from oocysts (**Fig. 2F**, **Fig. S6**). The dark-colored oocysts and sporozoites indicate the formation of melanin, an insect immune effector response(*15*), resulting in melanotic encapsulation of pathogens. Scanning electron-microscopy with energy-dispersive X-Ray analysis (SEM-EDX) revealed the presence of sulfur in the dark-colored parasites (**Fig. 3**), indicative of pheomelanin(*16*) that is a constituent of melanized ookinetes(*17*). The dark-colored material, hereinafter referred to as melanin, was mainly located inside (rupturing) oocysts and on sporozoites and distinct melanin deposition on the oocyst capsule was not observed (**Fig. S6)**. Transmission electron microscopy confirmed melanin deposition mainly inside oocysts, often surrounding sporozoites (**Fig. S7**).

**Fig. 2.**
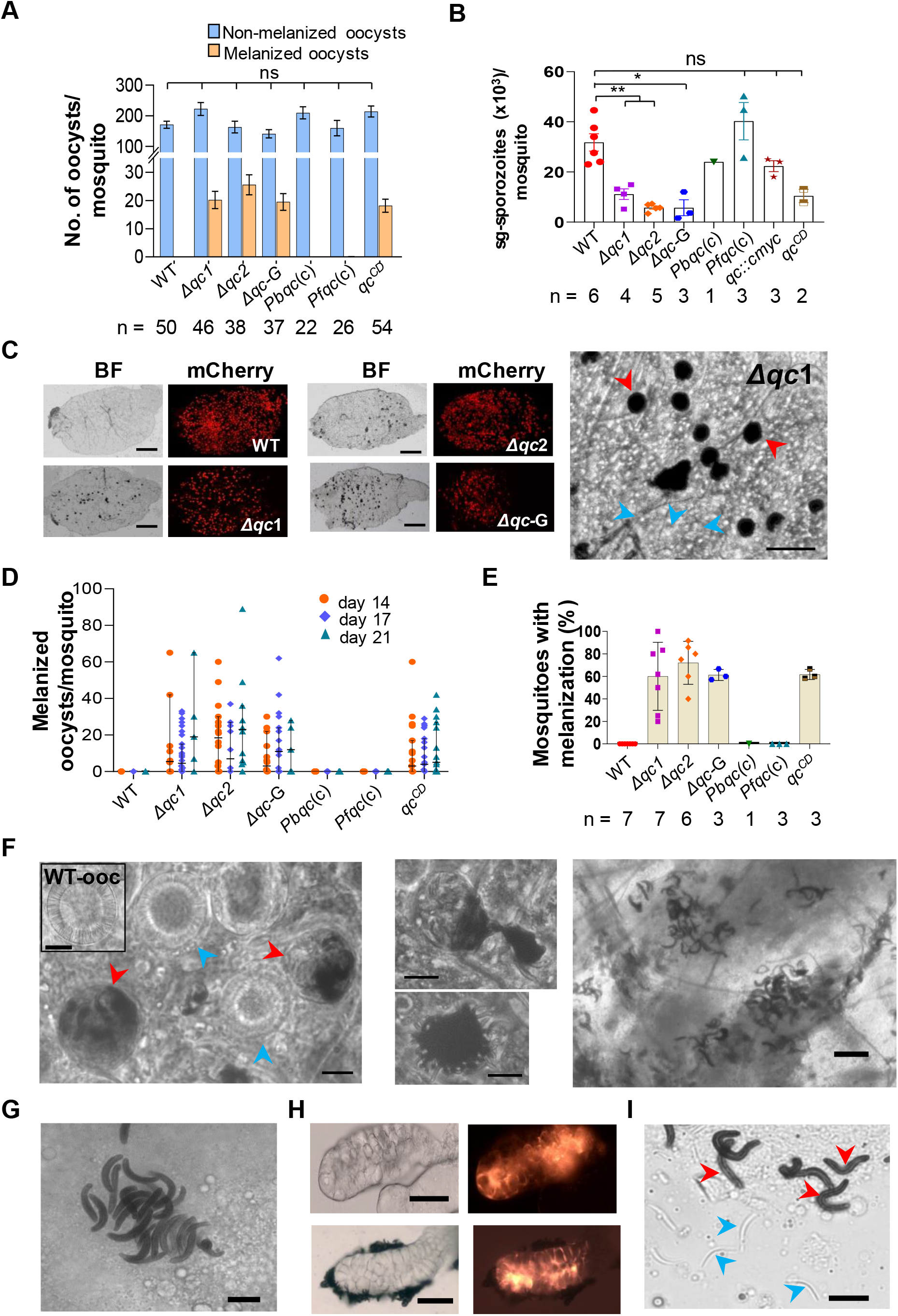
Melanization of oocysts and sporozoites in mosquitoes infected with *P. berghei* QC-null mutants and reduced sporozoite numbers in their salivary glands. **A**. Number of melanized and non-melanized oocysts per mosquito (mean ± SEM) in *A. stephensi* mosquitoes (n=number of mosquitoes) infected with QC-null mutants (*Δqc*), QC-null mutants complemented with *P. berghei* or *P. falciparum* QC (*Pbqc*(c) and *Pfqc*(c)) or infected with a QC catalytically-dead mutant (*qc*^*CD*^). WT = wild type. ns – not significant (Mann-Whitney test; statistical significance is shown relative to WT; P-values are shown in **Table S3**). **B**. Number of salivary gland (sg) sporozoites per mosquito (n=number of experiments; 60-80 mosquitoes/experiment) infected with different *qc* mutants (see **a** for mutant names). Mean and standard deviation (3-6 exp. for QC-null mutants (*Δqc*) and WT, 6 exp. For other mutants 2-4 experiments. ** P<0.005, *P<0.5 ns – not significant; (Mann-Whitney test; statistical significance is shown relative to WT. P values are shown in **Table S3**). **C**. Melanized oocyst in mosquito midguts (day 14), infected with three QC-null mutants (*Δqc*). Brightfield (BF) and fluorescence images of mCherry-expressing oocysts. WT - wild type. Scale bar: 200 μm. Right panel: Melanized (red arrows) and non-melanized (blue arrows) oocysts in a *Δqc*1 infected mosquito (day 14). Scale bar: 100 μm. **D**. Number of melanized oocysts per individual mosquito on day 14, 17 and 21 post infection of mosquitoes (n=15-20) with different *qc* mutants (see **A** for mutant names). Number mosquitoes on day 14, 17 and 21: WT (18, 21, 18); *Δqc1* (10, 32, 5); *Δqc2* (16, 10, 12); *Δqc-G* (14, 18, 5); *Pbqc*(c) (28, 11, 18); *Pfqc*(c) (16, 19, 22); *qc*^*CD*^ (17, 24, 15). **E**. Percentage of mosquitoes with melanized oocysts (n=number of experiments; 30-40 mosquitoes/experiment) at day 14 after infection with different *qc* mutant parasites (see **A** for mutants names). No melanized oocysts in WT and complemented *Pbqc*(c) and *Pfqc*(c) parasites. **F**. Examples of (partly) melanized QC-null oocysts showing melanization of sporozoites still inside or in the process of oocyst egress (left and middle panel). Inset: wild type oocyst (WT-ooc) with sporulation (day 14). Scale bar: 20 μm. Right panel: melanized QC-null sporozoites present in the hemocoel (abdominal region; day 21). Scale bar: 25 μm. **G**. Cluster of melanized (dark colored, enlarged) sporozoites in the hemocoel of QC-null infected mosquito (day 21). Scale bar: 10 μm. **H**. Brightfield and fluorescence images of salivary glands isolated at day 21 from mosquitoes infected with WT (upper panel) and QC-null parasites (lower panel). Both parasite lines express mCherry (right panels). Melanized, mCherry negative sporozoites cluster at the periphery of the salivary gland of a QC-null infected mosquito. Scale bar: 50 μm. **I**. Partially crushed salivary gland (day 21) of a QC-null infected mosquito showing melanized (red arrows) and non-melanized (blue arrows) sporozoites. Scale bar: 10 μm.

**Fig. 3.**
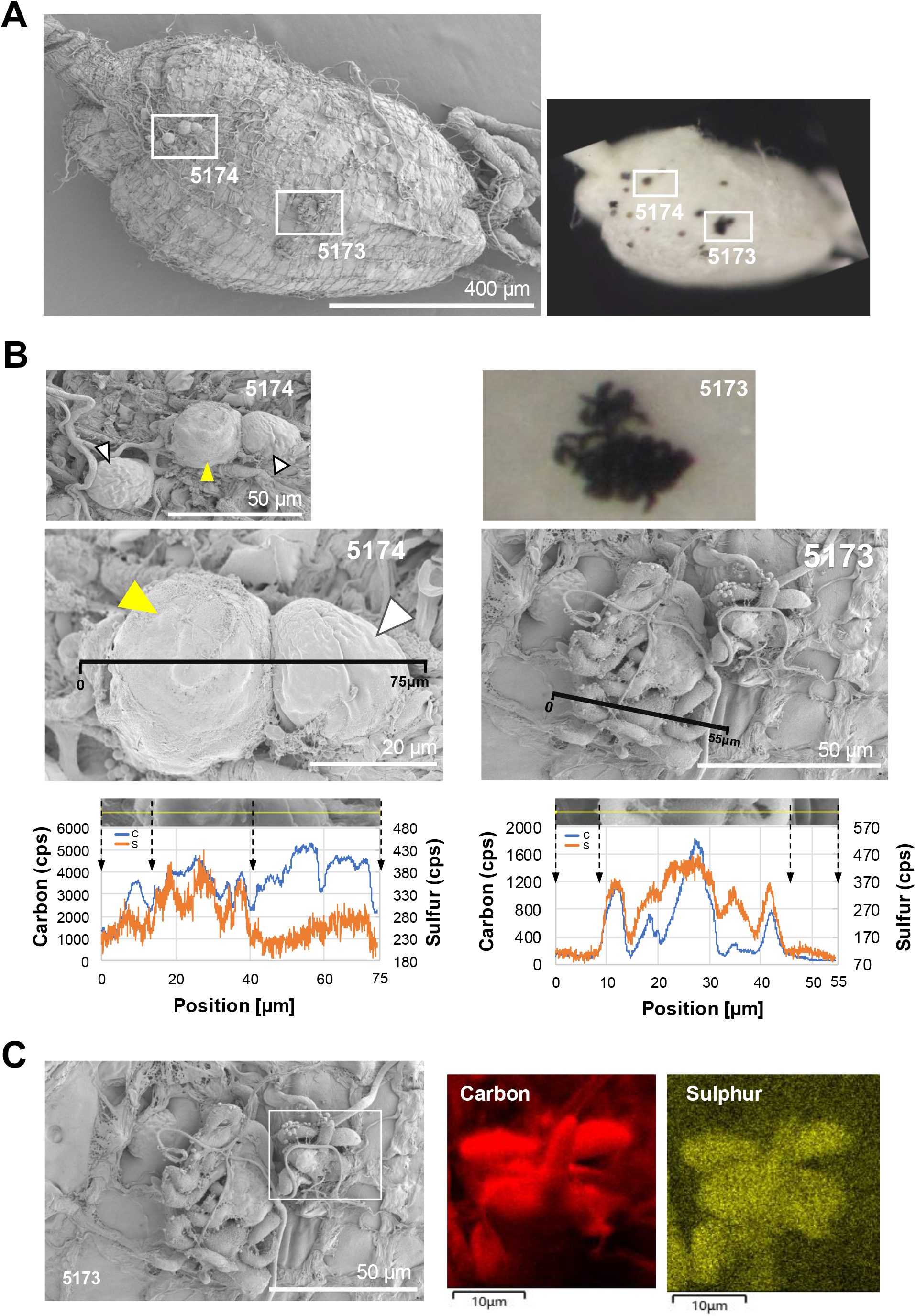
Scanning Electron Microscopy with Energy Dispersive X-Ray Analysis (SEM-EDX) of melanized and non-melanized QC-null oocysts and sporozoites. **A**. SEM (left) and bright-field (right) identification and selection of oocysts (5174) and sporozoites (5173) in a midgut of an *An. stephensi* mosquito infected with *PbΔqc*1 (day 14) **B**. Energy dispersive X-ray (EDX) line scan of elements Carbon (blue) and Sulfur (orange) together with SEM images (count per seconds: CPS) showing the presence of sulfur in melanized oocysts (left panel; dark-colored oocyst in 5174; see **A**) and in melanized, dark-colored sporozoites (right panel; 5173; see **A**) Black lines represent EDX line scan positions. Yellow arrow head indicate melanized black oocyst, and white arrow heads non-melanized oocysts (5174). Cluster of melanized sporozoites (5173) **C**. Energy dispersive X-ray (EDX) mapping analysis of Carbon and Sulfur together with SEM image of melanized sporozoites (5173; see **A**).

At day 21 p.i. non-motile, enlarged sporozoites covered by melanin were found in the hemocoel, often in clusters or attached to salivary glands (**Fig. 2G,H,I**). In contrast, most QC-null sporozoites inside salivary glands had a WT-like morphology, i.e. long and slender, without signs of melanization (**Fig. 2I, Fig. S6**). The motility of these QC-null sg-sporozoites and their infectivity to cultured hepatocytes was not significantly different from WT sg-sporozoites (**Fig. S8,9;**P values in **Table S3**). These observations reveal viability of QC-null sporozoites that invade salivary glands and indicate that the reduced sg-sporozoites numbers result from immune recognition and melanization of sporozoites while in transit to salivary glands. The melanization phenotype of QC-null parasites was rescued by genetic complementation through re-introduction of the *P. berghei* or the *P. falciparum qc* gene into the disrupted *qc* locus of *P. berghei* QC-null mutants (**Fig. S10**). Complemented parasites showed WT-like oocyst and sporozoite development without signs of melanization and sg-sporozoite numbers were also restored to WT numbers (**Fig. 2A,B,D,E, Fig. S10, Table S2**).

The importance of QC cyclase activity to prevent melanization, was analysed using a *P. berghei* mutant expressing a cyclase-dead QC containing two point mutations in the catalytic site (**Fig. S11**). Similar to QC-null mutants, melanization was observed in this mutant with reduced sg-sporozoite numbers (**Fig. A,B,D,E, Fig. S11, Table S2**), revealing that cyclase-activity is critical for preventing melanization.

Melanization of oocyst and sporozoites also occurred in *P. falciparum* QC-null mutants, with melanized oocysts in 25-34% of infected mosquitoes (1-20 melanized oocysts per mosquito), whereas melanized parasites were absent in WT-infected mosquitoes (**Fig. 4**). QC-null mutants similar asexual growth rate, gametocyte production (**Fig. S12**) and oocysts numbers as WT parasites, but showed modest reduction in the number of sg-sporozoites, that was not statistically significant (Mann-Whitney p=0.133; **Fig. 4**). Combined, we show that absence of QC results in immune recognition of oocyst and sporozoites of rodent and human malaria parasites

**Fig. 4.**
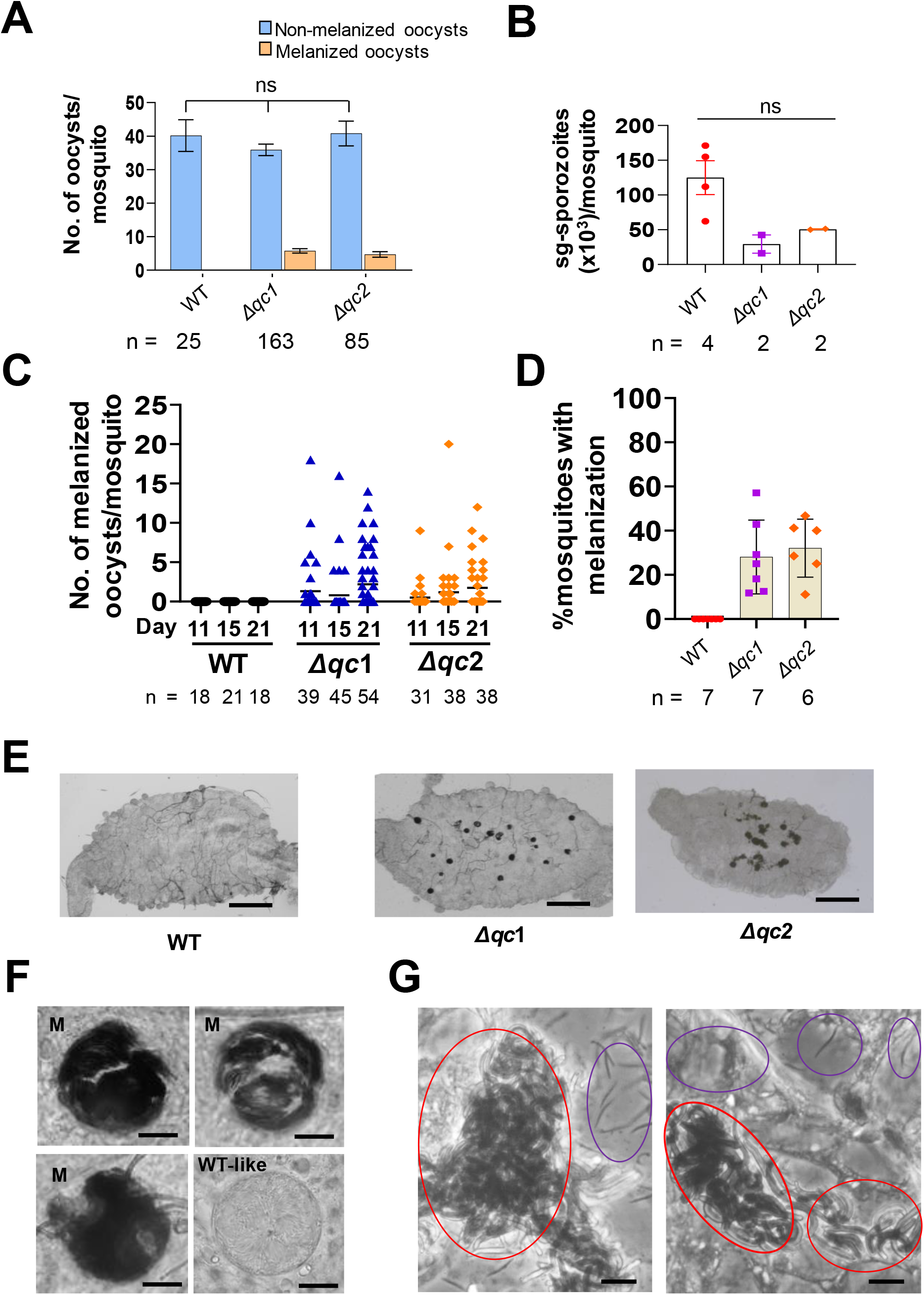
Melanization of oocysts and sporozoites of *P. falciparum* QC-null mutants. **A**. Number of melanized and non-melanized oocysts per mosquito (mean ± SEM) in *A. stephensi* mosquitoes (n= number of mosquitoes) infected with QC-null mutants (*Δqc*) or wild type (WT) parasites. ns-not significant (Mann-Whitney test, statistical significance is shown relative to WT. P values are shown in **Table S3**). **B**. Number of salivary gland (sg) sporozoites per mosquito (n=number of experiments; 60-80 mosquitoes/experiment), infected with QC-null mutants (*Δqc*) or WT parasites. Mean and standard deviation (2 exp. for QC-null mutants (*Δqc*) and WT-4 exp. ns - not significant (Mann-Whitney test, statistical significance is shown relative to WT. P values are shown in **Table S3**). **C**. Number of melanized oocysts per individual mosquito on day 11, 15 and 21 after infection with QC-null mutants or WT parasites (n= number of mosquitoes). **D**. Percentage of mosquitoes with melanized oocysts (n=number of experiments; 30-40 mosquitoes/experiment) at day 21 after infection with QC-null mutants or WT parasites. **E**. Melanized (dark-coloured) oocyst in midguts of mosquitoes (day 11) infected with QC-null mutants. No melanized oocysts were observed in WT parasites. Scale bar: 200 μm. **F**. Examples of (partly) melanized QC-null oocysts (M, day 15) showing melanization of sporozoites still inside or in the process of oocyst egress and an oocyst with typical features of WT sporozoite formation (WT-like). Scale bar: 20 μm. **H**. Melanized (red circles) and non-melanized, WT-like (blue circles) QC-null sporozoites obtained from a salivary gland (under a coverslip) isolated from an infected mosquito (day 21). Scale bar: 10 μm.

### Silencing of mosquito immune responses results in reduced melanization and increased sg-sporozoite numbers in QC-null-infected mosquitoes

Mosquito innate immune responses in the hemocoel can detect and eliminate parasites through multiple effector mechanisms (*18, 19*). Immune responses that target the ookinete stage involve activation of nitration responses by invaded midgut cells (*19*) and local release of hemocyte-derived microvesicles that promote complement activation (*20*). The thioester-containing protein 1 (TEP1), a final effector of mosquito complement, binds to the ookinete surface thereby initiating the formation of a complex that kills the parasite through lysis or melanization (*21*). A second TEP1-independent late-phase response, involving the STAT pathway (*22*) and hemocytes (*23*), targets the oocyst stage and hemocyte-mediated phagocytosis of sporozoites has also been reported (*24*) Whereas substantial melanization-associated killing of ookinetes has been described (*18*), extensive oocyst- or sporozoite melanization has not been reported. The melanization of QC-null parasites could result from either recognition of viable parasites as non-self by the immune system or a general mechanism of disposal of aberrant/dead parasites (*25*). To examine whether abnormal/dead QC-expressing parasites would trigger a similar melanization response as QC-null parasites, we re-analyzed oocysts of four published *P. berghei* mutants that produce WT-like oocyst numbers, but have severe defects in sporozoite formation (CSP- and ROM3-null mutants (*26, 27*)), egress from oocysts (CRMP4-null mutant (*28*)) or invasion of salivary glands (TRAP-null mutant (*29*)). Despite these defects during sg-sporozoite formation, no melanization was observed (**Fig. S13**), indicating that presence of aberrant parasites is not a general trigger for melanization. To further analyze whether viable QC-null sporozoites are recognized by hemocytes and melanized, we disrupted hemocyte function by systemic injection of polystyrene beads as described (*30*). Bead injection had no significant effect on the number of sg-sporozoites in WT-infected mosquitoes (**Fig. 5A**), but significantly increased the number of QC-null sg-sporozoites, indicating that lack of QC-activity results in active elimination of viable sporozoites by hemocytes (**Fig. 5B**, Mann-Whitney test; p<0.01). The potential participation of the complement-like system in immune recognition of QC-null sporozoites was evaluated by silencing LRIM1, a stabilizer of TEP1 (*31*). Disrupting the complement system had no significant effect on the number of QC-null sg-sporozoites (**Fig. 5C)**. In contrast, silencing the serine protease CLIPA8, a critical activator of melanization (*32*), reduced the percentage of QC-null infected mosquitoes with melanized oocysts from 75% to 5% (**Fig. 5D**; *X*^2^, p<0.0001) and significantly increased the number of sg-sporozoites (**Figure 5E**; Mann-Whitney, p=0.0001). Taken together, these findings demonstrate that hemocytes detect QC-null sporozoites as non-self and eliminate them through melanization.

**Fig. 5.**
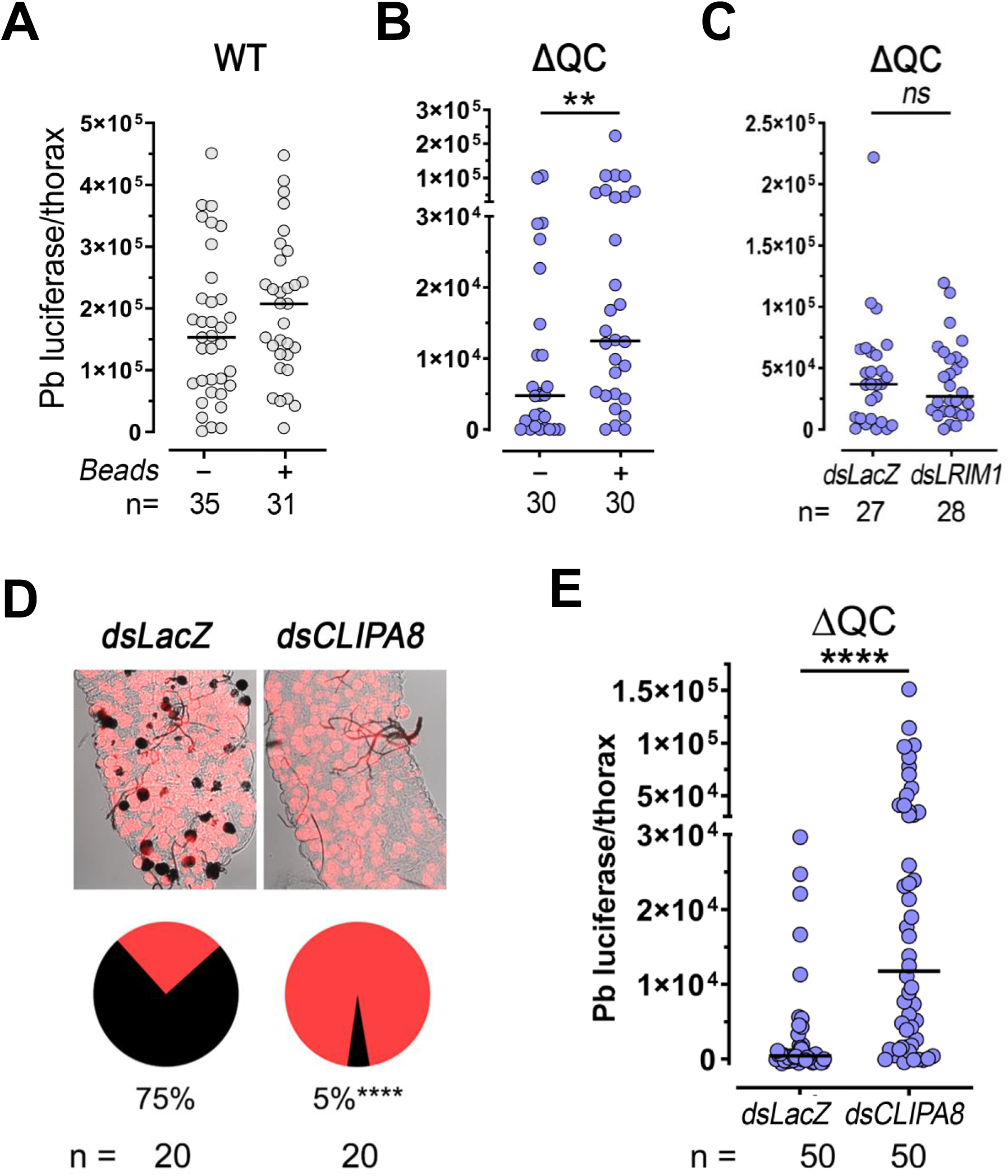
Silencing of mosquito immune responses results in increased sporozoite numbers in salivary glands of mosquitoes infected with *P. berghei* QC-null mutants. **A**. Sporozoite density (luciferase activity in individual mosquito thoraces at day 23 post infection (p.i.) after systemic injection of polystyrene beads in the thorax (day 12 p.i.) of mosquitoes infected with wild type (WT) parasites (n=number of mosquitoes). ns – not significant (Mann-Whitney test, P >0.05). **B**. Sporozoite density (luciferase activity in individual mosquito thoraces; day 23 p.i.) after systemic injection of polystyrene beads in the thorax (day 12 p.i.) of QC-null (*Δqc2*) infected mosquitoes (n=number of mosquitoes). ** P<0.01 (Mann-Whitney test). **C**. Sporozoite density (luciferase activity in individual mosquito thoraces; day 23 p.i.) after injection of dsRNA for LRIM1 (ASTE000814) (dsLRIM1) or LacZ (dsLacZ; control) (day 12 p.i.) of QC-null mutant (*Δqc2)* infected mosquitoes (n=number of mosquitoes). ns – not significant (Mann-Whitney test, P >0.05). **D**. Representative midguts and percentage of mosquitoes with melanized parasites (23 day p.i) after injection of dsRNA for CLIPA8 (ASTE009395) (dsCLIPA8) or LacZ (dsLacZ; control) (day 12 p.i.) of QC-null mutant (*Δqc2)* infected mosquitoes (n=number of mosquitoes). **** P<0.0001 (Chi-square test). **E**. Sporozoite density (luciferase activity in individual mosquito thoraces; day 23 p.i.) after injection of dsCLIPA8 or dsLacZ (control) (day 12 p.i.) of QC-null mutant (*Δqc2)* infected mosquitoes (n=number of mosquitoes). **** P<0.0001 (Mann-Whitney test).

### QC-null oocysts are not melanized when sporozoite formation or egress is prevented

The absence of melanized QC-null oocysts before day 10 p.i. and absence of distinct deposition of melanin on the oocyst capsule may suggest that QC-null sporozoites, either still inside rupturing oocysts or after release into the hemocoel, are specifically recognized by the immune system. If this is the case, abolishing sporozoite formation inside QC-null oocysts or blocking egress of OC-null sporozoites would prevent oocyst melanization. To test this hypothesis we deleted genes encoding proteins involved in sporozoite formation (CSP(*26*), ROM3 (*27*)) or in sporozoite egress (ECP1 (*33*), CRMP4 (*28*)) in *P. berghei* QC-null parasites (**Fig. S14)**. These ‘double knock-out’ QC-null mutants produced WT numbers of oocysts; however, melanized oocysts were completely absent (**Figure 6A; Fig. S15**). In contrast, melanized oocysts/sporozoites were observed in 79% of mosquitoes infected with a ‘double knock-out’ QC-null mutant lacking the *trap*(*29*) gene that releases sporozoites into the hemocoel that cannot invade the salivary glands (1-60 melanized oocysts/mosquito) (**Fig. 6A; Fig. S15**). These observations confirm that melanization only occurs when QC-null oocysts rupture and sporozoites are released into the hemocoel.

**Fig. 6:**
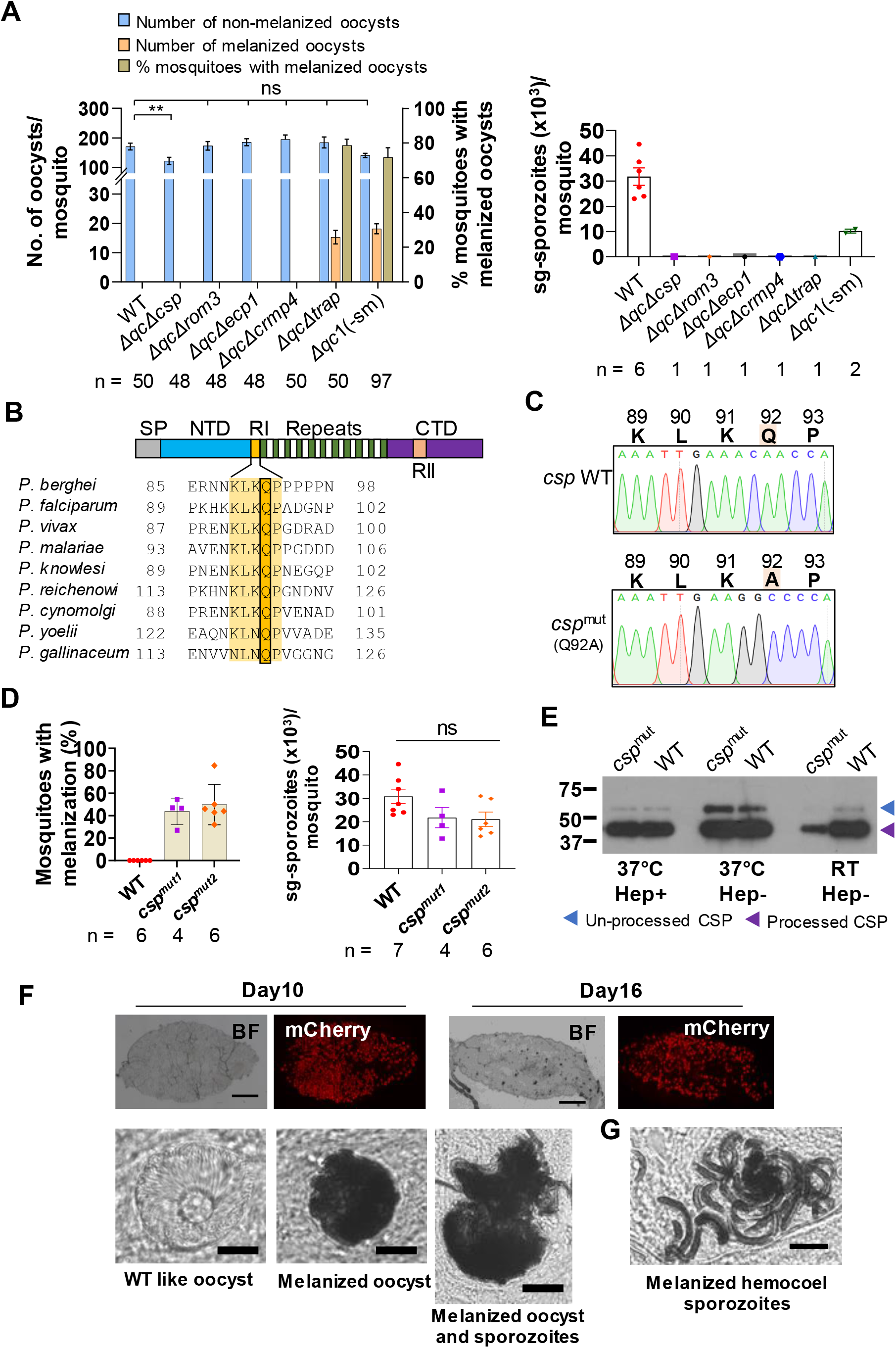
Absence of melanization of QC-null oocysts in the absence of sporozoite egress and melanization of oocysts and sporozoites that express the sporozoite surface protein CSP with a mutated QC-target glutamine. **A**. Left panel: Number of melanized and non-melanized oocysts per mosquito (mean ± SEM) in *A. stephensi* mosquitoes (n=40-50) infected with a QC-null mutant (*Δqc1*(-sm)), wild type (WT) and QC-null mutants (*Δqc*Δ*csp*, *ΔqcΔrom3*, *Δqc*Δ*ecp1, Δqc*Δ*crmp4*, and *ΔqcΔtrap*) lacking expression of CSP and ROM3 (no sporozoite formation), CRMP4 and ECP1 (no sporozoite egress), and TRAP (sporozoite egress); n=number of mosquitoes. Right y-axis shows the percentage of mosquitoes with melanized oocysts (3 experiments for all mutants except for *Δqc1*(-sm), n=4). Right panel: Number of salivary gland (sg) sporozoites per mosquito infected with the different mutants as shown in the left panel (n=number of experiments; 60-80 mosquitoes/experiment). ns - not significant (Mann-Whitney test, statistical significance is shown relative to WT. P values are shown in **Table S3**) **B**. Schematic showing different regions of circumsporozoite protein. Yellow shading: region I (RI). Black box: the conserved glutamine in R1 (KLKQP) of CSP from different *Plasmodium* species. RI, located just before the repeat region, is highly conserved and contains the cleavage site (*38*). **C**. Sanger-sequence DNA chromatograms of the PCR fragment from genomic DNA of both WT and *csp*^mut^ parasites, confirming the replacement of glutamine with alanine (Q92A; highlighted) in RI of *csp*^mut^. **D**. Left panel: Percentage of mosquitoes with melanized oocysts (n=number of experiments; 30-40 mosquitoes/experiment) at day 16 after infection with *csp*^mut^. No melanized oocysts were observed in mosquitoes infected with WT. Right panel: Number of salivary gland (sg) sporozoites per mosquito at day 21 (n=number of experiments; 60-80 mosquitoes/experiment) infected with WT, *csp*^mut1^ and *csp*^mut2^. Mean and standard deviation; ns – not significant; (Mann-Whitney test; statistical significance is shown relative to WT. P values are shown in **Table S3**). **E**. Western blotting of *P. berghei* sporozoite lysates showing CSP expression and processing in wild type (WT) and *csp*^mut^ parasites, expressing mutated CSP (Q92A), using anti-CSP antibody 3D11, recognizing *P. berghei* repeats. Processing of CSP at the sporozoite surface results in a full length (55kD) and a processed form of 45kD (*36, 77*). WT and *csp*^mut^ sporozoites were incubated with or without heparin at 37°C or room temperature (RT) for 10 min as it has been shown that CSP also undergoes processing when the sporozoite is in contact with heparan sulphate proteoglycans (*78*). **F**. Upper panel: midguts of mosquitoes with mCherry expressing *csp*^mut^ oocysts, showing the presence of melanized (dark-colored) oocysts (scale bar 200 μm). Lower panel: brightfield light microscope images of *csp*^mut^ oocysts (day 21) showing normal, WT-like (WT-ooc, left) and melanized (middle and right) oocysts. Right: Sporozoites melanized during the process of egress from the oocyst (scale bar: 20 μm). **G**. Melanized *csp*^mut^ sporozoites in the hemocoel (abdominal region; day 21). Scale bar: 10 μm.

### Sporozoites expressing the circumsporozoite surface protein (CSP) with a mutated QC-target glutamine are recognized by the mosquito immune system

Circumsporozoite surface protein (CSP) is the most abundant protein on the sporozoite surface (*13, 14*). We therefore hypothesized that CSP could be a prime target for immune recognition of QC-null parasites. We analyzed published proteomes for pGlu modification of CSP and identified multiple CSP peptides with pGlu at the N-terminus with the majority (>90%) having pGlu modification of the glutamine (Q) located in region 1 of CSP (**Fig. S16)**. This short, five amino acid region (KLKQP), contains a proteolytic cleavage site (*34, 35*) and the glutamine is conserved across different *Plasmodium* species (**Fig. 6B**). CSP cleavage is known to occur at the sporozoite surface (*36, 37*) and this processing step is essential for host cell invasion (*38*). To establish whether CSP is a main target for immune recognition of QC-null sporozoites, two independent *P. berghei* lines (*csp*^mut1,2^) were generated that express a mutated CSP with glutamine in region 1 replaced with alanine (Q92A), thereby preventing pGlu formation (**Fig. S17; Fig. 6C**). Both *csp*^mut^ mutants produced similar numbers of oocysts as WT parasites. No melanization of *csp*^mut^ oocysts was observed until day 10,whereas at day 14-16 p.i., melanized oocysts were detected in 26-48% of *csp*^mut^-infected mosquitoes (1-35 melanized oocysts/mosquito), mainly rupturing oocysts (**Fig. 6D,F**). Clusters of melanized sporozoites were found in the hemocoel (**Fig. 6F**) and *csp*^mut^ sg-sporozoite numbers were reduced compared to WT, although not statistically significant (**Fig. S17**; p values 0.073 and 0.073, for *csp*^mut1^ and *csp*^mut2^; Mann-Whitney test). We confirmed CSP processing in both WT and *csp*^mut^ sg-sporozoites by Western analysis (**Fig**. **6E**) and *csp*^mut^ sg-sporozoites showed WT-like infectivity to cultured hepatocytes (**Fig. S17**). These observations demonstrate that the absence of the single glutamine residue in CSP region 1 results in immune recognition of viable/infective sporozoites in the hemocoel, resulting in melanization. Combined, our observations indicate that malaria parasites evade mosquito immune recognition by QC-mediated post-translational modification of CSP.

## Discussion

Here, we present the functional characterization of a parasite QC and show that *Plasmodium* QC plays a major role in sporozoite evasion of mosquito immune recognition. No additional phenotypes of QC-null parasites were observed in other *Plasmodium* stages in the mosquito. The lack of melanization of QC-null oocysts, in the absence of sporozoite egress, demonstrates that sporozoites are the principal target for immune recognition of QC-null parasites. While it has been shown that the ookinete surface protein P47 mediates evasion of mosquito immunity by disrupting epithelial nitration (*39*), parasite factors that are implicated in immune evasion of sporozoites had not been reported. Our studies demonstrate that hemocytes are major effectors of the immune attack of QC-null sporozoites through a mechanism that involves melanization. We also provide direct evidence that QC-mediated modification of CSP is crucial to avoid immune recognition. However, it remains to be determined whether QC also modifies other sporozoite proteins besides CSP. Combined, our studies reveal a novel mechanism of immune evasion of malaria parasites in the mosquito, including a major role of CSP. This surface protein has multiple essential functions during key infection processes in both the mosquito and vertebrate host (*13, 14, 40*) and is a principal target for various malaria vaccination approaches (*13, 14, 41*). Human glutaminyl cyclases post-translationally modify multiple proteins involved in different processes, including evasion of immune recognition (*7, 8*). Our findings indicate that QC-mediated post-translational protein modification is an ancient immune evasion strategy, shared by evolutionarily distant eukaryotes.

## Materials and Methods

### Experimental animals (ethics statement): Leiden, LUMC (The Netherlands)

Female OF1 mice (6-7 weeks, Charles River Laboratories, NL) were used. All animal experiments were granted with a license by Competent Authority after an advise on the ethical evaluation by the Animal Experiments Committee Leiden (AVD1160020171625). All experiments were performed in accordance with the Experiments on Animals Act (Wod, 2014), the applicable legislation in the Netherlands in accordance with the European guidelines (EU directive no. 2010/63/EU). All experiments were executed in a licensed establishment for the use of experimental animals (LUMC). Mice were housed in individually ventilated cages furnished with autoclaved aspen woodchip, fun tunnel, wood chew block and Nestlets at 21 ± 2°C under a 12:12 hour (h) light-dark cycle at a relative humidity of 55 ± 10%.

Mosquitoes from a colony of *Anopheles stephensi* (line Nijmegen SDA500) were used. Larval stages were reared in water trays at a temperature of 28 ± 1°C and a relative humidity of 80%. Adult females were transferred to incubators with a temperature of 28 ± 0.2°C and a relative humidity of 80%. For the transmission experiments, 3- to 5-day-old mosquitoes were used.

### Experimental animals (ethics statement): NIH, US

Female BALB/c mice (6-7 weeks from Charles River Laboratories, USA) were used following approved NIH animal protocol LMVR-5. *An. stephensi* Nijmegen mosquitoes were reared under standard conditions at 27°C and 80% humidity on a 12 h light-dark cycle.

All experiments involving generation of mutant parasite lines and phenotype analyses were performed using highly standardized and approved protocols that have been developed to reduce the number of animals and minimize suffering and distress. In all experiments mice were killed at a parasitemia of 2-5% before malaria-associated symptoms occur. Mice were killed either by cardiac puncture (under isoflurane anesthesia) or CO_2_.

Humane endpoints: The animals/body condition was thoroughly examined daily. Animals will be humanely sacrificed in case the following defined end points are reached: visible pain (abnormal posture and/or movement), abnormal behavior (isolation, abnormal reaction to stimuli, no food and water intake). If distress of the animals is observed by the animal caretakers, this will be reported to the investigators and according to the aforementioned criteria, the animals will be taken out of the experiment and euthanized. In all experiments no mice were euthanized before termination of the experiment and no mice died before meeting criteria for euthanasia.

### Parasites

Parasites of the transgenic reference line 1868cl1 were used as ‘wild type (WT)’ *P. berghei* (*Pb*) ANKA parasite. Parasites of this reference ANKA line express mCherry and luciferase under the constitutive *hsp70* and *eef1a* promotors, respectively (RMgm-1320, www.pberghei.eu) (*42*). The following *Pb* ANKA mutant lines were used: Δ*rom3* (430cl1; mutant RMgm-178, www.pberghei.eu) (*27*), Δ*crmp4* (376cl1; RMgm-585, www.pberghei.eu) (*28*), Δ*trap* (2564cl3; RMgm-4680, www.pberghei.eu) (*29*) and Δ*csp* (3065cl1; RMgm-4681, www.pberghei.eu) (*26*). Parasites of Δ*csp* and Δ*rom3* (*csp*, PBANKA_0403200; *rom3*, PBANKA_0702700) lack signs of sporozoite formation/maturation inside oocysts (*26, 27*). Parasites of Δ*crmp4* (*crmp4*, PBANKA_ 1300800) form sporozoites that are unable to egress from the mature oocysts (*28*)). Parasites of Δ*trap* (*trap*, PBANKA_1349800) form sporozoites inside oocyst that egress from the oocysts but are unable to invade the salivary glands (*29*). In addition, the mutant line *PbΔcsp*-GIMO parasites (https://www.pberghei.eu/index.php?rmgm=4681) was used. In the genome of parasites of this line the *csp* open reading frame (*orf*) has been replaced by the h*dhfr*::y*fcu* selectable marker. Parasites of this line lack sporozoite formation/maturation inside oocysts.

*P. falciparum* parasites NF54 strain (*43*) were used as WT *P*. *falciparum* parasites (WT *Pf)*. The *Pf*NF54 strain was isolated in 1979 from a Dutch malaria patient and was first adapted to continuous in vitro culture system. Parasites from the *Pf*NF54 strain, and its derivative *Pf*3D7, are the most commonly used *P. falciparum* parasites in laboratory studies. The complete genome sequences of *Pf*3D7 and *Pf*NF54 has been published. The parasites of *Pf*NF54 and *Pf*3D7 have been deposited with the Malaria Research and Reference Reagent Resource Center (MR4; MRA-1000 and MRA-102), which was developed by the National Institute of Allergy and Infectious Diseases (NIAID) and is managed by the American Type Culture Collection (ATCC) (BEI Resources; https://www.beiresources.org/About/MR4.aspx). Parasites were cultured in RPMI-1640 culture medium supplemented with L-glutamine and 25mM HEPES (Gibco Life Technologies), 50 mg/L hypoxanthine (Sigma), 0.225% NaHCO_3_ and 10% human serum at a 5% hematocrit under 4% O_2_, 3% CO_2_ and 93% N_2_ gas-conditions at 75 rpm at 37°C in a semi-automated culture system (Infers HT Multitron and Watson Marlow 520U) as previously described (*44*). Fresh human serum and human red blood cells (RBC) were obtained from the Dutch National Blood Bank (Sanquin Amsterdam, the Netherlands; permission granted from donors for the use of blood products for malaria research and microbiology tested for safety). Production of genetically modified parasites and characterization of these parasites throughout their life cycle, including mosquito transmission, was performed under GMO permits IG 17-230_II-k and IG 17-135_III.

### Bioinformatic analyses

Gene and amino acid sequences of glutaminyl cyclase (QC; glutaminyl-peptide cyclotransferase, putative) from *Plasmodium* species were retrieved from PlasmoDB v.46 (www.plasmodb.org). Similarity and identity between QC of *Plasmodium* species was calculated using MATGAT (*45*). Three dimensional structure of *P. falciparum* QC (*Pf*QC; PF3D7_1446900), generated against the resolved QC structures from *C. papaya* (PDB ID: 2IWA) and *Z. mobilis* (PDB ID: 3NOL) using the I-TASSER structure prediction tool (*46, 47*). Pymol (https://pymol.org/2/) was used to visualize 3D protein structure. Secondary structure of *Pf*QC was aligned with the QCs from *C. papaya* (PDB ID: 2IWA) (*11*), *Z. mobilis* (PDB ID: 3NOL) (*9*), *X. campestris* (PDB ID: 3MBR) (*10*) and *M. xanthus* (PDB ID: 3NOK) (*9*).

Raw mass spectrometry data from the proteomics analysis of sporozoites (*48*) were obtained from the PRIDE database and searched against the *P. falciparum* 3D7 database using the Mascot search algorithm (version 2.2.07, Matrix Science) using Proteome Discoverer 2.4. (Thermo Scientific). Semi-trypsin was selected as enzyme specificity and one missed cleavage was allowed. Precursor and fragment mass tolerance were set at 20 parts per million and 0.4 Da, respectively. Carbamidomethyl (Cys) was set as a fixed modification. Oxidation (Met), acetyl (Protein N-term), Gln->pyro-Glu (N-term Gln) and Glu->pyro-Glu (N-term Glu) were set as variable modifications. Peptide-spectrum matches were adjusted to a 1% FDR.

### Enzymatic characterization of Plasmodium QC

A two-step cyclase activity assay was performed using recombinant WT *Pf*QC and a cyclase-dead PfQC, containing two mutated amino acid residues (F107A and Q109A) in the active site. The mutations F103A and Q105A were selected based on the active site mutational studies of QC from X. campestris (*10*). The gene encoding soluble WT *Pf*QC (without the transmembrane domain; aa56-383) was ordered as a gBlock (IDT) and inserted into the in-house bacterial expression vector pCPF101 using NcoI and XhoI restriction sites, yielding an C-terminally His-tagged construct. The cyclase-dead *Pf*QC was generated using site-directed mutagenesis. Both constructs were sequence-verified and were deposited on Addgene (www.addgene.org). For protein expression the constructs were transformed into BL21(DE3) Rosetta2 cells, using kanamycin (50 μg mL-1) and chloramphenicol (37 μg mL-1) as selection markers. Cells were grown at 37°C in Luria-Bertani (LB) Broth until an optical density (OD600) of 1.4 was reached, upon which the cultures were induced using 0.25 mM IPTG and incubated at 20°C overnight. The next day cells were pelleted through centrifugation (20 minutes at 4000 G) and the pellets were frozen until purification. For isolation of *Pf*QC, the pellets were lysed in lysis buffer (50 mM Tris pH8.5, 200 mM NaCl, 20 mM Imidazole) using sonication. The cell debris was removed using centrifugation (40 minutes at 21000 G) and the supernatant was loaded on Ni-charged NTA sepharose beads. After extensive washing with lysis buffer, the protein was eluted using said buffer supplemented with 200 mM imidazole. Elutions were analysed on gel, *Pf*QC-containing fractions were pooled and diluted using buffer A (20 mM HEPES pH 7.5, 50 mM NaCl, 1 mM DTT) before being loaded on a HiTrapQ anion exchange column using the NGC FPLC (Bio-rad). *Pf*QC was eluted using a salt gradient, with buffer B (20 mM HEPES pH 7.5, 1 M NaCl, 1 mM DTT). *Pf*QC containing fractions were pooled, concentrated and flash frozen for storage at −80°C. Recombinant *Pf*QC activities were determined using SensoLyte green glutaminyl cyclase activity assay kit (Anaspec). Briefly, enzyme activity is measured in a two-step homogeneous procedure using a green fluorescence substrate. During the first step, the substrate is incubated with glutaminyl cyclase or enzyme-containing samples and converted into the pyroglutamate (pGlu) form. Secondly, glutaminyl cyclase developer is added to remove pGlu residue and generate the green fluorophore. Purified *Pf*QC (0.5 mg/ml, 5ul/well) was mixed with fluorogenic substrate (1:100, 5 ul/well) in 384-well black opaque plate and allowed to react for 30 min at 37ᵒC. Developer (pyroglutamyl aminopeptidase) was added to remove pGlu residue and generate the green fluorophore (1:100, 5ul/well) and allowed to react for 30 min at 37ᵒC. Fluorescent signal was read at Ex/Em= 490/520nm. Produced fluorescence is proportional to the enzyme activity. Recombinant human QPCT was used a positive control.

### Generation and genotyping of different P. berghei QC-mutants

To generate QC-null mutants, the *Pbqc* gene (PBANKA_1310700) was deleted by standard methods of transfection (*49*) using the NotI linearized gene-deletion plasmid *Pb*GEM-342996, obtained from PlasmoGEM (Wellcome Trust Sanger Institute, UK; http://plasmogem.sanger.ac.uk/) (*50, 51*). This construct contains the positive-negative h*dhfr*::y*fcu* selectable marker (SM) cassette. Transfection of parasites (line 1868cl1) in two independent experiments, followed by pyrimethamine selection and subsequent cloning of the parasites (*49*) resulted in selection of two gene-deletion mutants *PbΔqc*1 (line 2930cl1) and *PbΔqc*2 (line 2931cl1). Correct integration of the constructs and deletion of the *qc* gene were verified by Southern analyses of Pulsed Field Gel (PFG)-separated chromosomes and diagnostic PCR analysis (*49*). PFG-separated chromosomes were hybridized with a mixture of two probes: a probe recognizing the h*dhfr* gene and a control probe recognizing gene PBANKA_0508000 on chromosome 5(*52*). PCR primers for genotyping are listed in **Table S1**.

An additional QC-null mutants, *PbΔqc*-G, was generated using DNA construct pL2335. This plasmid was generated by cloning the 5’ and 3’ regions of *Pbqc* into the restriction sites of HindIII/PstI and KpnI/EcoRI of the standard cloning vector pL0034, which contains the h*dhfr*::y*fcu* SM under the control of the *Pbeef1a* promoter (*53*). The 5’ and 3’ regions targeting sequences of *Pbqc* were amplified from *P. berghei* genomic DNA (gDNA) using primer sets 9641/9642 and 9643/9644 (see **Table S1** for the sequence of all primers). Plasmid pL2335 was linearized with HindIII and EcoRI and transfected in parasites (line 1868cl1) using standard GIMO-transfection procedures (*53*). Transfection of parasites, pyrimethamine selection and subsequent cloning of the parasites (*49*) resulted in selection of gene-deletion mutant *PbΔqc*-GIMO (line 3198cl1) with the *qc* open reading frame (*orf*) replaced by the h*dhfr*::y*fcu* SM. Correct deletion of the *Pbqc orf* was confirmed by diagnostic PCR analysis of genomic DNA (gDNA) and Southern analysis of PFG-separated chromosomes as described (*49*). The primers used for PCR genotyping are listed in **Table S1**.

To generate transgenic *P. berghei* parasites expressing C-terminal cmyc-tagged QC, DNA construct pL2351 was generated. The full length *Pbqc orf* without stop codon was amplified from genomic DNA using primer sets 9233/9234 (see **Table S1** for primer sequences) and used to replace the *Pbmrp2 orf* (PBANKA_144380) using the restriction sites NotI/BamHI of plasmid pL1672 (*54*). This plasmid contains the *Tgdhfr/ts* SM under the control of the *Pbeef1a* promoter. DNA-construct pL2351 was linearized with BsaBI (a unique restriction site present in *Pbqc orf*). Transfection of parasites of line 1868cl1 followed by pyrimethamine selection (*49*) resulted in selection of the *Pbqc::cmyc* tagged line (line 3272). Correct integration of the *cmyc*-tagging construct was confirmed by diagnostic PCR analysis of gDNA and Southern analysis of PFG-separated chromosomes (*49*). PCR primers for genotyping are listed in **Table S1**.

For complementation of *PbΔqc* QC-null parasites with either the *Pfqc* or *Pbqc* gene, the *PbΔqc*-GIMO parasite line (line 3198cl1) was used. For complementation with *Pfqc*, DNA construct pL2326 was generated. The *Pbqc 5’ utr* and *3’ utr* were amplified from *P. berghei* gDNA using primers 9483/9484 and 9485/9486 respectively and introduced sequentially into pBSKS+ plasmid (Stratagene) at XhoI/BamHI and EcoRI/SacII restriction sites respectively, to obtain intermediate plasmid SKK94. The *Pfqc orf* was amplified from WT *P. falciparum* NF54 gDNA using primers 9487/9488 and ligated into SKK94 using BamHI and EcoRI restriction sites, to obtain pL2326. For complementation with *Pbqc*, the *Pbqc orf* along with 1kb flanking sequences at both the 5’ and 3’ sides was amplified from WT *P. berghei* ANKA gDNA using primers 9483/9486 and ligated into pJET1.2 blunt cloning vector (ThermoScientific, Cat# K1231) to obtain pL2327 (see **Table S1** for primer sequences). The DNA-constructs pL2326 and pL2327 were sequenced to confirm absence of mutations and both plasmids were linearized with XhoI and SacII before transfection. Transfection of *PbΔqc*-GIMO parasites with both constructs, followed by applying negative selection with 5-fluorocytosine (5-FC) (*49, 52*) and subsequent cloning of the parasites resulted in selection of complemented parasites where the h*dhfr*::y*fcu* SM in the *qc* locus of *PbΔqc*-GIMO is replaced by the *orf* of either *Pfqc* (line 3213cl1; *Pfqc*(c)) or *Pbqc* (line 3216cl1; *Pbqc*(c)). Correct integration of the constructs into the genome of the complemented parasites was analyzed by diagnostic PCR analysis on gDNA and Southern analysis of PFG-separated chromosomes as described (*49*). PCR primers for genotyping are listed in **Table S1**.

To generate *Pbqc*^*CD*^ parasites, expressing the catalytic dead enzyme QC^*CD*^, DNA construct pL2348 was generated. Partial fragments of *Pbqc orf* were amplified from *P. berghei* genomic DNA using primers 9483/9515 and 9516/9486. The primers 9515 and 9516 cover an overlap of a 29 bp region, with the mutations F103A and Q105A that were selected based on studies of active site mutation of QC from *X. campestris* (*10*). These mutations were introduced by site-directed mutagenesis, following the overlap extension PCR. Subsequently, the full length *qc*^*CD*^, obtained by overlap extension of the two partial fragments using primers 9483/9486, was cloned in pJET 1.2 blunt cloning vector (K1232, Thermo Scientific). This resulted in construct pL2348 that was sequenced to confirm the presence of the two point mutations and absence of undesired mutations. Transfection of *PbΔqc*-GIMO parasites with the XhoI and SacII linearized construct pL2348, followed by applying negative selection with 5-fluorocytosine (5-FC) (*49, 52*), resulted in selection of *Pbqc*^*CD*^ parasites (line 3276) where the h*dhfr*::y*fcu* SM in the *qc* locus of the *PbΔqc-*GIMO line is replaced by the mutated *Pbqc orf.* Correct integration of the constructs into the genome of *Pbqc*^*CD*^ parasites was analyzed by diagnostic PCR analysis on gDNA and Southern analysis of PFG-separated chromosomes as described (*49*). In addition, a fragment of 624bp was amplified using primers 8991/8992 covering the F103A/Q105A mutated region from *Pbqc*^*CD*^ gDNA, cloned in pJET 1.2 blunt end cloning vector and sequenced using pJET forward and reverse primers to confirm the mutations. PCR primers for genotyping are listed in **Table S1**.

To generate the four *P. berghei* ‘double knockout’ mutants, the h*dhfr*::y*fcu* SM was first recycled from *PbΔqc*1 parasites by negative selection with 5-Fluorocytosine (5-FC) as described (*52, 55*). This treatment selects for parasites that have undergone homologous recombination between the two 3’-UTR sequences of *Pbdhfr*, present in the integrated construct *Pb*GEM-342996 (pL2243) that flank the h*dhfr*::y*fcu* SM cassette and thereby removing the SM (*52, 56*). Selection and cloning of the parasites resulted in the SM-free ‘single knockout’ mutant *PbΔqc*1(-sm) (line 3172cl1(2^nd^)). This line was used to delete the following genes in independent transfection experiments: *csp* (PBANKA_040320)*, crmp4* (PBANKA_1300800), *ecp1* (PBANKA_0304700), *rom3* (PBANKA_0702700), and *trap* (PBANKA_1349800). To delete the *csp*, *crmp4*, *ecp1*, *rom3* and *trap* genes, the gene-deletion construct pL2153 (RMgm-4681; www.pberghei.eu) and the following PlasmoGEM constructs (http://plasmogem.sanger.ac.uk/ (*57*)) *Pb*GEM-342964 (pL2379), *Pb*GEM-269819 (pL2367), *Pb*GEM-342020 (pL2368) and *Pb*GEM-338811 (pL2378) were used, respectively. All four constructs contain the h*dhfr::yfcu* SM. The constructs pL2153 and pL2111 were linearized using the restriction enzymes ApaI/NotI and pL2367 and pL2368 were linearized with NotI before transfection. Transfection of parasites of line *PbΔqc*1(-sm) with linearized constructs, positive selection of transfected parasites with pyrimethamine and cloning of selected parasites were performed as described (*49*). This resulted in the following four lines: *Pb*Δ*qc*Δ*csp* (line 3322cl1), *Pb*Δ*qc*Δ*crmp4* (line 3367cl4), *Pb*Δ*qc*Δ*ecp1* (line 3324cl2), *Pb*Δ*qc*Δ*rom3* (line 3326cl1) and *PbΔqcΔtrap* (line 3365cl1) that were used for genotype and phenotype analyses. Correct integration of the constructs into the genome of *PbΔqc*1(-sm) parasites was analyzed by Southern analysis of PFG-separated chromosomes as described (*49*).

To generate two independent *P. berghei* mutants (*Pbcs*^mut^) that express a mutated CS with the glutamine in region I replaced with an alanine (Q92A), DNA construct pL2360 was generated. The *cs 5’utr* and part of the *orf* (until the mutation in region I) was amplified as fragment-1 using the primers 9771 and 9772. The remainder of the *cs orf* and *3’utr* were amplified as fragment-2 using the primers 9773 and 9774. A unique restriction site EcoO109I was introduced in the mutated region using primers 9772 and 9773. Fragment-1 was digested with PstI/EcoO109I and fragment-2 with EcoO109I/SacI and subsequently introduced into the Pst/SacI sites of pUC19 vector by three fragment ligations. This resulted in construct pL2360 that was sequenced to confirm the presence of the point mutation and absence of undesired mutations. The plasmid pL2360 was linearized with XhoI and transfected into *PbΔcsp*-GIMO parasites (https://www.pberghei.eu/index.php?rmgm=4681) in two independent experiments, followed by applying negative selection with 5-fluorocytosine (5-FC) (*49, 52*) and cloning of selected parasites (*49*). This resulted in selection of the *Pbcs*^mut^ parasites (line 3299cl1, *Pbcs*^mut1^ and 3300cl, *Pbcs*^mut2^) where the h*dhfr*::y*fcu* SM in the *csp* locus of the *PbΔcsp-* GIMO line is replaced by the Q92A mutated *Pbcsp orf.* Correct integration of the construct into the genome of the *Pbcs*^mut^ parasites was analyzed by diagnostic PCR analysis on gDNA and Southern analysis of PFG-separated chromosomes as described (*49*). PCR primers for genotyping are listed in **Table S1**.

### Generation and genotyping of P. falciparum QC-null mutants

The *Pfqc* (PF3D7_1446900) gene was deleted in WT *Pf* parasites by standard methods of CRISPR/Cas9 transfection (*58*) using two different sgRNA-expressing plasmids, containing the Cas9 expression cassette, guide-RNA expression cassettes and h*dhfr* SM cassette, in combination with donor DNA plasmid pLf0149 that contains a blasticidin (BSD) SM cassette. The two different sgRNA-expressing plasmids were generated as follows: pLf0070 (*58*) was digested with BbsI and sgRNA062 and sgRNA063 were cloned using primers 9251/9252 and 9253/9254, respectively, resulting in the plasmids pLf0150 and pLf0151 (see **Table S1** for primer sequences). The donor DNA plasmid pLf0149 was generated to replace the *Pfqc orf* with a BSD SM cassette flanked by *flippase recognition target* (*frt*) sequences (these sequences allow to recycle the SM cassette (*59*)). The 5’ and 3’ targeting regions of *Pfqc* were amplified from WT *Pf* genomic DNA using the primers 9211/9212 and 9250/9214 (**Table S1**). Fragments were digested with HindIII/ApaI and NheI/BamHI respectively and ligated into plasmid pLf0103 to obtain pLf0149. Plasmid pLf0103 was generated as follows: first, the *BSD orf*, flanked by 34bp *frt* sequences and multiple cloning sites on either side, was synthesized as a gene block by Integrated DNA Technologies (IDT, Belgium), and inserted into plasmid pL0034 (*53*) to get pFB1. Next, the *Pfhsp70* (PF3D7_0818900) promoter was amplified from WT *Pf* genomic DNA using primers 8801/8802 and ligated into pFB1 using restriction sites XhoI and KpnI to get pFB2. Subsequently, *Pfhrp2* (PF3D7_0831800) *3’utr* was amplified from WT *Pf* genomic DNA using primers 8803/8804 (see **Table S1** for primer sequences) and ligated into pFB2 using the restriction sites AvrII and NotI to get pLf0103. Transfection of WT *Pf* parasites, selection and cloning of *PfΔqc* parasites in two independent experiments was performed as described (*44, 60*), resulting in the lines *PfΔqc*1 (exp. 189) and (*PfΔqc*2; exp. 250). In these experiments we used two different stabilates of WT *Pf* parasites (received from Radboud University Medical Center, Nijmegen, The Netherlands), stabilate 9002 (NF54 54/329 4514) for *PfΔqc*1 (exp. 189) and stabilate 9001 (*PfΔqc*2; exp. 250). For genotyping *PfΔqc*1 and *PfΔqc*2 parasites, diagnostic PCR and Southern analysis of digested gDNA was performed as described (*44*) (see **Table S1** for primer sequences). Southern blot analysis was performed with gDNA digested with ClaI, XhoI and BamHI (4 hrs at 37°C) in order to confirm the deletion of *Pfqc*. Digested DNA was hybridized with a probe targeting the *Pfqc orf*, amplified from WT *Pf* genomic DNA by PCR using primers 9597/9598 **(Table S1)** and the *Pftrap* probe, used as DNA loading control, was amplified using primers 9193/9274.

### Phenotype analysis of P. berghei QC mutants

The *in vivo* multiplication rate of asexual blood stages was determined during the cloning procedure of the different QC mutants (*61*). *In vitro* cultivation of ookinetes, using gametocyte-enriched infected blood was performed as described (*62*).

For mosquito transmission experiments, female *An. stephensi* mosquitoes were fed on infected mice with a 2-5% parasitemia. Development and production of oocysts and sporozoites in infected mosquitoes were analyzed by light- and fluorescence-microscopy as described (*63*). Salivary gland sporozoites (sg-sporozoites) were analyzed and counted, obtained from isolated salivary glands at day 21–23 p.i. Sg-sporozoites were isolated after manual dissection of mosquito salivary glands and counted in a Bürker cell counter using phase-contrast microscopy as described (*64*).

Sporozoite infectivity *in vivo* was determined by determination of parasite liver loads in OF1 mice by *in vivo* imaging and determination of prepatent period in mice injected intravenously with 1 × 10^4^ isolated sg-sporozoites as described (*52, 64, 65*). Blood stage infections were monitored in Giemsa stained tail blood smears. The prepatent period is defined as the day on which the parasitemia reaches 0.5-2%.

Sporozoite infectivity *in vitro* was determined in cultures of Huh7 cells (JCRB0403, human hepatoma cell line obtained from JCRB Cell Bank, Japan) as described (*64, 65*). In brief, a total of 5 × 10^4^ sg-sporozoites were added to monolayers of Huh7 cells in complete RPMI-1640 medium. At 24 and 48 h post infection, samples were stained with Hoechst 33348 (10μM) and live imaging of mCherry-expressing liver stages was performed using a DM RA Leica inverted fluorescent microscope (40× magnification). Image analysis was done with Leica LAS X software.

Motility of sg-sporozoites was analysed by *in vitro* microscopic live imaging as described (*66*). In brief, sg-sporozoites were isolated in RPMI-1640 culture medium supplemented with 10% fetal bovine serum (FBS) and diluted to 20×10^6^ sporozoites/ml. For imaging, 10 μl of the sporozoite solution was placed on the glass portion of a confocal dish (ø14 mm; MatTek Corporation), covered with a 12 mm cover slip and imaged within 45 min. The images were analysed with the quantitative analysis software SMOOT_*In vitro*_, an in-house developed graphical user interface (GUI), written in the MATLAB programming environment (version r2017b, The MathWorks Inc.) (*66*). Images of sg-sporozoites were taken on a Leica TCS (true confocal scanning) SP5 or SP8X WLL (white light laser) microscope (Leica Microsystems, Wetzlar). Sg-sporozoites expressed mCherry, which was excited at 587 nm and the emission was collected between 600 and 650 nm. The movies were recorded with a rate of 35 frames per minute, 400 frames per movie, three movies per condition. For imaging a 40× objective (Leica HCX PL APO CS 40×/1.25–0.75 numerical aperture OIL) was used. The movies were recorded at room temperature (RT) using the Leica software (LAS X version 1.1.0.12420; Leica Microsystems, Wetzlar). Maximum projections of the microscopy movies were generated using Fiji software (*67*) and were further processed using SMOOT_*In vitro*_ (*66*). Via SMOOT_*In vitro*_ analysis sg-sporozoites could be segmented per movie frame, based on fluorescence signal intensity, size and crescent shape by applying a binary threshold, a median filter, skeleton adjustment and spot removal. The movement pattern of the sg-sporozoites was classified per segment as floating, stationary or circling and the velocity of the circling sg-spz was determined. On average ~ 100 sg-spz were analysed per condition.

QC expression in *Pbqc::cmyc* parasites was analysed in blood stages, gametocytes, *in vitro* cultured ookinetes (*68*) and sg-sporozoites by Western blotting and indirect immunofluorescence analysis (IFA) as described (*69, 70*). Circumsporozoite protein (CSP) expression in WT and *Pbcsp*^mut^ parasites was analysed by Western blotting.

For IFA, the samples were fixed with 4% paraformaldehyde in phosphate-buffered saline (PBS) for 15 min at room temperature (RT), permeabilized with 0.5% Triton X-100 in PBS for 15 min at RT and then washed twice with PBS and blocked with 10% FBS in PBS for 30-60 min at RT. Primary incubation was performed with rabbit anti-cmyc (Sigma, # C3956) antibody (1:200 dilution in blocking solution) for 1 h at RT. After incubation, slides were washed three times with PBS, followed by incubation with Alexa Fluor 488 conjugated goat anti-rabbit secondary antibody (Invitrogen, #A11008) diluted 1:500 in blocking medium for 45 min at RT. After washing three times with PBS, nuclei were stained with Hoechst-33342 (10 μM) for 30 min at room temperature and washed twice with PBS. The slides were mounted with cover slips (VECTASHIELD PLUS Antifade Mounting Medium, Vector laboratories, #H-1900) and sealed with nail polish. Imaging was performed using a Leica fluorescence MDR microscope.

For western blotting, the parasite proteins were separated by electrophoresis on a 12% SDS-PAGE gel and transferred on to Hybond ECL nitrocellulose membrane (Amersham Biosciences) for 2 h at 200 mAh. Membranes were blocked for non-specific binding with 3% skim milk (Elk, Campina, The Netherlands) in PBS with 0.1% Tween 20 (PBST) overnight at 4°C. Blots were hybridized with rabbit anti-cmyc (Sigma, # C3956) polyclonal antibody (1:1000 dilution), mouse anti-Bip polyclonal antibody used as an internal control (*71*) (1:1000 dilution, kind gift from Volker Heussler; University of Bern) or mouse monoclonal anti-CSP (3D11) antibody (*72*) and the membranes were washed with PBS and incubated with horseradish peroxidase (HRP)-conjugated donkey anti-rabbit (GE Healthcare, #NA934V) and/or sheep anti-mouse (GE Healthcare, #NA931V) IgG secondary antibody for 1 h at RT and developed in Amersham ECL Western Blotting Detection Kit according to the manufacturer’s instructions (GE Healthcare).

Melanization of oocysts and sporozoites was analyzed between day 8 and 23 post infection (p.i.) of *An. stephensi* mosquitoes by light-, fluorescence- and transmission electron microscopy (TEM). For light- and fluorescence microscopy, infected *An. stephensi* mosquito midguts or sporozoites were manually dissected using Leica MZ16 FA stereo-fluorescent microscope (as described above). The midguts were imaged with Leica MZ camera at 10X magnification using Leica LAS X software. The melanized and normal oocysts were observed under Leica DM2500 light microscope and documented at 100X using Leica DC500 digital camera using Leica LAS X software.

Melanized oocysts were defined as dark (black/brown) colored oocyst with the dark color covering an area of 50-100% of the oocyst. Melanized hemocoel sporozoites were defined as large, non-motile, dark-colored forms with >90% of their surface was covered with a dark-colored layer. Melanized sporozoites in the hemocoel were analyzed by dissecting the abdomen on the lateral sides.

For Transmission Electron Microscopy (TEM) analyses midguts were manually dissected at day 14 and fixed with 2% paraformaldehyde and 2.5% glutaraldehyde (TAAB Laboratories Equipment Ltd.) in 0.1 M phosphate buffer (pH 7.4) overnight at 4°C, followed by post-fixation with 2% OsO_4_ in the same buffer. Thereafter, the fixed midguts were dehydrated with a graded series of ethanol (50, 70, 90, 95, 100%) and embedded in Epok 812 (Oken Shoji). Ultrathin sections (80 nm thickness) were cut with an ultramicrotome UC6 (Leica) and transferred to 150-mesh copper grids, and stained with uranyl acetate and lead citrate and were then examined using an HT7700 transmission electron microscope (Hitachi) (*73*).

For line-scanning and mapping analysis by Scanning Electron Microscopy with Energy Dispersive X-Ray Analysis (SEM-EDX) of melanized and non-melanized oocysts and sporozoites, midguts of *PbΔqc*1 infected mosquitoes (day 14) were fixed with 2% paraformaldehyde and 2.5% glutaraldehyde in 0.1 M phosphate buffer (pH 7.4). After dehydration through an ascending series of acetone, the acetone was replaced with liquid CO2 in the chamber of a critical point dryer (CPD 300, Leica), and the specimens were dried by critical point drying. The specimens were coated with osmium by using an osmium plasma coater (Neoc-Pro/P, Meiwafosis). Observation of the selected points by using a SEM (Regulus8220, Hitachi High-Tech), the EDX analysis was performed with the installed EDX system (EMAX Evolution EX-370_X-max50, Oxford Instruments). The spectrum data were acquired at an acceleration voltage of 20kV.

### Phenotype analysis of P. falciparum mutants

The growth rate of asexual blood stages of *P. falciparum* parasites was determined in 10 ml cultures maintained in the semi-automated culture system as described (*44*). Gametocyte production and exflagellation were quantified in gametocyte cultures as described (*58*).

For mosquito transmission experiments, female *An. stephensi* mosquitoes were fed on gametocyte cultures using the standard membrane feeding assay (SMFA) (*58, 74*). Oocyst and sporozoite production were monitored in infected mosquitoes as described (*58, 63*). Oocysts were analyzed on day 11, 15 and 21 p.i. and the percentage of melanized oocysts was determined by analyzing manually dissected midguts using a Leica MZ16 FA stereo-fluorescent microscope. The midguts were imaged with a Leica MZ camera at 10X magnification using Leica LAS X software. Individual melanized and WT-like oocysts were observed under a Leica DM2500 light microscope and documented with at 100X using Leica DC500 digital camera using Leica LAS X software. Sg-sporozoite numbers were analyzed in infected mosquitoes at day 18-21 p.i.. For counting sporozoites, salivary glands from 30-60 mosquitoes were dissected and homogenized using a grinder in 100 μl of RPMI-1640 medium (pH 7.2) and sporozoites were analyzed in a Bürker cell counter using phase-contrast microscopy (*58*).

### Disruption of the mosquito immune system

Mosquito hemocyte depletion was carried out as previously described (*20*). Briefly, *An. stephensi* mosquitoes received an injection of 200,000 1 μm diameter polystyrene beads (FluoSpheres, Life Technologies) in 138 nl in water using a nanoject injector (Drumond Scientific Co.). Polystyrene beads were washed extensively with water before injecting mosquitoes in the side of the thorax 12 days p.i. Water injection was used as a control.

The abundance of sporozoites was determined by measuring luciferase activity in the thorax of mosquitoes infected with *P. berghei* WT and mutant parasites that express luciferase constitutively.

Mosquito thoraces were collected 23 days p.i. in individual tubes, frozen in dry ice immediately and stored at −30°C until processing. The Luciferase Assay System kit (Promega) was used to measure the luciferase activity in sporozoite samples following the manufacturer’s instructions. Briefly, mosquito thoraces were individually homogenized in 100μl of 1X passive lysis buffer using a pellet pestle. Positive and negative control stocks were prepared, positive controls were made by homogenizing individual abdomens from WT-infected mosquitoes, while negative controls were made by homogenizing individual uninfected mosquitoes. The homogenized thoraces were incubated at RT for 10 min and then centrifuged at 13,000 rpm for 1 min, 20μl of supernatant was pipetted in duplicate into a 96 well, black, flat-bottomed, opaque plate. The luminescence assay was run in a Cytation5 apparatus set for 25°C, injection of 50μl Luciferase Assay Reagent (LAR) and reading of 1 sec/well. The 95% confidence interval of the 4 negative controls were subtracted as background from measurements done in the same plate.

dsRNA-mediated gene silencing of LRIM1 (ASTE000814) and CLIPA8 (ASTE009395) orthologues in *An. stephensi* was done as previously described (*75*). Briefly, mosquitoes were injected with 69 nL of a 3 μg/μL dsRNA solution 12 days p.i. with *P. berghei* WT and mutant parasites. The control dsRNA (LacZ) and *An. stephensi LRIM1 and CLIPA8 dsRNA* were produced as previously described (*75, 76*). Briefly, dsRNA was made using the MEGAscript RNAi Kit (Ambion), using DNA templates obtained by nested PCR using cDNA from whole female mosquitoes. LRIM1 external PCR was performed using the primers : LRIM1_Ast814ExF and LRIM1_Ast814ExR (see **Table S1** for primer sequences). PCR conditions were: 94°C for 3 min, 25 cycles of 94 °C for 30 s, 55 °C for 1 min, and72 °C for 1 min with final extension 72 °C for 5 min. Nested PCR was carried out with primers LRIM1_Ast814InF and LRIM1_Ast814InR containing a T7 promoter (see **Table S1** for primer sequences). PCR conditions were 94 °C for 3 min, 30 cycles of 94 °C for 30 s, 55 °C for 30 s, and 72 °C for 1 min with final extension 72 °C for 5 min, using 1 μL of the external PCR product. CLIPA8 external PCR was performed using the primers F_dsRNA_Asteph009395b and R_dsRNA_Asteph009395b and nested PCR was carried out with primers F_T7nest_dsRNA_As009395b and R_T7nest_dsRNA_As009395b containing a T7 promoter (see **Table S1** for primer sequences).

Silencing was assessed in whole sugar fed mosquitoes by quantitative real-time PCR (qPCR) using the S7 ribosomal protein gene as internal reference. The primers Aste_LRIM1_F and Aste_LRIM1_R were used for qPCR of LRIM1, F_qPCR_As009395b and R_qPCR_As009395b for CLIPA8 (see **Table S1** for primer sequences). Ribosomal protein S7 primers S7F and S7R were used for the internal reference (see **Table S1** for primer sequences). qPCRs were performed under standard conditions using 0.5 μM of each primer with an initial denaturation step of 15 min at 95 °C and then 45 cycles of 10 s at 94 °C, 20 s at 55 °C, and 30 s at 72 °C, with a final extension of 5 min at 72 °C. The silencing efficiency in dsRNA injected mosquitoes was 85%, relative to dsLacZ-injected controls.

### Statistical analysis

Data were analysed using GraphPad Prism software v7 or above (GraphPad Software, Inc). Statistical analyses were performed using the non-parametric unpaired t-test following Mann-Whitney test, t-test for gliding motility assay and Chi-square test (one-sided) for analyzing CLIP8 dsRNA-mediated inhibition experiments.

## List of Supplementary Figures and Tables

**Fig S1**. Amino acid sequence comparison of QCs of different *Plasmodium* species

**Fig S2**. Amino acid sequence comparison of glutaminyl cyclase (QC) of different organisms

**Fig S3**. Published genome-wide proteomic and transcriptomic data of *Plasmodium* QC expression

**Fig S4**. Generation, genotyping and characterization of transgenic *P. berghei* parasites expressing C-terminal cmyc-tagged QC

**Fig S5**. Generation, genotyping and characterization of three *P. berghei* QC-null mutants (*PbΔqc*)

**Fig. S6**. Light- and fluorescence microscope images of melanized of oocysts and sporozoites of *P. berghei* QC-null mutants

**Fig. S7**. Transmission electron microscopy images (TEM) of melanized of wild type (WT) and QC-null oocyst (day 14)

**Fig S8**. Gliding motility of QC-null salivary gland (sg) sporozoites

**Fig S9**. Infectivity of salivary gland (sg) sporozoites of different *Pbqc* mutants

**Fig S10**. Generation, genotyping and characterization QC-null mutants complemented with *P. berghei* QC (*Pbqc*(c)) or *P. falciparum* QC (*Pfqc*(c))

**Fig S11**. Generation, genotyping and characterization of the *P. berghei* cyclase-dead mutant (*qc*^*CD*^)

**Fig S12**. Generation and genotyping of *P. falciparum* QC-null mutants (*PfΔqc*)

**Fig S13**. Absence of melanized oocysts or sporozoites at day 23 in mosquitoes infected with four *P. berghei* mutants with aberrant oocyst and/or sporozoite formation

**Fig S14**. Generation, genotyping and characterization of the *P. berghei* ‘double knockout’ mutants lacking expression of *qc* and genes involved in sporozoite formation, sporozoite egress or invasion of salivary glands.

**Fig. S15**. Absence of melanized oocysts and sporozoites at day 21 in mosquitoes infected with QC-null mutants lacking either expression of CSP, CRMP4 or ROM3 and melanized oocyst and sporozoites of a QC-null mutant lacking expression of TRAP

**Fig S16**: Analysis of published mass spec data of *P. falciparum* circumsporozoite protein (CSP) for the presence of N-terminal pGlu

**Fig S17**: Generation and genotyping of *P. berghei* mutant expressing CSP with an amino acid replacement (mutation) in region I (*csp*^*mut*^)

**Table S1**: List of primers used

**Table S2**. Gametocyte, ookinete and oocyst production of wild type *P. berghei* parasites (WT) and parasites of *Δqc* lines (Δ*qc* 1, 2, G, 1(-sm)), complemented lines (*Pfqc*(c); *Pbqc*(c)); cmyc-tagged line (*Pbqc::cmyc*) and catalytically-dead mutant line (*Pbqc*^*CD*^)

**Table S3**: p-values

## Acknowledgements

We thank Els Baalbergen (LUMC, Leiden, The Netherlands) for insectary support; Shinya Miyazaki, Catherin Marin-Mogollon and Yuki Miyazaki (LUMC, Leiden, The Netherlands) for *P. falciparum* QC studies; Hans van Leeuwen (TNO, Rijswijk, The Netherlands) for discussion on *in silico* modelling studies; Kevin Lee, Yonas Gebremicale and André Laughinghouse (NIAID, USA) for insectary support; Ton Schumacher and Meike Logtenberg (NKI, Amsterdam, The Netherlands) for QC-activity discussions; George Janssen and Peter van Veelen (CPM, LUMC, Leiden, The Netherlands) for mass-spec data discussions. We acknowledge Shahid Khan^§^ for his contribution in designing the first set of experiments. (^§^deceased).

## Funding

C.B. and A.M. were supported by the Intramural Research Program of the Division of Intramural Research, NIAID/NIH Z01AI000947. F.A.S. was supported by a Leiden University Medical Center fellowship. T.A., S.K., H.H. and T.A. were supported in part by grants for Research on Emerging and Re-emerging Infectious Diseases from Japan Agency for Medical Research and Development (AMED) (JP21fk0108096j0303, JP21fk0108139j3002 and JP21fk0108138j0602) and from the Japan Society for the Promotion of Science (JSPS) (KAKENHI JP21K06999).

## Author contribution

Conceptualization: S.K.K., A.M., C.B., F.A.S., and C.J.J.

Methodology: S.K.K., A.M., S.K., H.H., H.K., T.An., R.Q.K., M.A.C., C.B., F.A.S., and C.J.J.

Investigation: S.K.K., A.M., T.Ar., J.F.A.G., J.R., S.C., H.J.K., S.B., C.K., R.W., N.R., A.F.E.H., R.Q.K., S.K., H.H., H.K., T.An., P.J.H., B.M.F., C.B., F.A.S., and C.J.J.

Visualization: S.K.K., A.M., T.Ar., J.F.A.G., J.R., S.C., H.J.K., S.B., C.K., R.W., N.R., A.FE.H., R.Q.K., S.K., H.H., H.K., T.An., P.J.H., B.M.F., C.B., F.A.S., and C.J.J.

Funding acquisition: T.An., C.B., F.A.S., and C.J.J.

Project administration: C.B., F.A.S., and C.J.J.

Supervision: T.An., C.B., F.A.S., and C.J.J.

Writing – original draft: S.K.K., A.M., T.Ar., T.An., P.J.H., C.B., F.A.S., and C.J.J.

Writing – review & editing: S.K.K., A.M., T.Ar., T.An., C.B., F.A.S., and C.J.J.

## Competing interests

Authors declare that they have no competing interests.

## Data availability

The authors declare that the data supporting the findings of this study are available in the main text or supplementary materials. *Plasmodium* mutant lines generated in this study are available from Dr. C.J. Janse (Leiden University Medical Center, LUMC, Leiden The Netherlands) after completing a materials transfer agreement (MTA).

## Notes

### Competing Interest Statement

The authors have declared no competing interest.

