## Supplementary figures and tables for "Malaria parasite evades mosquito immunity by glutaminyl cyclase mediated protein modification"

**This PDF file includes:**

Figs. S1 to S17
Tables S1 to S3

### Fig. S1

## A

| | PF3D7 | PBANKA | PVP01 | PKNH | PcyM | PGAL8A | PY17X | PCHAS |
|---|---|---|---|---|---|---|---|---|
| **PF3D7** | | 50.1 | 53.0 | 53.0 | 50.4 | 54.3 | 50.3 | 49.7 |
| **PBANKA** | 75.7 | | 54.5 | 54.7 | 42.5 | 56.4 | 91.6 | 83.3 |
| **PVP01** | 74.2 | 77.3 | | 80.7 | 45.5 | 87.5 | 55.1 | 54.3 |
| **PKNH** | 71.5 | 73.5 | 88.5 | | 48.2 | 80.7 | 55.0 | 54.5 |
| **PcyM** | 70.2 | 64.6 | 66.4 | 69.6 | | 46.2 | 43.5 | 43.4 |
| **PGAL8A** | 74.0 | 78.1 | 96.6 | 88.3 | 66.4 | | 56.4 | 55.1 |
| **PY17X** | 74.7 | 98.2 | 77.3 | 73.6 | 63.9 | 78.1 | | 82.3 |
| **PCHAS** | 73.1 | 94.2 | 77.3 | 73.7 | 63.6 | 76.6 | 93.7 | |

## B

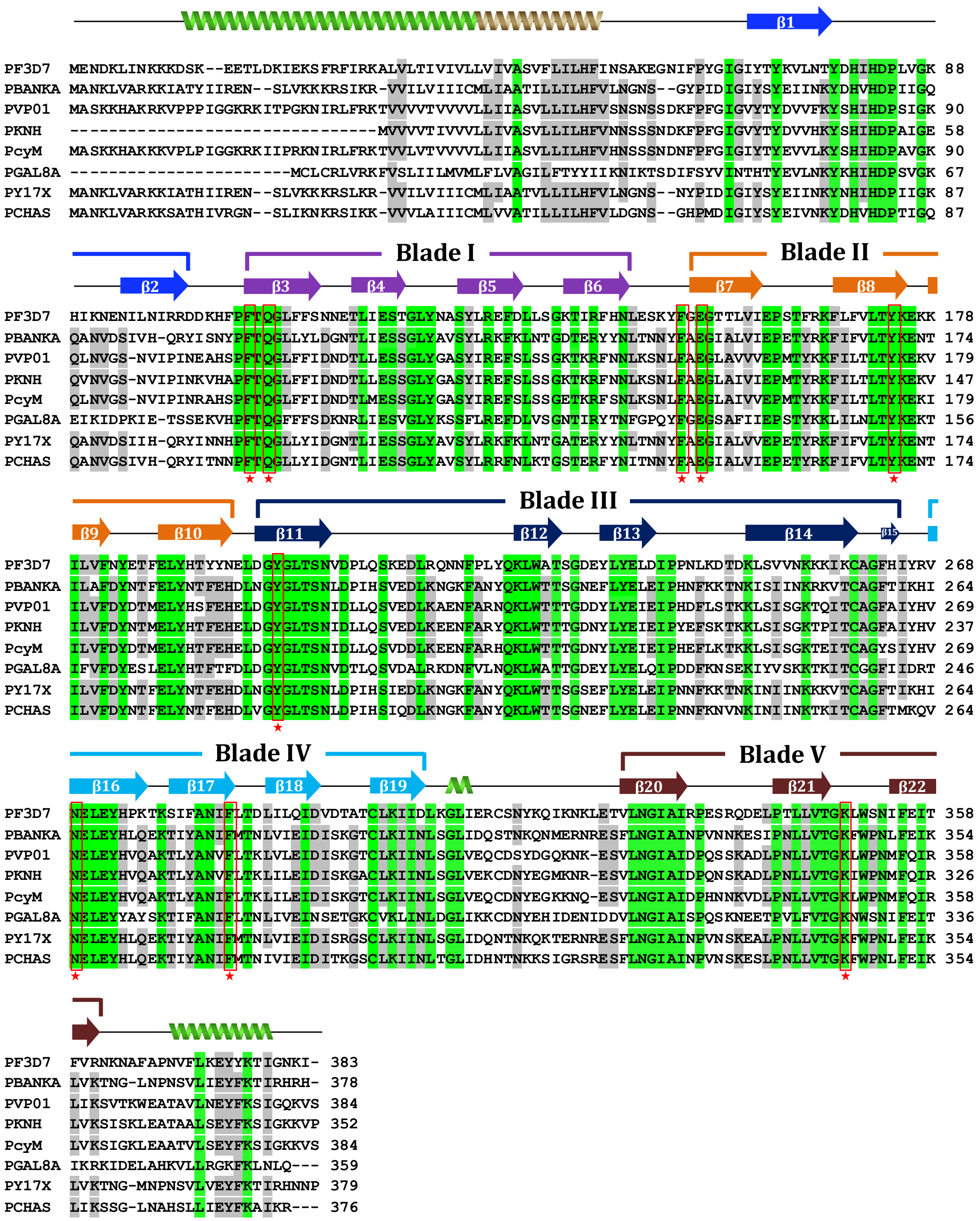

**Fig. S1. Amino acid sequence comparison of QCs of different *Plasmodium* species**

**A**. Sequence identity (%, purple) and similarity (%, blue) between QCs from *P. falciparum* (PF3D7, PF3D7_1446900), *P. berghei* (PBANKA, PBANKA_1310700), *P. vivax* (PVP01, PVP01_1260100), *P. knowlesi* (PKNH, PKNH_1235300), *P. cynomolgi* (PcyM, PcyM_1268900), *P. gallinaceum* (pGAL8A, PGAL8A_00206100), *P. yoelii* (PY17X, PY17X_1314500) and *P. chabaudi* (PCHAS, PCHAS_1314000).

**B.** Sequence alignment of QCs from the different *Plasmodium* species. Secondary and tertiary structural elements (2 α-helices, 22 β-sheets and 5 blades) based on the predicted *Pf*QC structure are shown above the alignment (helices – coils and β-sheets – arrows). The α-helix in the predicted *P. falciparum* transmembrane domain (amino acids 32-54) is indicated in brown and conserved residues in green. Conserved active site residues of all QCs, identified by aligning with the resolved QCs from *C. papaya* and different bacterial species (*Z. mobilis, X. campestris and M. xanthus*), are marked as magenta stars.

### Fig. S2

**A**

| | *PfQC* | *MxQC* | *ZmQC* | *XcQC* | *CpQC* |
|---|---|---|---|---|---|
| ***PfQC*** | | 21.1 | 26.8 | 23.0 | 25.2 |
| ***MxQC*** | 35.0 | | 33.0 | 35.7 | 32.5 |
| ***ZmQC*** | 44.4 | 53.0 | | 48.1 | 34.4 |
| ***XcQC*** | 38.1 | 55.4 | 65.7 | | 37.4 |
| ***CpQC*** | 43.6 | 50.0 | 59.7 | 53.5 | |

**B**

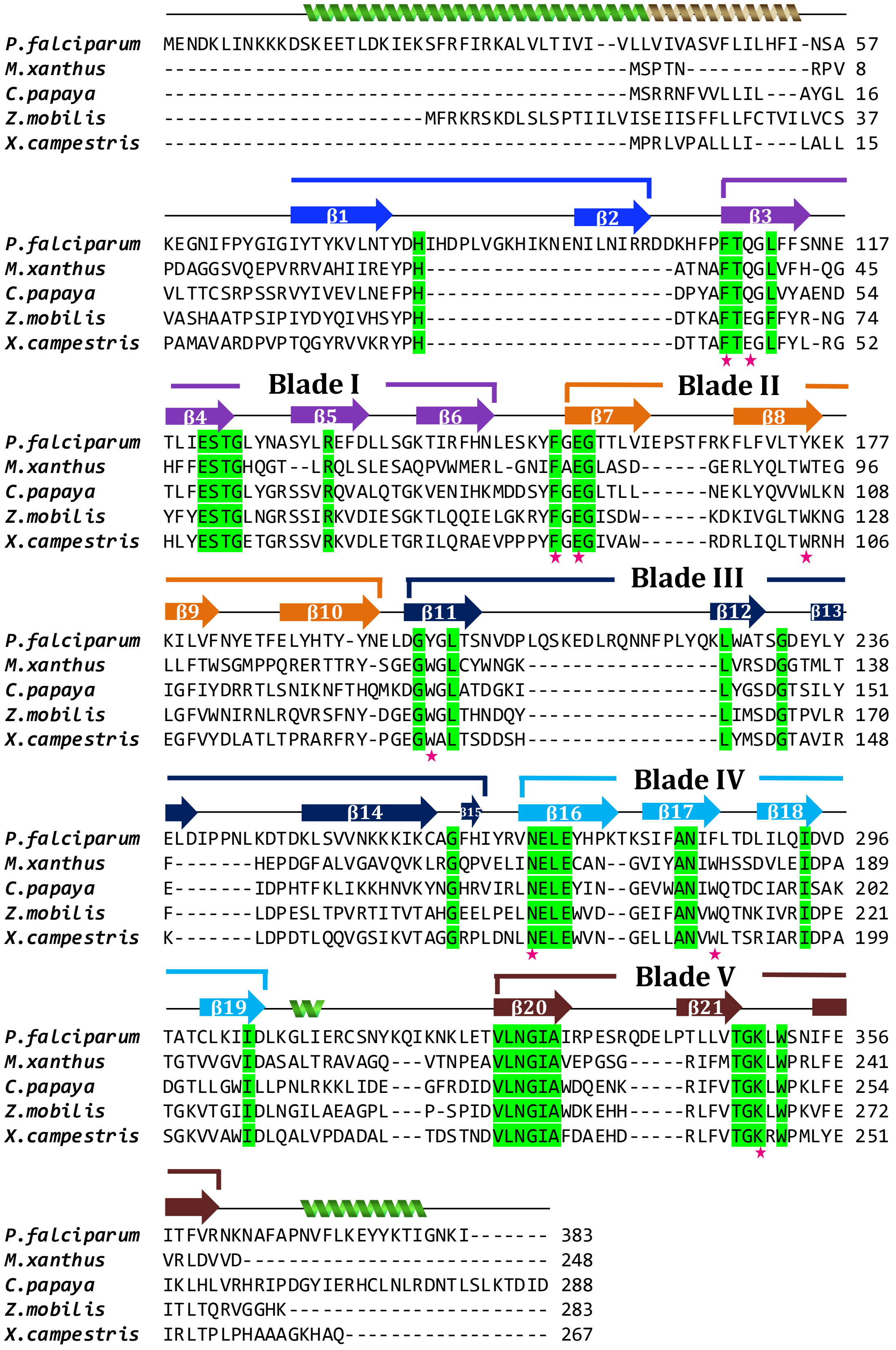

**Fig. S2. Amino acid sequence comparison of glutaminyl cyclase (QC) of different organisms**

**A.** Sequence identity (%, purple) and similarity (%, blue) between QCs from *P. falciparum* (*Pf*QC), different bacterial species and the plant *C. papaya* QC[1,10-12]. Uniprot IDs: A0A144A3U7 (*P. falciparum*; *Pf*), E7FH78 (Myxococcus *xanthus; Mx*), O81226 (*C. papaya; Cp*), A0A0H3G2U5 (*Zymomonas mobilis* subsp. Mobilis ZM4; *Zm*) and Q8P8M4 (*Xanthomonas campestris; Xc*).

**B.** Sequence alignment of QCs from *P. falciparum*, the plant *C. papaya* and different bacterial species. Secondary and tertiary structural elements (2 α-helices, 22 β-sheets and 5 blades) based on the predicted structure (**Fig. 1A**) are shown above the alignment. The α-helix in the predicted *P. falciparum* transmembrane domain (amino acids 32-54) is indicated in brown and conserved residues in green. Active site residues of *Pf*QCs, identified by aligning the amino acid sequence with the QCs from *C. papaya* and different bacterial species (*Z. mobilis, X. campestris and M. xanthus*)[11] , are marked as magenta stars.

### Fig. S3

**A**

**Number of QC spectra detected in *Plasmodium* proteomes**

| | blood stages | gametocytes | ookinetes | oocysts | sporozoites | liver stages |
|---|---|---|---|---|---|---|
| *P. falciparum* | 0 | 14 (n=3) | N.A. | 4 | 257 | N.A. |
| *P. berghei* | 0 | 0 | N.A. | N.A. | N.A. | N.A |
| *P. yoelii* | N.A. | N.A. | N.A. | 5 | 211 | 0 |

N.A. – not available

**B**

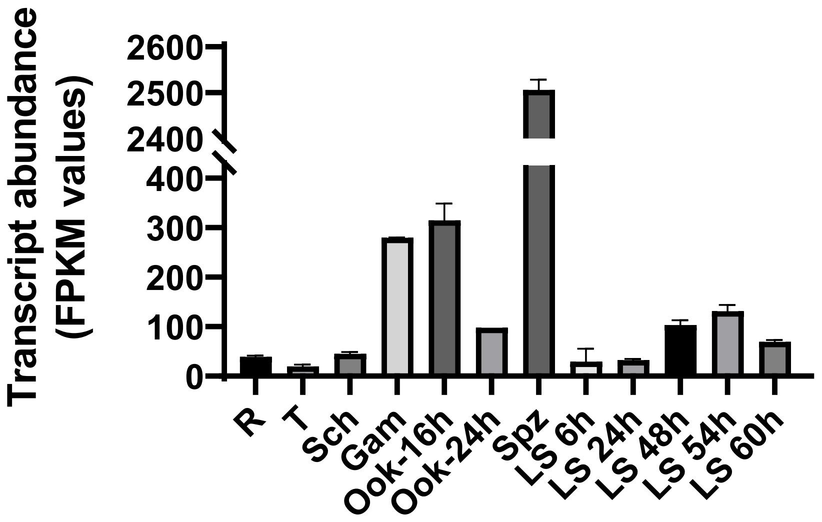


**C**

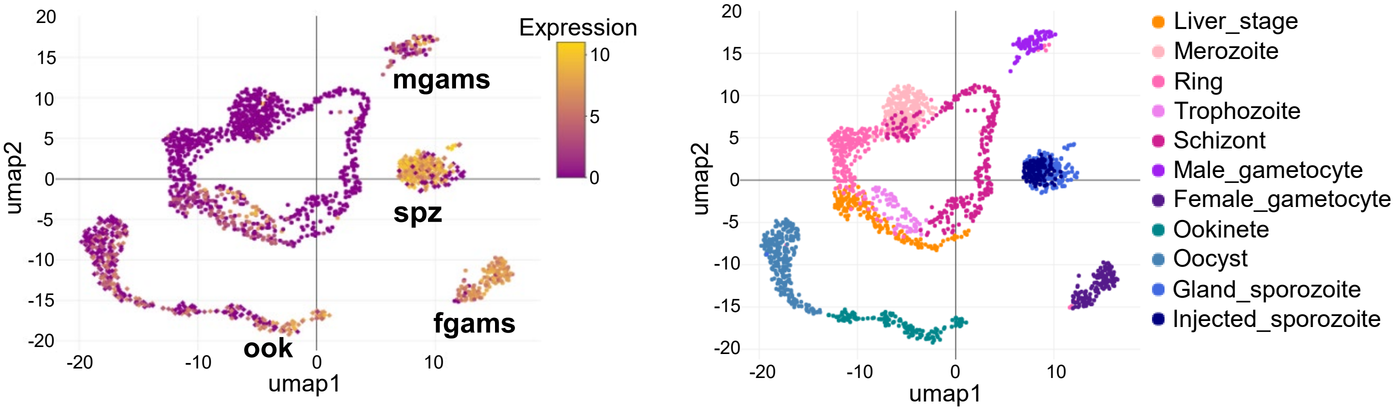


**Fig. S3. Published genome-wide proteomic and transcriptomic data of *Plasmodium* QC expression**

**A.** Number of QC-specific spectra detected in different life cycle stages of several *Plasmodium* species in genome-wide proteomic studies published in PlasmoDB (PlasmoDB v.46, www.plasmodb.org). **B.** *P. berghei qc* transcript levels (FPKM-values) from genome-wide transcriptomic studies across different life cycle stages[44,45]. R - rings, T - trophozoites, Sch - schizonts, Gam - gametocytes, Ook – ookinetes (*in vitro* cultured, 16 and 24 hour after activation of gametocytes) and LS - liver stages (*in vitro* cultured; at different hours after infection of hepatocytes). **C.** *P. berghei qc* expression evidence from single-cell transcriptomic studies, published in the Malaria Cell Atlas[46]. Left panel shows the level of *P. berghei qc* expression and right panel shows UMAP (uniform manifold approximation and projection) of single-cell transcriptomes sampled from the different life cycle stages. Highest expression is observed in female gametocytes (fgams), ookinetes (ook) and sporozoites (spz).

**Fig. S4**

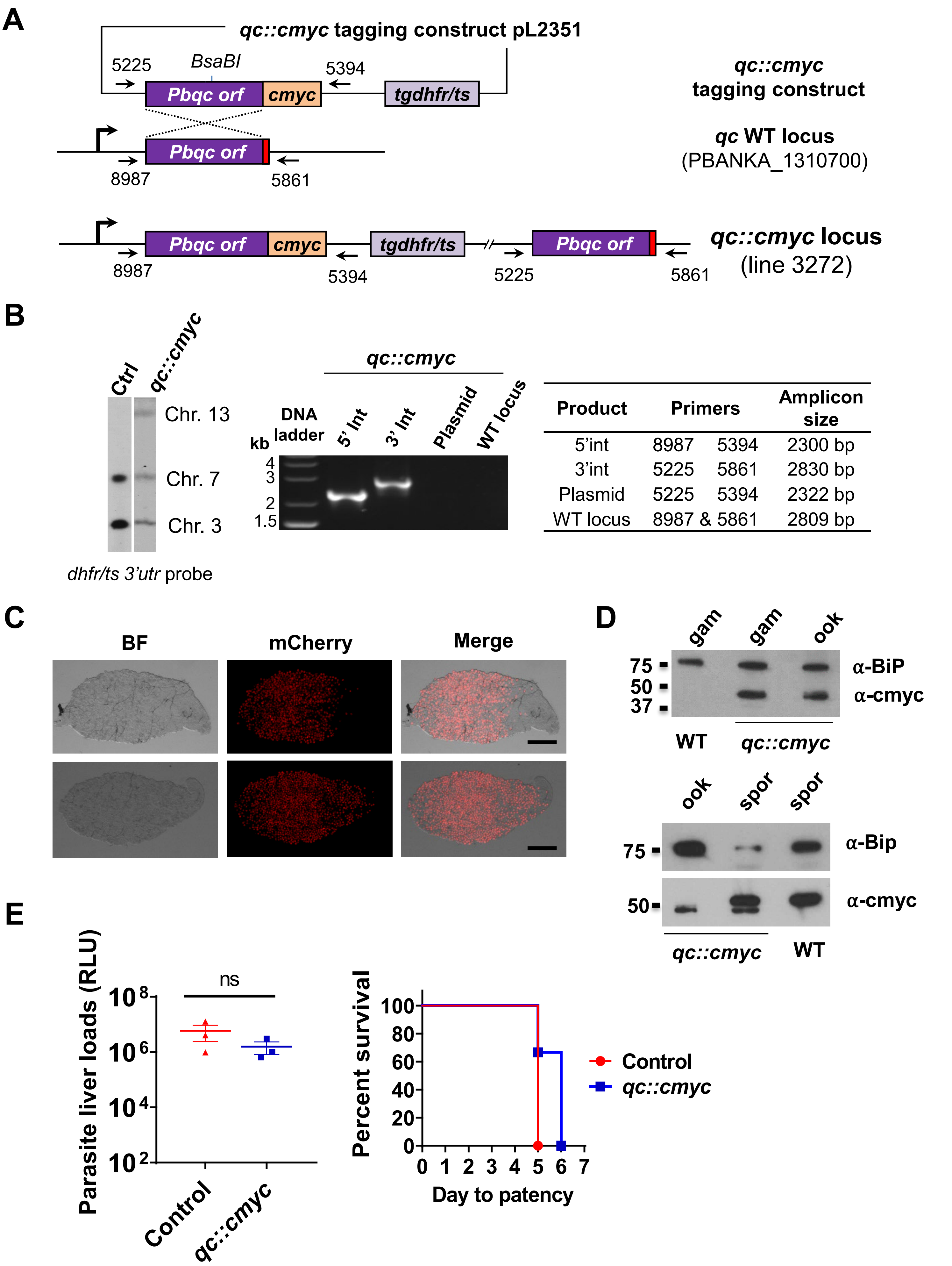
A
qc::cmyc tagging construct pL2351
5225
BsaBI
5394
Pbqc orf
cmyc
tgdhfr/ts
qc::cmyc
tagging construct
Pbqc orf
8987
5861
qc WT locus
(PBANKA_1310700)
Pbqc orf
cmyc
tgdhfr/ts
Pbqc orf
8987
5394
5225
5861
qc::cmyc locus
(line 3272)
B
Ctrl
qc::cmyc
Chr. 13
Chr. 7
Chr. 3
dhfr/ts 3'utr probe
qc::cmyc
kb
DNA ladder
5' Int
3' Int
Plasmid
WT locus
4
3
2
1.5
Product | Primers | Amplicon size
5'int | 8987 5394 | 2300 bp
3'int | 5225 5861 | 2830 bp
Plasmid | 5225 5394 | 2322 bp
WT locus | 8987 & 5861 | 2809 bp
C
BF
mCherry
Merge
D
gam
gam
ook
75
50
37
α-BiP
α-cmyc
WT
qc::cmyc
ook
spor
spor
75
α-Bip
50
α-cmyc
qc::cmyc
WT
E
Parasite liver loads (RLU)
ns
10^8
10^6
10^4
10^2
Control
qc::cmyc
Percent survival
100
80
60
40
20
0
0 1 2 3 4 5 6 7
Day to patency
Control
qc::cmyc

**Fig S4. Generation, genotyping and characterization of transgenic *P. berghei* parasites expressing C-terminal cmyc-tagged QC**

**A.** Schematic representation of the DNA construct pL2351 used for generation of the transgenic line *Pbqc::cmyc* (line 3272) expressing QC C-terminally tagged with cmyc. The construct contains the *tgdhfr* selectable marker cassette (SM) and a region of the target gene for integration (purple) in the endogenous *qc* gene (middle panel) by single cross-over homologous recombination. Integration of the construct into the target gene results in a C-terminal cmyc-tagged copy of the target gene (lower panel). BsaBI: restriction enzyme used for linearization of plasmid before transfection. Black arrows: location of primers used for diagnostic PCR; red box: stop codon.

**B.** Genotype analysis of *Pbqc::cmyc* parasites by Southern analysis of chromosomes (chr.) separated by pulsed-field gel electrophoresis (PFGE) and diagnostic PCR analysis. Hybridisation of PFG-separated chr. of *Pbqc::cmyc* with the *Pbdhfr 3'utr* probe confirms integration of pL2351 into the *Pbqc* gene on chr. 13. In addition, this probe recognizes the endogenous *dhfr/ts* 3'utr on chr. 7 and the *mCherry* and *luciferase* reporter cassettes integrated into chr. 3. As a control parasite (ctrl), line 2117cl1 is used with the *hdhfr::yfcu* cassette integrated into chr. 3. PCR analysis shows correct integration at both 5' and 3' regions (5'int and 3'int), absence of plasmid and wild type *qc* locus; see **A** for primer numbers and locations. Primers sequences are shown in **Table S1**.

**C.** Oocyst in mosquito midguts (day 17) infected with *Pbqc::cmyc* parasites. BF – brightfield. WT - wild type. Scale bar: 200 µm.

**D.** *QC* expression in wild type (WT) and *Pbqc::cmyc* gametocytes (gam), ookinetes (ook) and sporozoites (spor) by Western analysis using anti-cmyc antibodies. As a loading control anti-Bip antibody was used. Staining with anti-cmyc showed a (non-specific) hybridization to a fragment of 55-60 kD in both WT and *Pbqc::cmyc* sporozoites, in addition to the specific 50 kD *Pb*QC::cmyc fragment in *Pbqc::cmyc* parasites.

**E.** Infectivity of *Pbqc::cmyc* sporozoites as determined by parasite liver load (left panel) and prepatent period (right panel) after intravenous injection of mice (n=3) with $1 \times 10^4$ sporozoites. Parasite liver loads were determined at 44 h by measuring *in vivo* luciferase activity and depicted as relative light units (RLU) (ns=non-significant; Mann-Whitney test, P value shown in **Table S3**). Prepatent period (day at which a parasitemia of 0.5–2% is observed) is shown as Kaplan-Meier curves (non-significant; Log-Rank (Mantel-Cox) test, P value shown in **Table S3**).

### Fig. S5

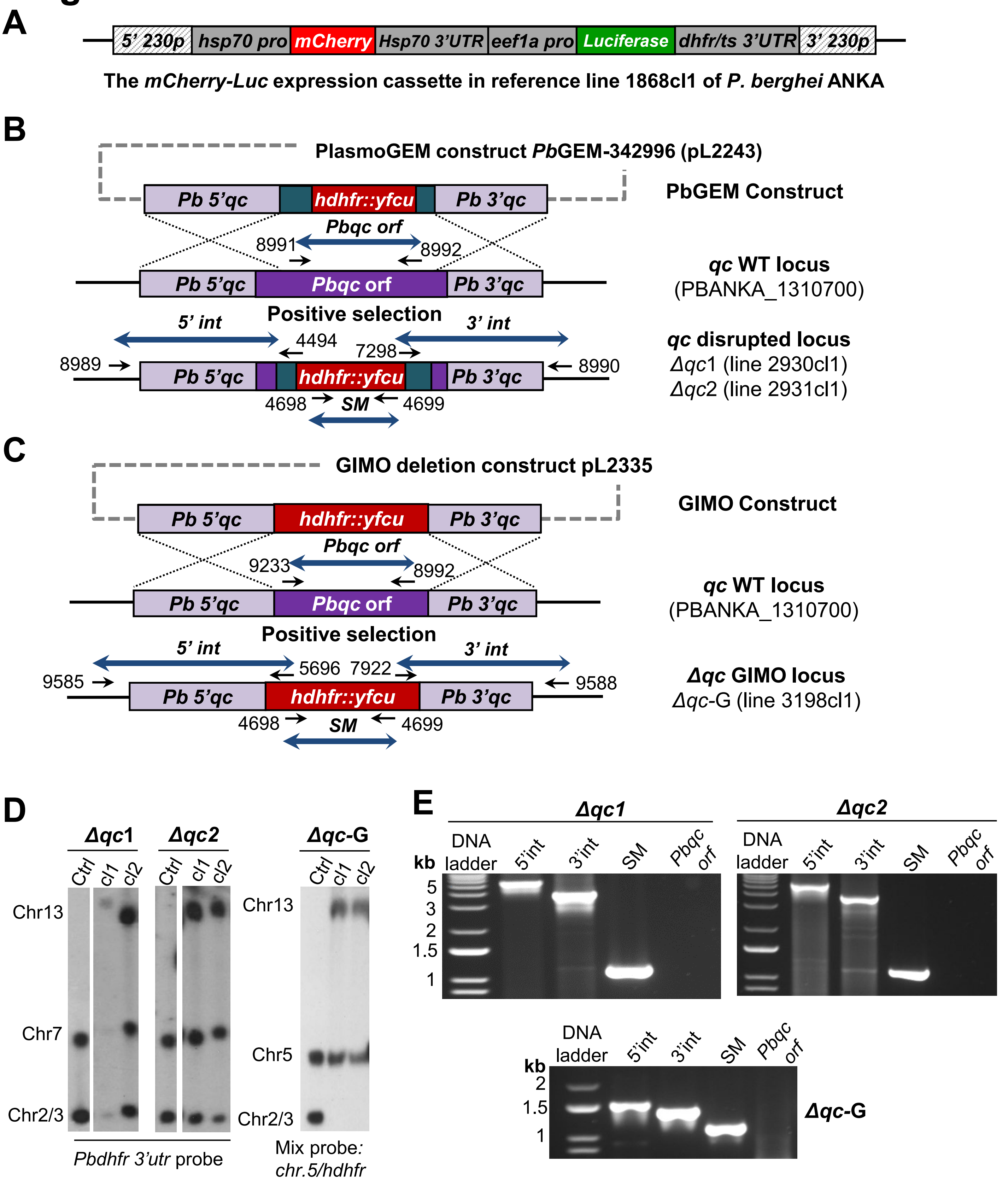


F

| Template | Product | Primers | | Amplicon size |
|---|---|---|---|---|
| ***Δqc1*** | 5'int | 8989 | 4494 | 5419 bp |
| (2930cl1) | 3'int | 7289 | 8990 | 3866 bp |
| ***Δqc2*** | SM | 4698 | 4699 | 1108 bp |
| (2931cl1) | ORF | 8991 | 8992 | 624 bp |

| Template | Product | Primers | Amplicon size |
|---|---|---|---|
| | 5'int | 9585 & 5696 | 1504 bp |
| ***Δqc-G*** | 3'int | 7922 & 9588 | 1398 bp |
| (3198cl1) | SM | 4698 & 4699 | 1108 bp |
| | ORF | 9233 8992 | 1424 bp |

**Fig. S5. Generation, genotyping and characterization of three *P. berghei* QC-null mutants (*PbΔqc*)**
**A.** Schematic representation of the *Pb230p* locus of the reference reporter *P. berghei* parasite line (1868cl1), containing *mCherry* and *firefly luciferase* reporter genes under the constitutive *hsp70* and *eef1a* promoters. This line was used to generate the QC-null mutants.
**B.** Schematic representation of the generation of the QC-null mutants *PbΔqc*1 (2930cl1) and *PbΔqc*2 (2931cl1). The PlasmoGEM construct PbGEM-342996 (pL2243) was used to replace the *Pbqc* open reading frame (*orf*) with the positive/negative selectable marker (SM; h*dhfr::yfcu*) cassette by double cross-over homologous recombination, resulting in the *PbΔqc*1 and *PbΔqc*2 lines after positive selection with pyrimethamine and cloning. Black arrows: location and primer name used for diagnostic PCR (see **F**).
**C.** Schematic representation of the generation of the QC-null mutant *PbΔqc*-GIMO (3198cl1). The GIMO deletion construct (pL2335) was used to replace the *Pbqc orf* with the positive/negative SM (h*dhfr::yfcu*) cassette by double cross-over homologous recombination, resulting in the generation of *PbΔqc*-GIMO after positive selection with pyrimethamine and cloning. Black arrows: location and primers used for diagnostic PCR (see **F**).
**D.** Genotype analysis of *PbΔqc*1, *PbΔqc*2 and *PbΔqc*-GIMO parasites by Southern analysis of chromosomes (chr.) separated by pulsed-field gel electrophoresis (PFGE). Hybridisation of PFGE-separated chr. of *PbΔqc*1 and *PbΔqc*2 with the *Pbdhfr 3'utr* probe confirms integration of pL2243 in the *Pbqc* gene on chr. 13. In addition, this probe recognizes the endogenous *dhfr/ts* gene on chr. 7 and the reporter *luciferase* cassette integrated into chr. 3. Hybridisation of chr. of *PbΔqc*-GIMO with a mixture of a *hdhfr* probe and a probe specific for chr. 5 confirms the integration of pL2335 into the *Pbqc* gene on chr. 13. As an additional control (ctrl), parasite line 2117cl1 that has *hdhfr::yfcu* SM integrated into chr. 3. Blood stage growth of asexual blood stages was determined during the cloning period and was comparable to growth of wild type *P. berghei* ANKA parasites with a mean multiplication rate of 10x per 24 hour. Number of clones (n) per line: *PbΔqc*1 n=3, *PbΔqc*2 n=6 and *PbΔqc*-GIMO n=3. Clone 1 of all three lines was used for further analyses.
**E.** Genotype analysis of *PbΔqc*1, *PbΔqc*2 and *PbΔqc*-GIMO parasites by diagnostic PCR confirms the absence of *Pbqc* open reading frame (*orf*) in *PbΔqc*1, *PbΔqc*2 and *PbΔqc*-GIMO, the correct integration of the constructs at both the 5' and 3' regions (5'int and 3'int), and the presence of the positive/negative SM; see **B, C** for primer numbers and locations. Primer sequences are shown in **Table S1** and expected PCR fragment sizes and the primer names are listed in **F**.
**F.** Expected fragment sizes and names of primers used for diagnostic PCR in **E.**

Fig. S6

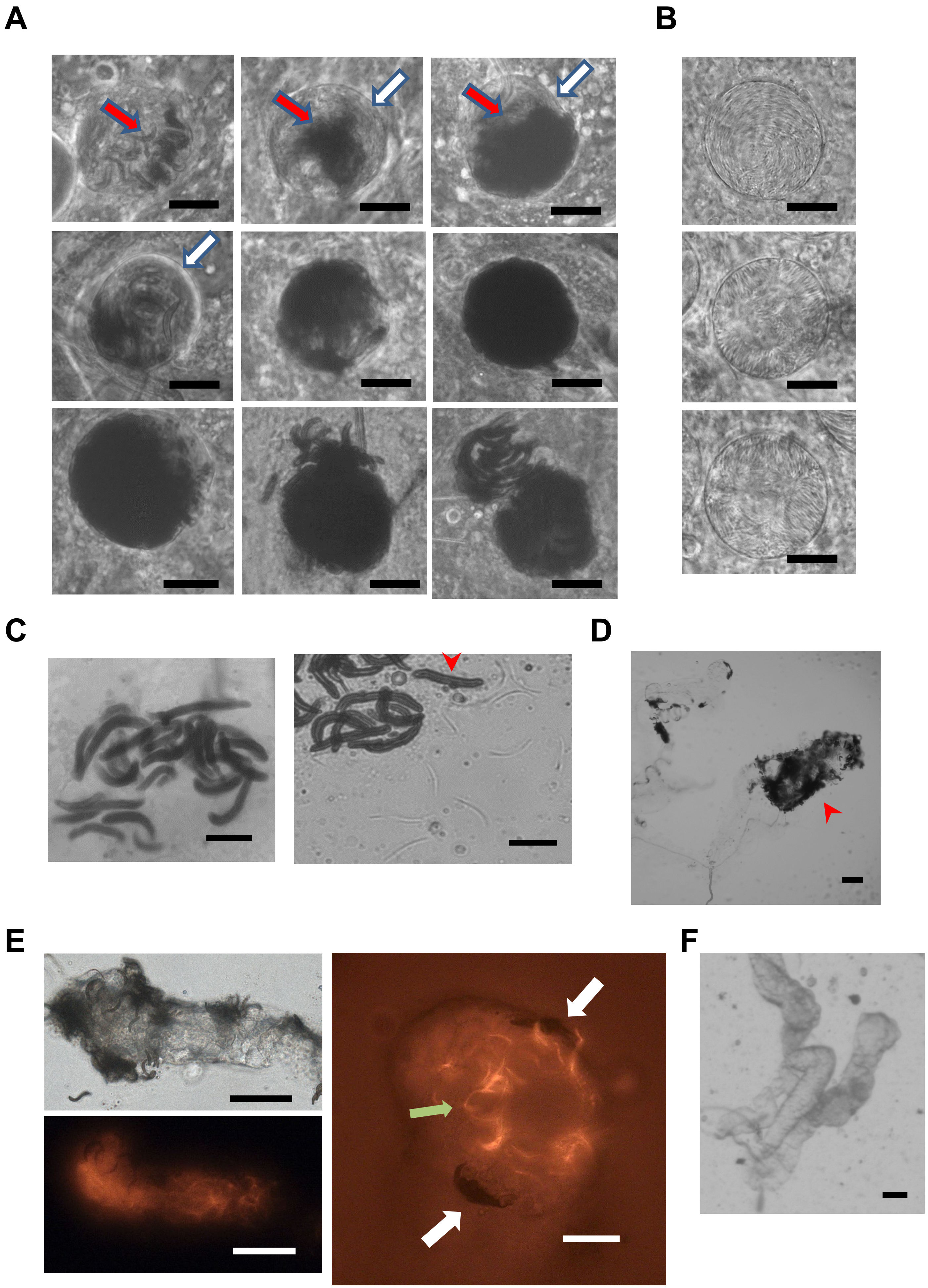

**Fig. S6. Light- and fluorescence microscope images of melanized of oocysts and sporozoites of *P. berghei* QC-null mutants**

**A.** QC-null melanized oocysts (day 14) showing different degrees of melanization. No signs of oocyst melanization was detected before day 10 post-infection and melanization was observed mainly inside sporulated oocysts (red arrows), showing the presence of aberrant, enlarged and dark-colored sporozoites, either still inside oocysts or in the process of release from oocysts. No distinct melanin deposition at the outside of the oocyst wall (white arrows). Scale bar: 20 µm.

**B.** QC-null non-melanized oocysts (day 14) with typical features of wild type sporozoite formation. Scale bar: 20 µm.

**C.** Left: Clusters of melanized QC-null sporozoites found in the hemocoel (day 21) Scale bar: 10 µm. Right: Partially crushed salivary gland (day 21) of a QC-null infected mosquito showing melanized (red arrows) and non-melanized (blue arrows) sporozoites. Scale bar: 10 µm.

**D.** Lobes of salivary glands of a QC-null infected mosquitoes (day 21), showing clusters of melanized sporozoites (red arrows). Scale bar: 20 µm.

**E.** Left: Lobes of salivary glands of QC-null infected mosquitoes (day 21), showing the clustering of mCherry-negative sporozoites at the outside of the gland. Scale bar: 50 µm and the presence of mCherry-expressing sporozoites inside the gland. Right: A magnification of a QC-null-infected salivary gland (day 21), visualizing two clusters of mCherry-negative sporozoites at the outside of the gland (white arrows) and individual mCherry-positive sporozoites inside the gland (green arrow). Scale bar: 20 µm.

**F.** Lobes of salivary glands of WT-infected mosquitoes (day 21), showing the absence of melanized sporozoites. Scale bar: 20 µm.

Fig. S7

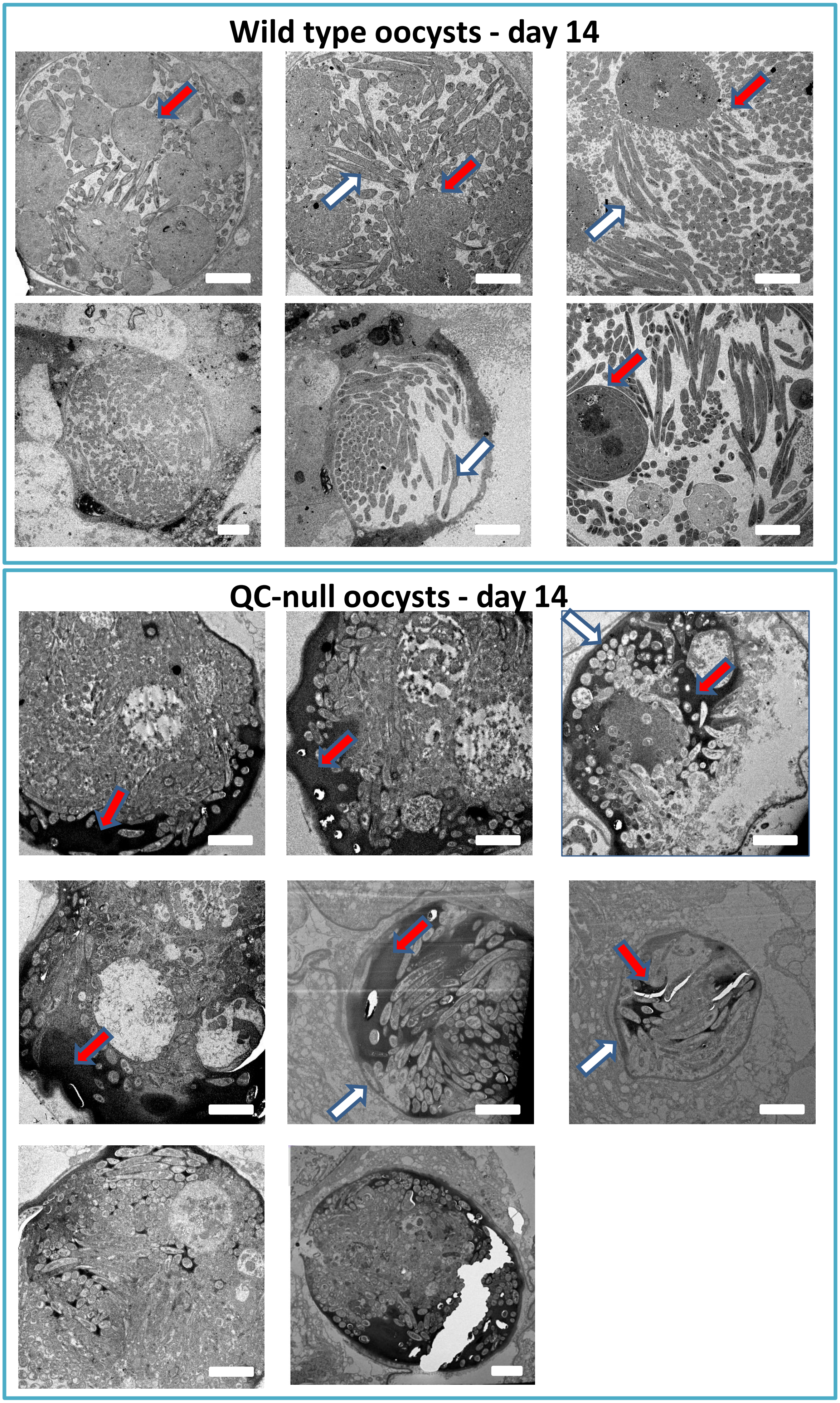

**Fig. S7. Transmission electron microscopy images (TEM) of melanized of wild type (WT) and QC-null oocyst (day 14)**

Upper panel: 6 images of different WT oocysts showing sporoblasts (red arrows), sporozoite formation and sporozoites (white arrows). Scale bar: 5 µm.

Lower panel: 8 images of different QC-null oocysts showing melanin deposition inside oocysts (red arrows), often surrounding sporozoites. No distinct melanin deposition at the outside of the oocyst wall (white arrows). Scale bar: 5 µm.

Fig. S8

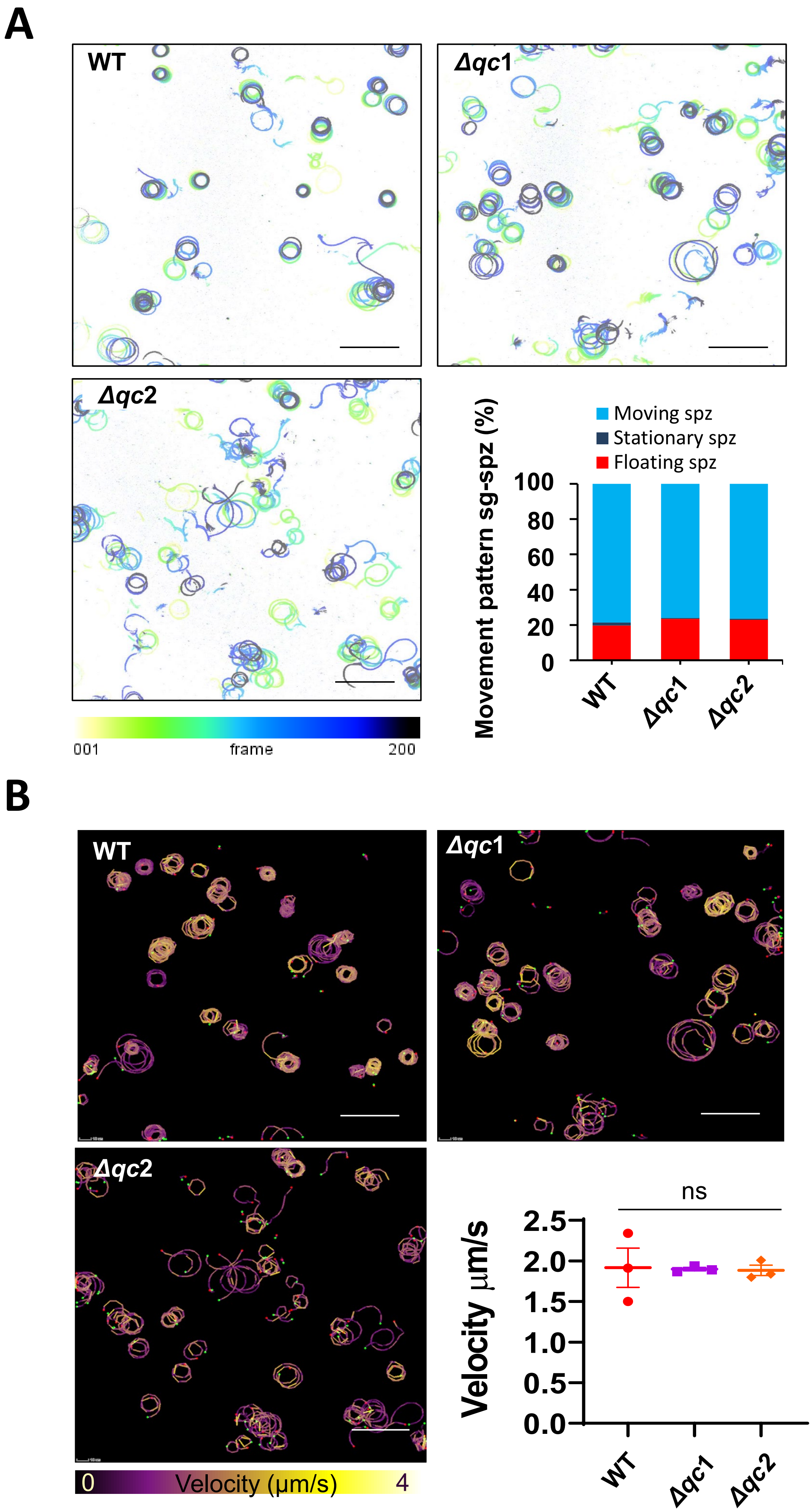

A
WT
Δqc1
Δqc2
001
frame
200
Moving spz
Stationary spz
Floating spz
Movement pattern sg-spz (%)
100
80
60
40
20
0
WT
Δqc1
Δqc2
B
WT
Δqc1
Δqc2
0
Velocity (μm/s)
4
ns
Velocity μm/s
2.5
2.0
1.5
1.0
0.5
0.0
WT
Δqc1
Δqc2

**Fig. S8. Gliding motility of QC-null salivary gland (sg) sporozoites**

**A.** Movie projections of movement patterns (lines and circles) of QC-null and wild type (WT) sg-sporozoites (control). Quantitative analysis of the movement patterns (moving, stationary and floating) of QC-null sg-sporozoites did not show significant differences (p=0.99, student t- test) with those of WT sporozoites. Scalebar: 50 µm.

**B.** Movie projections (circles) of the velocity of QC-null sg-sporozoites and WT sg-sporozoites (control) on glass surface (n=3). The velocity of QC-null sg-sporozoites was not significantly different (p>0.99; Mann-Whitney test) with that of the WT sg-sporozoites. Scalebar: 50 µm.

Fig. S9

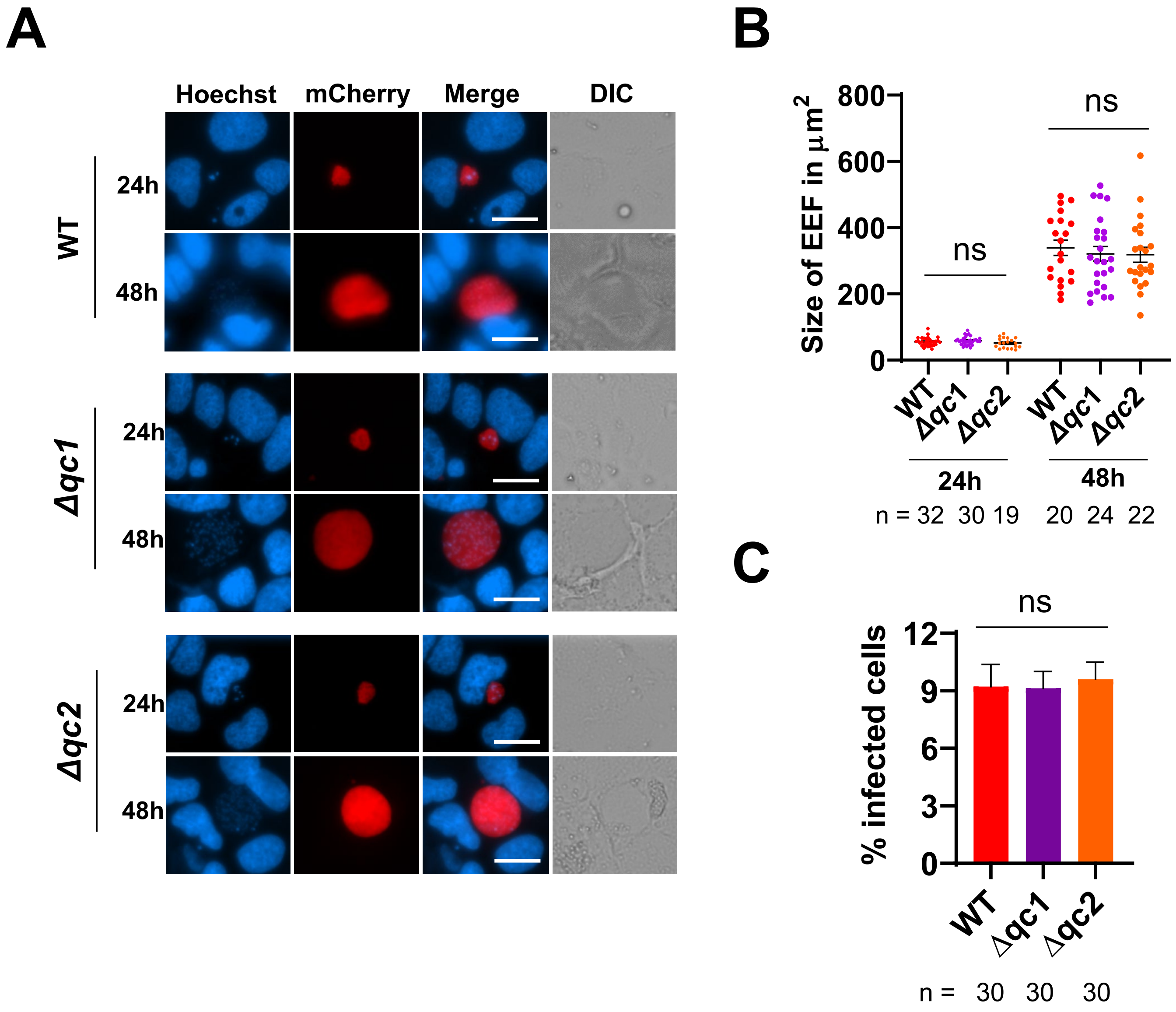


**Fig. S9. Infectivity of salivary gland (sg) sporozoites of different *Pbqc* mutants**

**A.** Fluorescent imaging of live, mCherry expressing QC-null and WT liver stages in cultured Huh7 hepatocytes. Liver-stages express cytoplasmic mCherry (under control of the constitutive *hsp70* promoter). Images were taken at 24 and 48 h after infection of hepatocytes with $5x10^4$ sporozoites. Nuclei stained with Hoechst-33342. Scale bar: 20µm.

**B.** Sizes (mean ± SEM) of QC-null and WT liver stages (n=number of liver stages) in cultured Huh7 hepatocytes at 24 and 48 h after infection as measured by determining the area of the parasite at its greatest circumference using the mCherry-positive area ($µm^2$). No significant (ns) differences in size were observed between QC-null and WT liver stages (Mann-Whiney test, statistical significance is shown relative to WT. P values are shown in **Table S3**).

**C.** Infectivity of WT and QC-null sporozoites in cultured Huh7 hepatocytes, shown as the percentage of infected cells hepatocytes (n=number of microscopic fields at 40X magnification). No significant (ns) differences were observed between QC-null and WT parasites (Mann-Whitney test, statistical significance is shown relative to WT. P values are shown in **Table S3**).

### Fig. S10

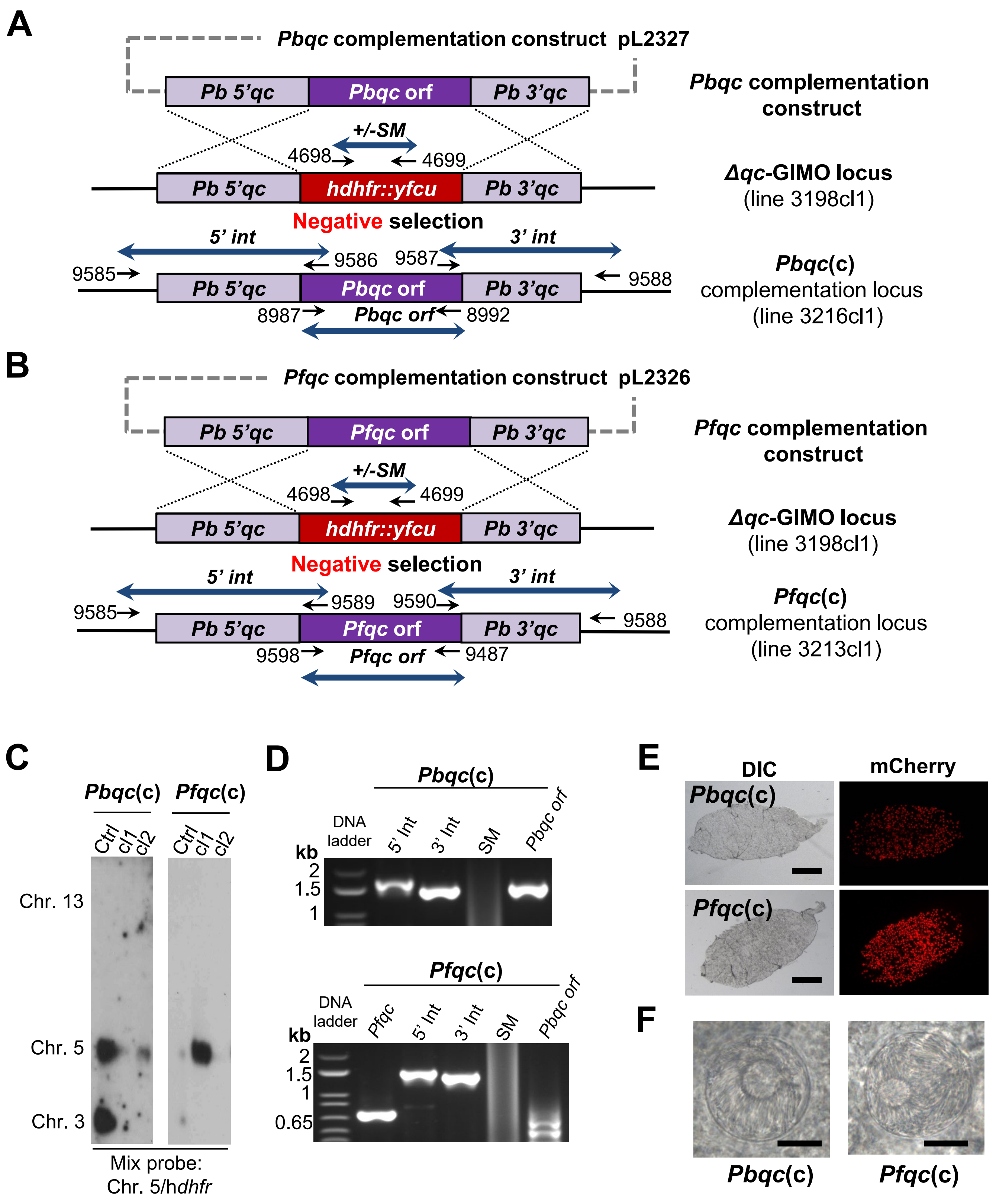


**G**

| Template | Product | Primers | | Amplicon size |
|---|---|---|---|---|
| | 5'int | 9585 | 9586 | 1698 bp |
| ***Pbqc*(c)** (3216cl1) | 3'int | 9587 | 9588 | 1460 bp |
| | SM | 4698 | 4699 | 1108 bp |
| | *Pbqc orf* | 8987 | 8992 | 1492 bp |

| Template | Product | Primers | | Amplicon size |
|---|---|---|---|---|
| | *Pfqc orf* | 9598 | 9487 | 725 bp |
| | 5'int | 9585 | 9589 | 1481 bp |
| ***Pfqc*(c)** (3213cl1) | 3'int | 9590 | 9588 | 1386 bp |
| | SM | 4698 | 4699 | 1108 bp |
| | *Pbqc orf* | 9233 | 8992 | 1492 bp |

**Fig. S10. Generation, genotyping and characterisation QC-null mutants complemented with *P. berghei* QC (*Pbqc*(c)) or *P. falciparum* QC (*Pfqc*(c))**

**A.** Schematic representation of the generation of the complemented line *Pbqc*(c) *(*3216cl1). The *Pbqc* complementation construct pL2327 was used to replace the positive/negative selectable marker (SM; h*dhfr::yfcu*) cassette in the *Δqc*-GIMO mutant after negative (5-FC) selection. Construct pL2327 integrates by double cross-over homologous recombination using the same targeting regions of construct pL2335 (**Fig. S5**), resulting in the introduction of the *Pbqc* open reading frame (*orf*) under the control of *Pbqc* regulatory sequences. Black arrows: location and primers used for diagnostic PCR (see **G**).

**B.** Schematic representation of the generation of the complemented line *Pfqc*(c) (3213cl1). The *Pfqc* complementation construct (pL2326) was used to replace the positive/negative selectable marker (SM; h*dhfr::yfcu*) cassette in the *Δqc*-GIMO mutant after negative (5-FC) selection. Construct pL2326 integrates by double cross-over homologous recombination using the same targeting regions of construct pL2335 (**Fig. S5**), resulting in the introduction of the *Pfqc* open reading frame (*orf*) under the control of *Pbqc* regulatory sequences. Black arrows: location and primers used for diagnostic PCR (see **G**).

**C.** Genotype analysis of *Pbqc*(c) and *Pfqc*(c) parasites by Southern analysis of chromosomes (chr.) separated by pulsed-field gel electrophoresis (PFGE). Hybridisation of PFGE-separated chr. of *Pbqc*(c) and *Pfqc*(c) with a mixture of a *hdhfr* probe and a probe specific for chr. 5 confirms the integration of pL2327and pL2326 respectively into the *Pbqc*-GIMO locus on chr. 13. As an additional control (ctrl), parasite line 2117cl1 is used with the *hdhfr::yfcu* SM integrated into chr. 3.

**D.** Genotype analysis of *Pbqc*(c) and *Pfqc*(c) parasites by diagnostic PCR confirms the absence of the *hdhfr::yfcu* SM, the correct integration of the constructs at both the 5' and 3' regions (5'int and 3'int) and the presence of the *Pbqc orf* or the *Pfqc orf*. See **A** and **B** for primer numbers and locations. Primer sequences are shown in **Table S1** and expected PCR fragment sizes and primer names are listed in the Table in **G**.

**E.** Examples of bright field and mCherry fluorescent images of midguts of *A. stephensi*, 14 days after infection with *Pbqc*(c) or *Pfqc*(c), showing mCherry-expressing oocysts and the absence of melanized, dark-coloured oocysts. Scale bar: 200 µm.

**F.** Light microscopic images of *Pbqc*(c) and *Pbfc*(c) oocysts (day 14) with typical features of WT sporozoite formation. Scale bar: 20 µm.

**G.** Expected fragment sizes and primer names used for diagnostic PCR in **D.**

#### Fig. S11

**A**

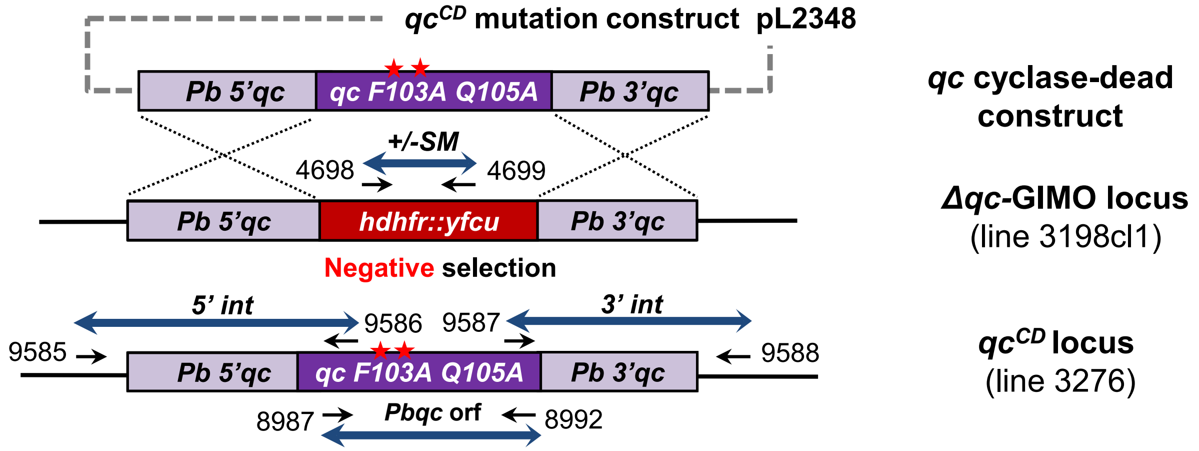


**B**

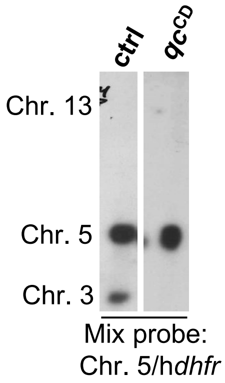


**C**

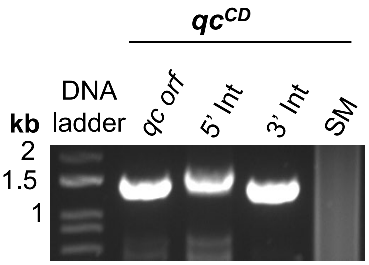


| Template | Product | Primers | | Amplicon size |
|---|---|---|---|---|
| *qc^CD* (3276) | 5'int | 9585 | 9586 | 1698 bp |
| | 3'int | 9587 | 9588 | 1460 bp |
| | SM | 4698 | 4699 | 1108 bp |
| | *Pbqc orf* | 8987 | 8992 | 1492 bp |

**D**

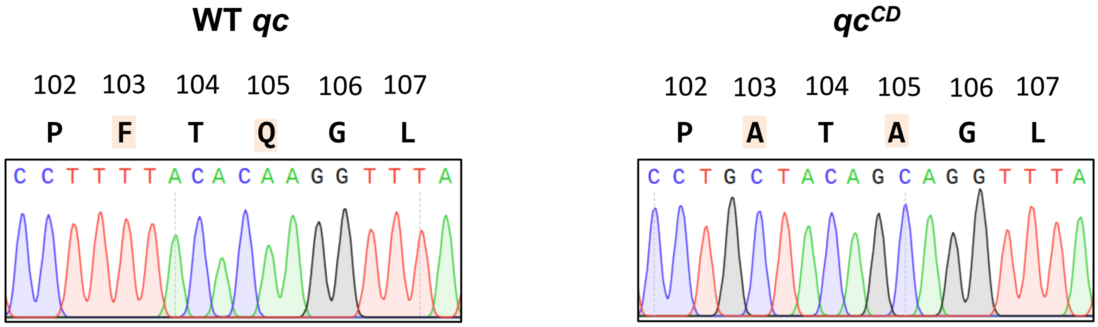


**E**

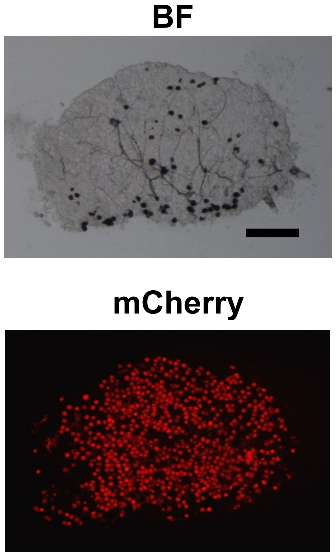


**F**

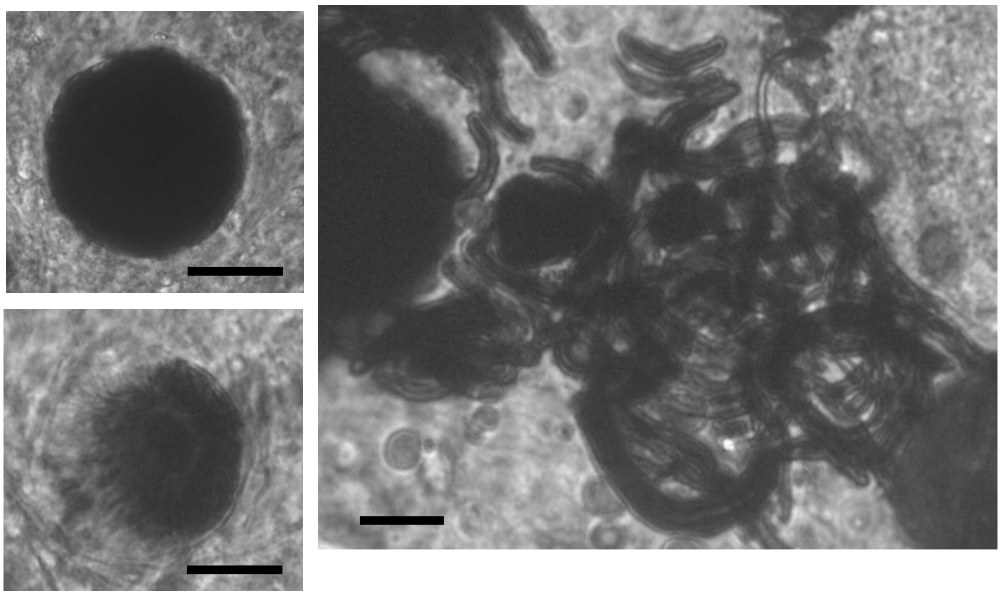

**Fig. S11. Generation, genotyping and characterisation of the *P. berghei* cyclase-dead mutant (*qc*$^{cD}$)**

**A.** Schematic representation of the generation of the cyclase-dead mutant *qc*$^{CD}$ line (line 3276), expressing a cyclase-dead (CD) QC containing two point mutations in the catalytic site (F103A/Q105A; red arrows). The cyclase-dead complementation construct (pL2348) was used to replace the positive/negative selectable marker (SM; h*dhfr::yfcu*) cassette in the *Δqc*-GIMO (3198cl1) line after negative (5-FC) selection. Construct pL2348 integrates by double cross-over homologous recombination using the same targeting regions employed in generating the construct pL2335 (**Fig. S5**), resulting in the introduction of the cyclase-dead *qc* open reading frame (*orf*) under the control of *qc* regulatory sequences. Black arrows: location and primer name used for diagnostic PCR (see **D**).

**B.** Genotype analysis of *qc*$^{CD}$ parasites by Southern analysis of chromosomes (chr.) separated by pulsed-field gel electrophoresis (PFGE). Hybridisation of PFG-separated chr. of *qc*$^{CD}$ with a mixture of *hdhfr* and a probe specific for chr. 5 confirms the integration of pL2348 into the *qc*-GIMO locus on chr. 13. As an additional control (ctrl), parasite line 2117cl1 is used with the *hdhfr::yfcu* SM integrated into chr. 3.

**C.** Genotype analysis of *Pbqc*$^{CD}$ parasites by diagnostic PCRs confirms the absence of the *hdhfr::yfcu* SM, the correct integration of the construct at both the 5' and 3' regions (5'int and 3'int) and the presence of the *Pbqc* F103A Q105A. See **A** for primer numbers and locations. Primer sequences are shown in **Table S1** and expected PCR fragment sizes and the primer numbers are listed in the Table.

**D.** Sanger-sequence DNA chromatograms of PCR fragments from genomic DNA of WT and *qc* cyclase dead (*qc*$^{CD}$) parasites, confirming the replacement of catalytic site amino acids phenyl alanine (F103) and glutamine (Q105) to alanines (F103A Q105A, highlighted).

**E.** Brightfield and fluorescence microscope images of mCherry-expressing *qc*$^{CD}$ oocysts (day 14) showing melanised and normal oocysts. Scale bar: 200 µm.

**F.** Brightfield microscope images of melanised *qc*$^{CD}$ oocysts (day 14; scale bar: 20 µm) and brightfield microscope image of melanised *qc*$^{CD}$ hemocoel sporozoites (day 23). Scale bar: 10 µm.

#### Fig. S12

A

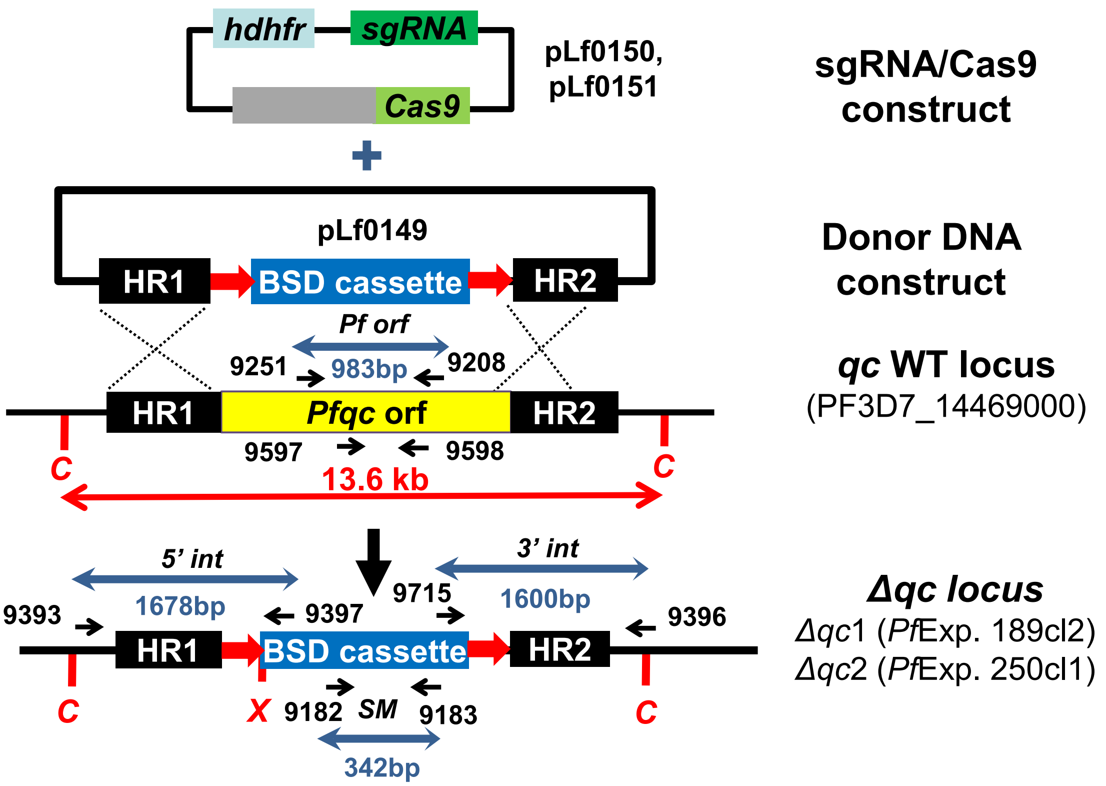


B

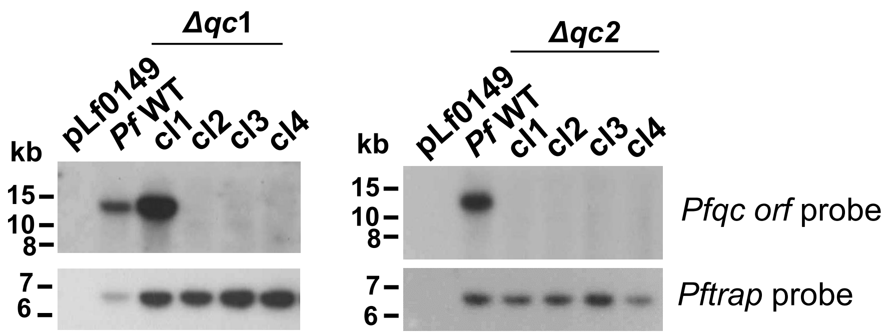


C

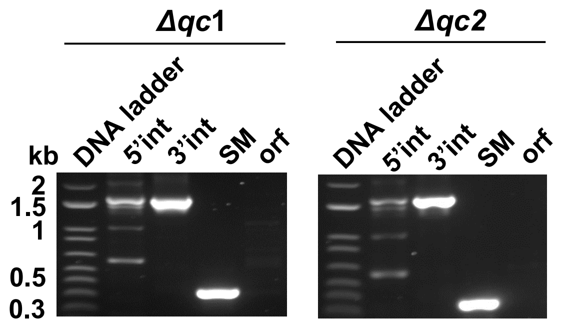


| Template | Product | Primers | Amplicon size |
|---|---|---|---|
| *Δqc1* (189cl2) and *Δqc2* (250cl1) | 5'int | 9393 and 9397 | 1678bp |
| | 3'int | 9175 and 9 | 1638bp |
| | SM | 9182 and 9183 | 342bp |
| | ORF | 9251 and 9208 | 983bp |

D

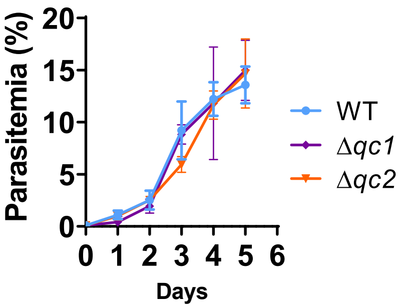


E

| Parasites | Gametocytes stage V (%) [a] | No.of exflagellations mean (SD) [b] |
|---|---|---|
| WT (n=4) | 2.1% | 665 (120) |
| *Δqc1* | 1.9% (2 exp.) | 564 (1 exp.) |
| *Δqc2* | n.d. | 569 (2 exp) |

**Fig. S12. Generation and genotyping of *P. falciparum* QC-null mutants (*PfΔqc*)**

**A.** Schematic representation of the two sgRNA/Cas9 plasmids (pLf0150 and pLf0151) and the donor DNA plasmids (pLf0149) used to generate two *PfΔ*qc lines (exp. 189 and exp. 250). The *Pfqc* homology regions (HR1, HR2), the BSD selectable marker cassette, location of primers (in black arrows), restriction fragments sizes and PCR amplicons (in blue) are indicated (C: *Cla*I; in red). XhoI cuts in the integrated region of donor DNA and BamHI cuts the plasmid outside the integration region. The figure is not drawn to scale. Primer sequences (in black and bold) are shown in **Table S1**. WT - wild type; BSD – blasticidin S deaminase selectable marker (SM); *hdhfr* – SM in Cas9 plasmid; *orf* – open reading frame.

**B.** Southern analysis of restriction digested DNA of WT and four clones (cl) of *PfΔqc*1 and *PfΔqc*2 parasites confirms *Pfqc* gene deletion in *Pfqc1* cl2-4 and *Pfqc2* cl1-4. DNA was hybridized with a probe targeting the *Pfqc orf* region. The hybridization fragments of WT (13.6 kb) or of *PfΔqc* single crossover mutants (9 kb or 10 kb) are absent in the positive clones, indicating removal of the *Pfqc orf* by double cross-over integration of the donor DNA construct. As a control for DNA loading, a *Pftrap* probe was used, recognizing a 6.8 kb fragment. Clone 2 of *PfΔqc*1 and clone 1 of *PfΔqc*2 were used for all analyses.

**C.** Diagnostic PCR confirms correct 5'- and 3'- integration of the donor DNA construct, presence of the BSD SM and absence of the *Pfqc orf* in *PfΔqc*1 and *PfΔqc*2. Primer locations and product sizes are shown in **(A),** primer sequences in **Table S1** and expected PCR fragment sizes and the primer names are listed in the Table.

**D.** Growth of asexual blood-stages of *PfΔqc*1, *PfΔqc*2 and WT *Pf*NF54. Parasitemia (mean and S.D of 4 independent cultures) is shown during a 5-day culture period (in the semi-automated culture system). Cultures were initiated with a parasitemia of 0.1%.

**E.** Table showing gametocyte, oocyst, and sporozoite production of *PfΔqc*1, *PfΔqc*2 and WT *Pf*NF54 lines.

[a] Mean percentage of stage V male and female gametocytes (per 100 red blood cells) in day 14 gametocyte cultures in 5 experiments (exp) with standard deviation (SD).

[b] Mean number of exflagellating male gametocytes (per $10^5$ red blood cells) at 10-20 min after activation in day 14 gametocyte cultures.

[c] Mean number of oocysts per mosquito at day 10-12 after feeding. Range corresponds to the mean number of oocysts in multiple experiments (Per line; 10-30 mosquitoes per exp.).

[d] Mean number of salivary gland sporozoites per mosquito at 18 - 24 after feeding. Range corresponds to the mean number of sporozoites in multiple experiments (per line; 50 - 90 mosquitoes per exp.).

Fig. S13

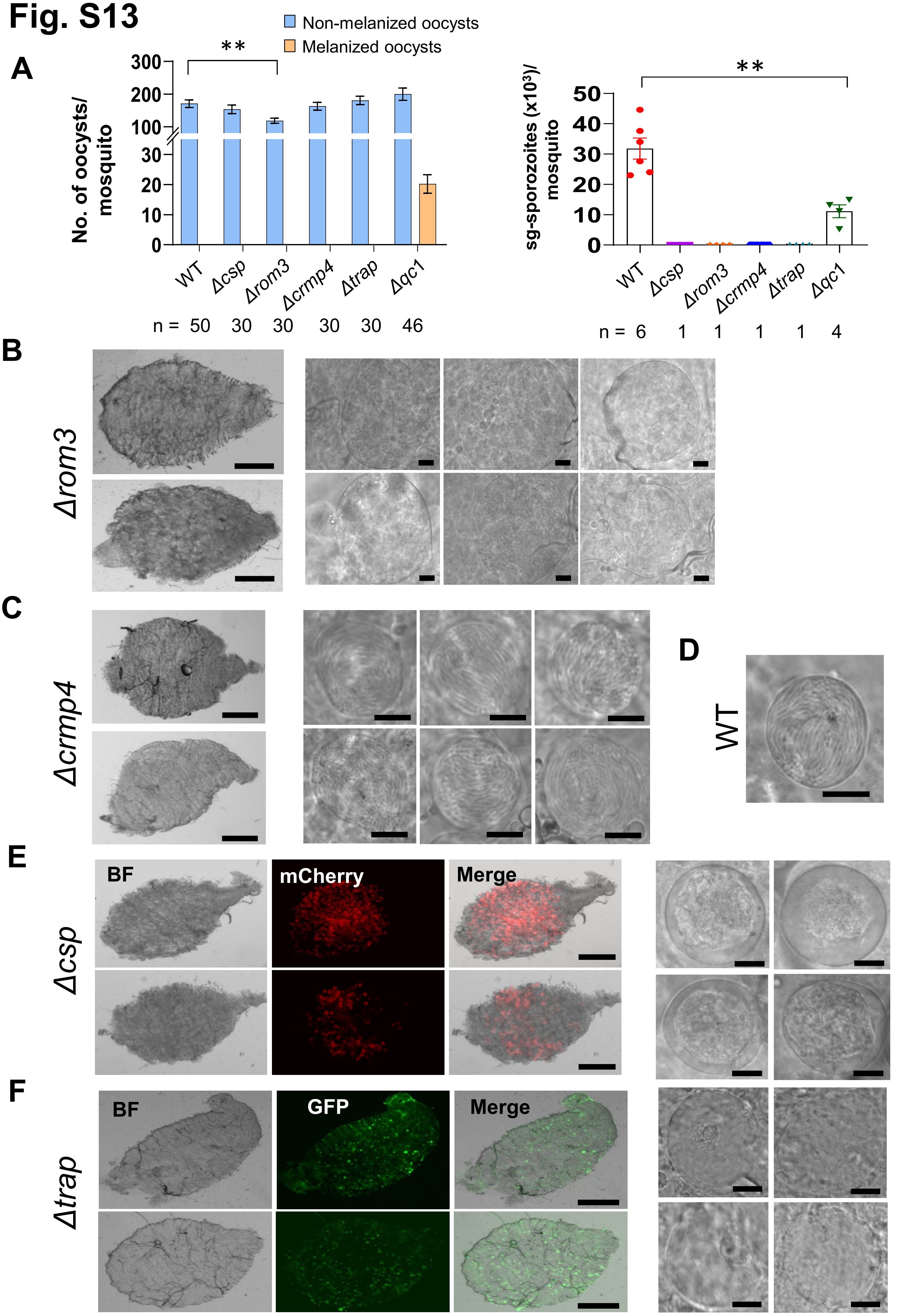

A
Non-melanized oocysts
Melanized oocysts
No. of oocysts/ mosquito
300
200
100
30
20
10
0
**
WT
Δcsp
Δrom3
Δcrmp4
Δtrap
Δqc1
n = 50 30 30 30 30 46
sg-sporozoites (x10³)/ mosquito
50
40
30
20
10
0
**
WT
Δcsp
Δrom3
Δcrmp4
Δtrap
Δqc1
n = 6 1 1 1 1 4
B
Δrom3
C
Δcrmp4
D
WT
E
Δcsp
BF
mCherry
Merge
F
Δtrap
BF
GFP
Merge

**Fig. S13. Absence of melanized oocysts or sporozoites at day 23 in mosquitoes infected with four *P. berghei* mutants with aberrant oocyst and/or sporozoite formation**

**A.** Left panel: Number of melanized (dark-coloured) and non-melanized oocysts per mosquito (mean $\pm$ SEM) in *A. stephensi* mosquitoes (n=number of mosquitoes) infected with a QC-null mutant (*Δqc1*), wild type (WT) and mutants *Δrom3*, Δ*crmp4*, Δ*csp*, and *Δtrap* (**P<0.05, Mann-Whitney test; statistical significance is shown relative to WT. P values are shown in **Table S3**). Right panel: Number of salivary gland (sg) sporozoites per mosquito (n=number of experiments; 60-80 mosquitoes/experiment) infected with the different parasites as shown in the left panel. (**P<0.05, Mann-Whitney test; statistical significance is shown relative to WT. P values are shown in **Table S3**

**B.** Midguts of mosquitoes with ROM3-null oocysts (*Δrom3*; line 430cl1). ROM3-null oocysts are enlarged compared to wildtype oocysts without any signs of sporulation (sporozoite formation). Scale bar: 200 µm (left panel) and 20 µm (right panel).

**C.** Midguts of mosquitoes CRMP4-null oocysts (*Δcrmp4*; line 376cl1). CRMP4-null oocysts form sporozoites but do not rupture and release sporozoites. Scale bar: 200 µm (left panel) and 20 µm (right panel).

**D.** WT mature oocyst containing sporozoites Scale bar: 20 µm

**E.** Midguts of mosquitoes with mCherry-expressing CSP-null oocysts (*Δcsp*; line 3065cl1). CSP-null oocysts are vacuolated without signs of sporulation (sporozoite formation). Scale bar: 200 µm (left panel) and 20 µm (right panel).

**F.** Midguts of mosquitoes infected with GFP-expressing TRAP-null oocysts (*Δtrap*; 2564cl3). TRAP-null oocysts form sporozoites that are released in the hemocoel but more than 90% of the sporozoites are unable to invade the salivary glands. Scale bar: 200 µm (left panel) and 20 µm (right panel).

#### Fig. S14

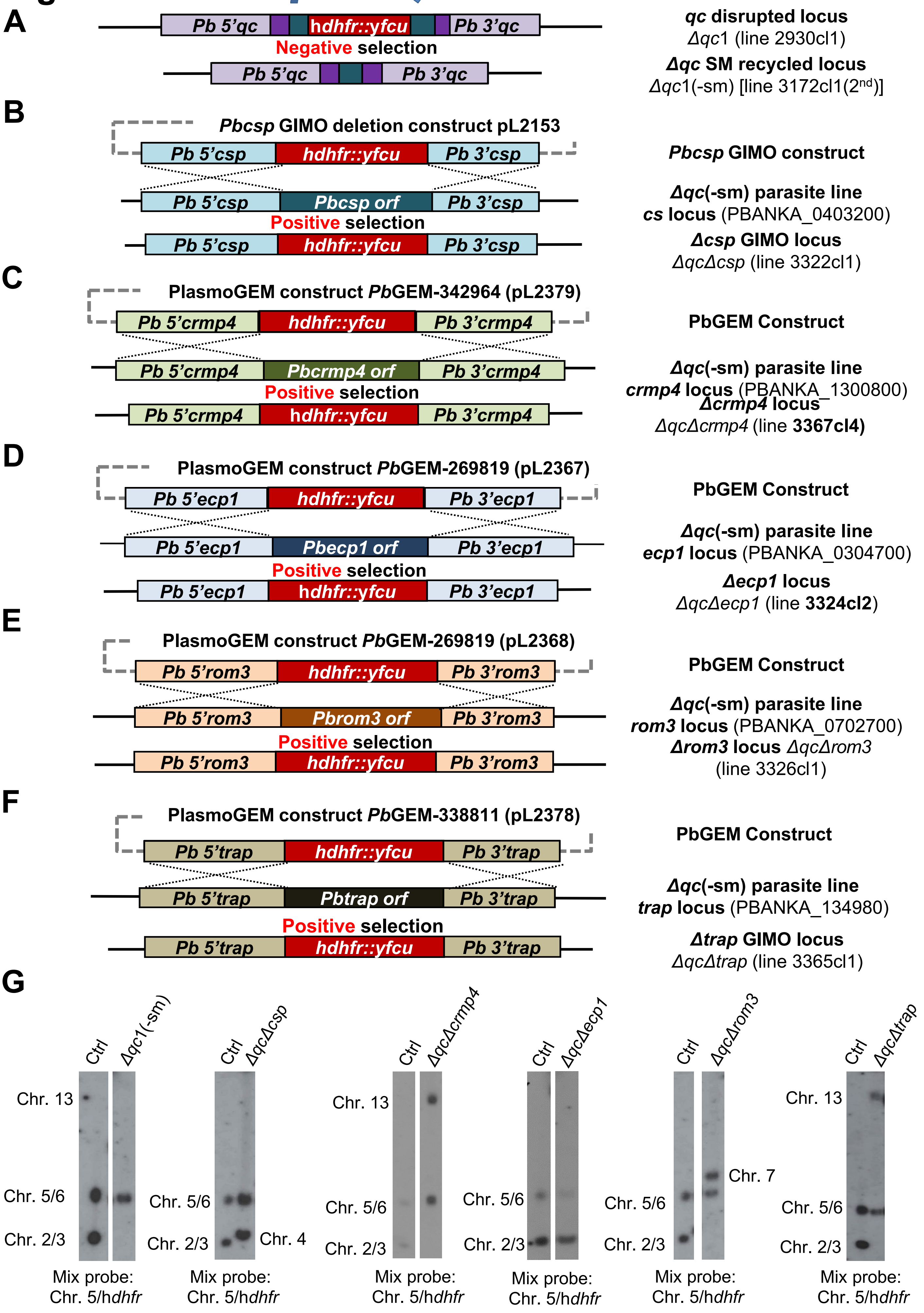

A
Pb 5'qc
hdhfr::yfcu
Pb 3'qc
Negative selection
Pb 5'qc
Pb 3'qc
qc disrupted locus
Δqc1 (line 2930cl1)
Δqc SM recycled locus
Δqc1(-sm) [line 3172cl1(2nd)]
B
Pbcsp GIMO deletion construct pL2153
Pb 5'csp
hdhfr::yfcu
Pb 3'csp
Pbcsp GIMO construct
Pb 5'csp
Pbcsp orf
Pb 3'csp
Δqc(-sm) parasite line
cs locus (PBANKA_0403200)
Positive selection
Pb 5'csp
hdhfr::yfcu
Pb 3'csp
Δcsp GIMO locus
ΔqcΔcsp (line 3322cl1)
C
PlasmoGEM construct PbGEM-342964 (pL2379)
Pb 5'crmp4
hdhfr::yfcu
Pb 3'crmp4
PbGEM Construct
Pb 5'crmp4
Pbcrmp4 orf
Pb 3'crmp4
Δqc(-sm) parasite line
crmp4 locus (PBANKA_1300800)
Positive selection
Δcrmp4 locus
Pb 5'crmp4
hdhfr::yfcu
Pb 3'crmp4
ΔqcΔcrmp4 (line 3367cl4)
D
PlasmoGEM construct PbGEM-269819 (pL2367)
Pb 5'ecp1
hdhfr::yfcu
Pb 3'ecp1
PbGEM Construct
Pb 5'ecp1
Pbecp1 orf
Pb 3'ecp1
Δqc(-sm) parasite line
ecp1 locus (PBANKA_0304700)
Positive selection
Δecp1 locus
Pb 5'ecp1
hdhfr::yfcu
Pb 3'ecp1
ΔqcΔecp1 (line 3324cl2)
E
PlasmoGEM construct PbGEM-269819 (pL2368)
Pb 5'rom3
hdhfr::yfcu
Pb 3'rom3
PbGEM Construct
Pb 5'rom3
Pbrom3 orf
Pb 3'rom3
Δqc(-sm) parasite line
rom3 locus (PBANKA_0702700)
Positive selection
Δrom3 locus ΔqcΔrom3
Pb 5'rom3
hdhfr::yfcu
Pb 3'rom3
(line 3326cl1)
F
PlasmoGEM construct PbGEM-338811 (pL2378)
PbGEM Construct
Pb 5'trap
hdhfr::yfcu
Pb 3'trap
Pb 5'trap
Pbtrap orf
Pb 3'trap
Δqc(-sm) parasite line
trap locus (PBANKA_134980)
Positive selection
Δtrap GIMO locus
Pb 5'trap
hdhfr::yfcu
Pb 3'trap
ΔqcΔtrap (line 3365cl1)
G
Ctrl
Δqc1(-sm)
Chr. 13
Chr. 5/6
Chr. 2/3
Mix probe:
Chr. 5/hdhfr
Ctrl
ΔqcΔcsp
Chr. 5/6
Chr. 2/3
Chr. 4
Mix probe:
Chr. 5/hdhfr
Ctrl
ΔqcΔcrmp4
Chr. 13
Chr. 5/6
Chr. 2/3
Mix probe:
Chr. 5/hdhfr
Ctrl
ΔqcΔecp1
Chr. 5/6
Chr. 2/3
Mix probe:
Chr. 5/hdhfr
Ctrl
ΔqcΔrom3
Chr. 7
Chr. 5/6
Chr. 2/3
Mix probe:
Chr. 5/hdhfr
Ctrl
ΔqcΔtrap
Chr. 13
Chr. 5/6
Chr. 2/3
Mix probe:
Chr. 5/hdhfr

**Fig. S14. Generation, genotyping and characterisation of the *P. berghei* 'double knockout' mutants lacking expression of *qc* and genes involved in sporozoite formation, sporozoite egress or invasion of salivary glands.**

**A.** Schematic representation of the generation of marker-free *PbΔqc*1 (*PbΔqc*1(-sm)) line. *PbΔqc*1 line contains the h*dhfr*::y*fcu* (red box) selectable marker (SM) cassette in the *Δqc* locus The SM cassette was removed by applying negative selection with 5-Fluorocytosine (5-FC), resulting in selection and cloning of SM-free *PbΔqc*1 parasites (3172cl1 (2nd)). In these parasites the SM cassette is removed by homologous recombination between the two 3'-UTR sequences (green boxes) present in the integrated construct in the *PbΔqc*1 genome that flank the h*dhfr*::y*fcu* SM cassette. The *PbΔqc*1(-sm) line was used to generate the 'double knock-out' mutants shown in **B-F**. **B.** Schematic representation of the generation of *ΔqcΔcsp* double knockout line (3322cl1). The GIMO deletion construct (pL2153) was used to replace the *Pbcsp orf* with the positive/negative SM (h*dhfr::yfcu*) cassette in *PbΔqc*1(-sm) line by double cross-over homologous recombination, resulting in the generation of *PbΔqcΔcsp* after positive selection with pyrimethamine and cloning. **C.** Schematic representation of the generation of *ΔqcΔcrmp4* double knockout line (3367cl4). The PlasmoGEM deletion construct (PbGEM-342964; pL2379) was used to replace the *Pbcrmp4 orf* with the positive/negative SM (h*dhfr::yfcu*) cassette in *PbΔqc*1(-sm) line by double cross-over homologous recombination, resulting in the generation of *PbΔqcΔcrmp4* after positive selection with pyrimethamine and cloning. **D.** Schematic representation of the generation of *ΔqcΔecp1* double knockout line (3324cl2). The PlasmoGEM deletion construct (PbGEM-269819) was used to replace the *Pbecp1 orf* with the positive/negative SM (h*dhfr::yfcu*) cassette in *PbΔqc*1(-sm) line by double cross-over homologous recombination, resulting in the generation of *PbΔqcΔexp1* after positive selection with pyrimethamine and cloning. **E.** Schematic representation of the generation of *ΔqcΔrom3* double knock-out line (3326cl1). The PlasmoGEM gene deletion construct (PbGEM-269819; pL2368) was used to replace the *Pbrom3 orf* with the positive/negative SM (h*dhfr::yfcu*) cassette in *PbΔqc*1(-sm) line by double cross-over homologous recombination, resulting in the generation of *PbΔqcΔrom3* after positive selection with pyrimethamine and cloning. **F.** Schematic representation of the generation of *ΔqcΔtrap* double knockout line (3365cl1). The PlasmoGEM deletion construct (PbGEM-338811; pL2378) was used to replace the *Pbtrap orf* with the positive/negative SM (h*dhfr::yfcu*) cassette in *PbΔqc*1(-sm) line by double cross-over homologous recombination, resulting in the generation of *PbΔqcΔtrap* after positive selection with pyrimethamine and cloning. **G.** Genotype analysis of *PbΔqcΔqc (-sm), ΔqcΔcsp*, *ΔqcΔcrmp4*, *ΔqcΔecp1, ΔqcΔrom3* and *PbΔqcΔtrap* parasites by Southern analysis of chromosomes (chr.) separated by pulsed-field gel electrophoresis (PFGE). Hybridisation of PFGE-separated chr. with a mixture of a *hdhfr* probe and a probe specific for chr. 5 confirms integration of pL2153 in the *Pbcsp* gene on chr. 4, pL2379 in the *Pbcrmp4* gene on chr. 13, pL2367 in the *Pbecp1* gene on chr. 3, pL2368 in the *Pbrom3* gene on chr. 7, and pL2378 in the *Pbtrap* gene on chr. 13. As an additional control (ctrl), parasite line 2117cl1 that has *hdhfr::yfcu* SM integrated into chr. 3. Blood stage growth of asexual blood stages was determined during the cloning period and was comparable to growth of wild type *P. berghei* ANKA parasites with a mean multiplication rate of 10x per 24 hour.

Fig. S15

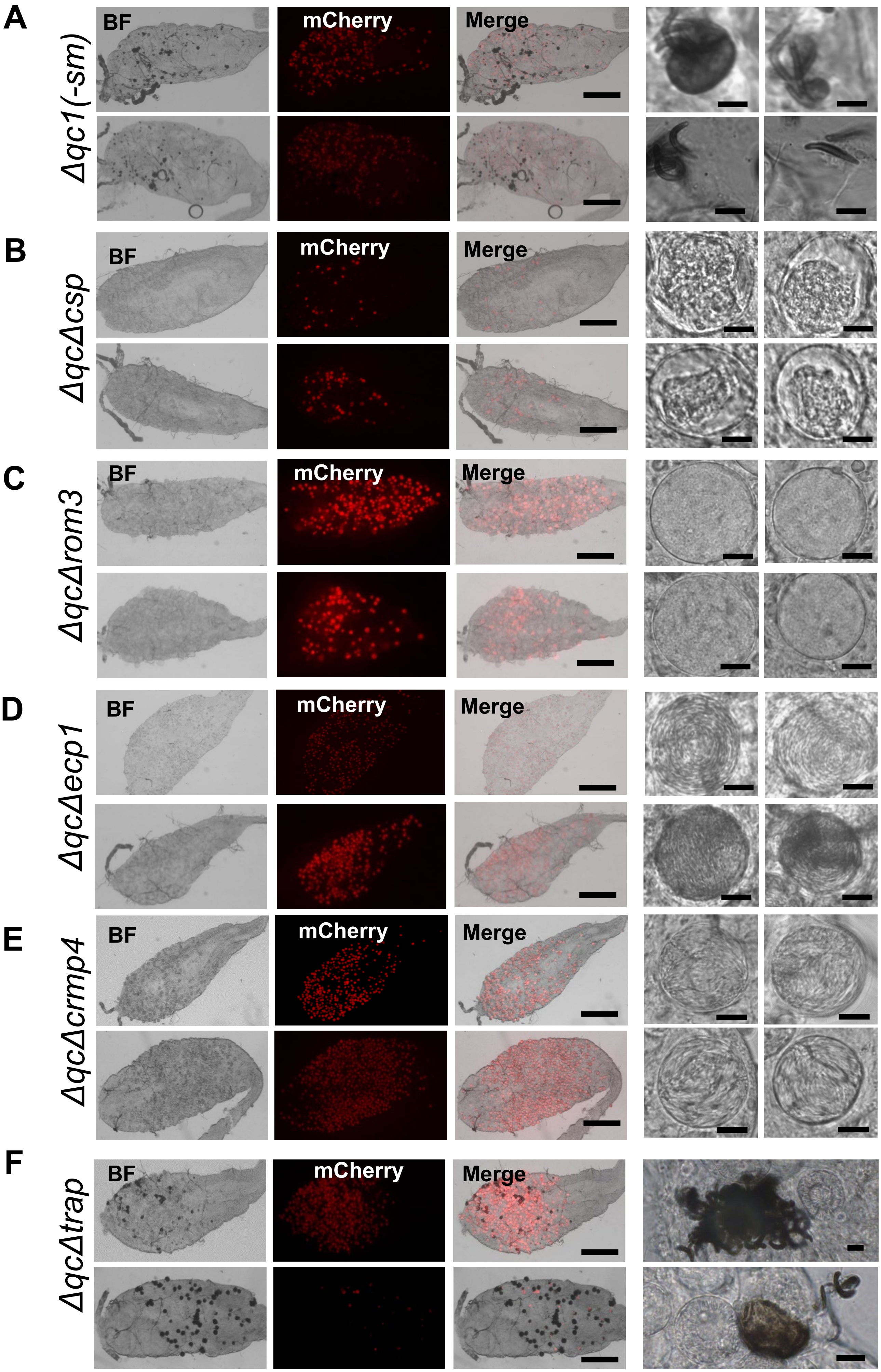

A
Δqc1(-sm)
B
ΔqcΔcsp
C
ΔqcΔrom3
D
ΔqcΔecp1
E
ΔqcΔcrmp4
F
ΔqcΔtrap
BF
mCherry
Merge

**Fig. S15. Absence of melanized oocysts and sporozoites at day 21 in mosquitoes infected with QC-null mutants lacking either expression of CSP, CRMP4 or ROM3 and melanized oocyst and sporozoites of a QC-null mutant lacking expression of TRAP**

**A.** Midguts with melanized (dark-coloured) oocyst (left 3 panels) and melanized oocysts and sporozoites (right 2 panels) of mosquitoes infected with mCherry-expressing QC-null mutant *Δqc1*(-*sm*) (3172cl1-2$^{nd}$). Scale bars: 200 µm (left panels) and 20 µm (right panels).

**B.** Midguts with oocyst (left 3 panels) and oocysts and sporozoites (right 2 panels) of mosquitoes infected with mCherry-expressing 'double knock-out' mutant Δ*qc*Δ*csp* (line 3322cl1), lacking expression of QC and CSP. CSP-null oocysts are vacuolated without signs of sporulation (sporozoite formation). Scale bars: 200 µm (left panels) and 20 µm (right panels).

**C.** Midguts with oocyst (left 3 panels) and oocysts and sporozoites (right 2 panels) of mosquitoes infected with mCherry-expressing 'double knock-out' mutant Δ*qc*Δ*rom3* (line 3326cl1), lacking expression of QC and ROM3. ROM3-null oocysts are enlarged compared to wildtype oocysts without any signs of sporulation (sporozoite formation). Scale bars: 200 µm (left panels) and 20 µm (right panels).

**D.** Midguts with oocyst (left 3 panels) and oocysts and sporozoites (right 2 panels) of mosquitoes infected with mCherry-expressing 'double knock-out' mutant Δ*qc*Δ*ecp1* (line 3324cl2), lacking expression of QC and ECP1. ECP1-null oocysts form sporozoites but do not rupture and release sporozoites. Scale bars: 200 µm (left panels) and 20 µm (right panels).

**E.** Midguts with oocyst (left 3 panels) and oocysts and sporozoites (right 2 panels) of mosquitoes infected with mCherry-expressing 'double knock-out' mutant Δ*qc*Δ*crmp4* (line 3367cl4), lacking expression of QC and CRMP4. CRMP4-null oocysts form sporozoites but do not rupture and release sporozoites. Scale bars: 200 µm (left panels) and 20 µm (right panels).

**F.** Midguts with melanized (dark-coloured) oocyst (left 3 panels) and melanized oocysts and sporozoites (right 2 panels) of mosquitoes infected with mCherry-expressing 'double knock-out' mutant Δ*qc*Δ*trap* (line 3365cl1), lacking expression of QC and TRAP. TRAP-null oocysts form sporozoites that are released in the hemocoel but more than 90% of the sporozoites are unable to invade the salivary glands. Scale bars: 200 µm (left panel) and 20 µm (right panel).

### Fig. S16

**A**

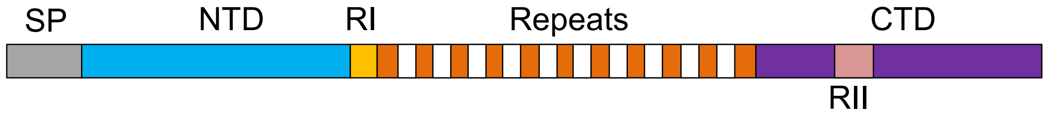


**B**

| Peptide sequence | # PSMs |
|---|---|
| QGNGQGHNMPNDPNR | 1 |
| QPADGNPDPNANPN | 3 |
| QPADGNPDPNANPNVD | 6 |
| QPADGNPDPNANPNVDPN | 12 |
| QPADGNPDPNANPNVDPNANPNVD | 3 |
| QPADGNPDPNANPNVDPNANPNVDPN | 3 |
| QPADGNPDPNANPNVDPNANPNVDPNANPNVD | 3 |
| QPADGNPDPNANPNVDPNANPNVDPNANPNVDPNANPN | 2 |

pGlu peptides with 'Q' from Region I of CSP

**C**

MMRKLAILSVSSFLFVEALFQEYQCYGSSSNTRVLNELNYDNAGTNLYNELEMNYYGKQENWYSLKK
NSRSLGENDDGNNEDNEKLRKPKHKKLKQPADGNPDPNANPNVDPNANPNVDPNANPNVDPNANPNA
NPNANPNANPNANPNANPNANPNANPNANPNANPNANPNANPNANPNANPNANPNANPNANPNVDPN
ANPNANPNANPNANPNANPNANPNANPNANPNANPNANPNANPNANPNANPNANPNANPNANPNANP
NANPNKNNQGNGQGHNMPNDPNRNVDENANANSAVKNNNNEEPSDKHIKEYLNKIQNSLSTEWSPCS
VTCGNGIQVRIKPGSANKPKDELDYANDIEKKICKMEKCSSVFNVVNSSIGLIMVLSFLFLN

**Fig. S16. Analysis of published mass spec data of *P. falciparum* circumsporozoite protein (CSP) for the presence of N-terminal pGlu**

**A.** Schematic of circumsporozoite protein showing different regions. SP - signal peptide, NTD - N-terminal domain, RI – region I, RII – region II plus, CTD – C-terminal domain.

**B.** Peptide sequences and peptide spectrum matches (PSM) of each peptide identified by mass spec analysis of CSP of wild type *P. falciparum* sporozoites[14]. pGlu peptides with 'Q' from Region I of CSP are in red.

**C.** Amino acid sequence of *P. falciparum* circumsporozoite protein (CSP). Region I (KLKQP) highlighted in yellow. Position of the peptides with pGlu identified by mass spec analysis are in red and blue (see **B**).

### Fig. S17

**A**

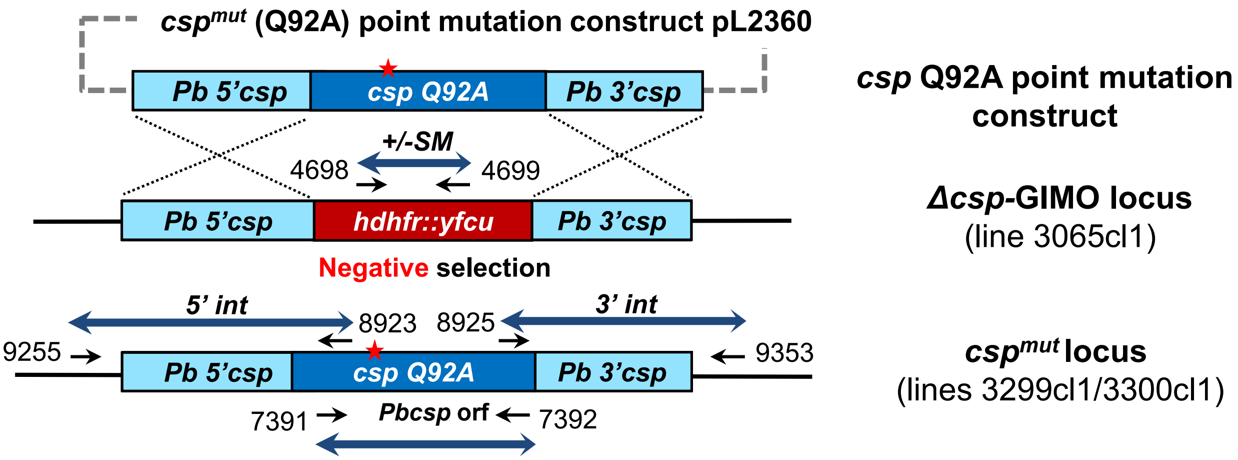


**B**

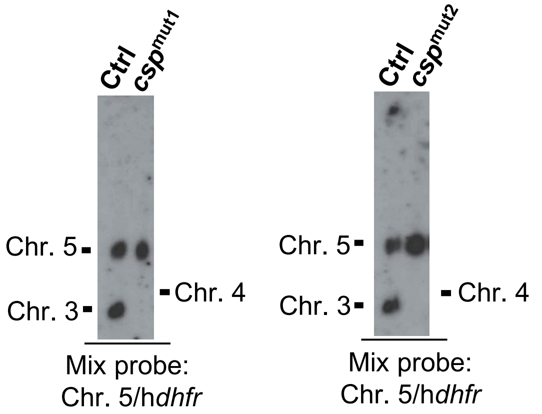


**C**

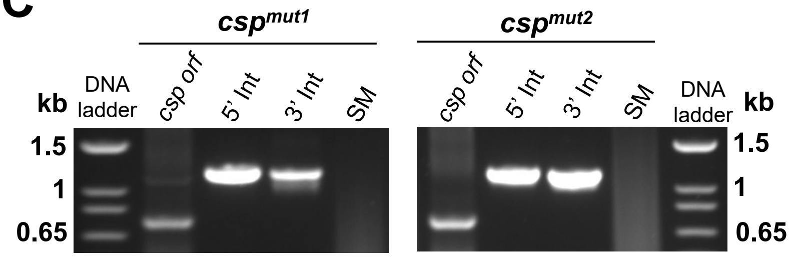


| Template | Product | Primers | | Amplicon size |
|---|---|---|---|---|
| ***csp***$^{mut1}$ | 5'int | 9255 | 8923 | 1199 bp |
| **(3299cl1)** | 3'int | 8925 | 9353 | 1181 bp |
| ***csp***$^{mut2}$ | SM | 4698 | 4699 | 1108 bp |
| **(3300cl1)** | *Pbcsp orf* | 7391 & 7392 | | 774 bp |

**D**

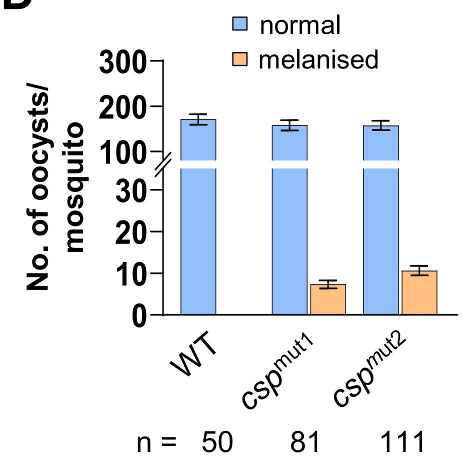


**E**

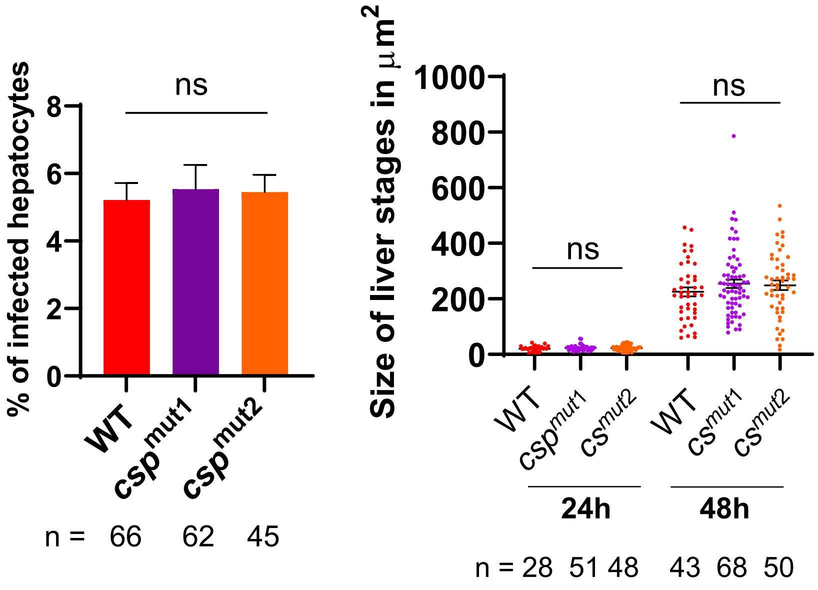

**Fig. S17. Generation and genotyping of *P. berghei* mutant expressing CSP with an amino acid replacement (mutation) in region I (*csp*$^{mut}$)**

**A.** Schematic representation of the generation of CSP mutant lines, *csp*$^{mut1}$ (3299cl1) and *csp*$^{mu2}$ (3300cl1), that express a mutated *csp* gene with the glutamine in region I replaced with an alanine (Q92A; red star in the schematic). The construct containing the mutated *csp* (pL2360) was used to replace the positive/negative selectable marker (SM; h*dhfr::yfcu*) cassette in the *Δcsp*-GIMO (3065cl1) line after transfection and applying negative selection with 5-Fluorocytosine (5-FC). Construct pL2360 integrates by double cross-over homologous recombination using *Pbcsp* targeting regions resulting in the replacement of the SM cassette by *csp* (Q92A) open reading frame (*orf*) under the control of *csp* regulatory sequences. Black arrows: location and primer names used for diagnostic PCR (see **D**).

**B.** Genotype analysis of *csp*$^{mut1}$ and *csp*$^{mut2}$ parasites by Southern analysis of chromosomes (chr.) separated by pulsed-field gel electrophoresis (PFGE). Hybridisation of PFG-separated chr. of *csp*$^{mut1}$ and *csp*$^{mut2}$ with a mixture of *hdhfr* and a probe specific for chr. 5 confirms the integration of pL2360 into the *csp*-GIMO locus on chr. 4. As an additional control (ctrl), parasite line 2117cl1 is used with the *hdhfr::yfcu* SM integrated into chr. 3.

**C.** Genotype analysis of *csp*$^{mut1}$ and *csp*$^{mut2}$ parasites by diagnostic PCRs confirms the absence of the *hdhfr::yfcu* SM, the correct integration of the construct at both the 5' and 3' regions (5'int and 3'int) and the presence of the *csp orf* (Q92A). See **A** for primer numbers and locations. Primer sequences are shown in **Table S1** and expected PCR fragment sizes and the primer numbers are listed in the Table.

**D.** Number of melanized (dark-coloured) and non-melanized oocysts per mosquito (mean $\pm$ SEM) in *A. stephensi* mosquitoes (n=number of mosquitoes) infected with two mutants (*csp*$^{mut1}$, *csp*$^{mut2}$) expressing a mutated CSP with the glutamine in region I replaced with an alanine (Q92A) and with WT parasites. ns-not significant (Mann-Whitney test, statistical significance is shown relative to WT. P values in **Table S3**)

**E.** Infectivity of salivary gland (sg) sporozoites of *csp*$^{mut1}$ and *csp*$^{mut2}$ lines *in vitro*. Left panel: percentage of infected hepatocytes (n=number of microscopic fields at 40X magnification). No significant (ns) differences were observed between *csp*$^{mut}$ and WT parasites (Mann-Whitney test, statistical significance is shown relative to WT. P values are shown in **Table S3**). Right panel: sizes (mean ± SEM) of *csp*$^{mut1}$, *csp*$^{mut2}$ and WT liver stages (n=number of liver stages) in cultured Huh7 hepatocytes at 24 and 48 h after infection as measured by determining the area of the parasite at its greatest circumference using the mCherry-positive area ($\mu m^2$). No significant (ns) differences in size were observed between *csp*$^{mut}$ and WT liver stages (Tukey's multiple comparisons test, statistical significance is shown relative to WT. P values are shown in **Table S3**).

**Table S1: List of primers used**

| Primer code | Sequence (5' to 3') | Enzymes | Product (bp) | Primer description |
|---|---|---|---|---|
| **Primers used to generate constructs** | | | | |
| ***P. berghei qc*** **GIMO construct (pL2335)** | | | | |
| 9641 | GACAAGCTTCCAAATCAGCCTTAAAAGATAAGTCCA | HindIII | 1100 | *Pbqc GIMO* HR1 forward |
| 9642 | CATCTGCAGATTTCTCAATAATCTTATTTGACTCTCCCC | PstI | | *Pbqc GIMO* HR1 reverse |
| 9643 | TATGGTACCGGGAAATATGAAAATTGTTTATGTAC | KpnI | 1058 | *Pbqc GIMO* HR2 forward |
| 9644 | ACGGAATTCGCTAGTGGGTGCAACGTTATTTTATT | EcoRI | | *Pbqc GIMO* HR2 reverse |
| ***Pfqc*** **complementation construct (pL2326)** | | | | |
| 9483 | AAGCTCGAGCCAAATCAGCCTTAAAAGATAAGTCCA | XhoI | 1100 | *Pbqc* HR1 forward |
| 9484 | CATGGATCCATTTCTCAATAATCTTATTTGACTCTCCCC | BamHI | | *Pbqc* HR1 reverse |
| 9485 | TATGAATTCGGGAAATATGAAAATTGTTTATGTAC | EcoRI | 1058 | *Pbqc* HR2 forward |
| 9486 | ACGCCGCGGGCTAGTGGGTGCAACGTTATTTTATT | SacII | | *Pbqc* HR2 reverse |
| 9487 | CAGGGATCCATGGAAAATGATAAATTAATAAATAA | BamHI | 2225 | *Pfqc orf* forward |
| 9488 | CCCGAATTCATATTTTGTTACCTATCGTTTTATAAT | EcoRI | | *Pfqc orf* reverse |
| ***Pbqc*** **complementation construct (pL2327)** | | | | |
| 9483 | AAGCTCGAGCCAAATCAGCCTTAAAAGATAAGTCCA | XhoI | 4184 | *Pbqc* HR1 forward |
| 9486 | ACGCCGCGGGCTAGTGGGTGCAACGTTATTTTATT | SacII | | *Pbqc* HR2 reverse |
| ***Pbqc*** **cmyc tagging construct (pL2351)** | | | | |
| 9233 | GTAGCGGCCGCATGGCAAATAAATTAGTTGCGAG | NotI | 2023 | *Pbqc orf* full length forward |
| 9234 | GTCGGATCCATGGCGATGTCTAATAGTTTTAAAATATTC | BamHI | | *Pbqc orf* full length reverse |
| ***Pbqc*** **enzyme dead mutant construct (pL2348)** | | | | |
| 9483 | AAGCTCGAGCCAAATCAGCCTTAAAAGATAAGTCCA | XhoI | 2135 | *Pbqc* HR1 forward |
| 9515 | ATAATAAACCTGCTGTAGCAGGATAATTGCTTATATATC | | | *Pbqc orf* mutation reverse |
| 9516 | CAATTATCCTGCTACAGCAGGTTTATTATATTTAGACGGG | | 2096 | *Pbqc orf* mutation forward |
| 9486 | ACGCCGCGGGCTAGTGGGTGCAACGTTATTTTATT | SacII | | *Pbqc* HR2 reverse |
| **Plasmid pLf0103** | | | | |
| 8801 | CTCCTCCTCGAGCGCATAAATATCTGGTGAAATACAA | XhoI | 970 | *Pfhsp70* promoter forward |
| 8802 | GGTGGTGGTACCGAACCTTTTGCACTAGCCAATTTTT | KpnI | | *Pfhsp70* promoter reverse |
| 8803 | CCTCCTCCTAGGATTTAATAATAGATTAAAAATATTA | AvrII | 590 | *Pfhrp2 3'utr* forward |
| 8804 | GCGGGCGGGCGGCCGCTTTAATAAATATGTTCTTATATATA | NotI | | *Pfhrp2 3'utr* reverse |

| | | | | |
|---|---|---|---|---|
| ***P. falciparum qc* KO construct (pLf0149)** | | | | |
| 9211 | AAGAAGAAGCTTAAAAGAATAAGAAAATAAGGAACCTTTTAA | HindIII | 798 | *Pfqc* HR1 forward |
| 9212 | GGGGGGCCCGGCGCGCCATTTATTTTGTTTTAATTAATTTTTGTTTTTC | ApaI, AscI | | *Pfqc* HR1 reverse |
| 9250 | CGTGCTGCTAGCATATCCCACCCAACTTAAAAGATACAG | NheI | 1232 | *Pfqc* HR2 forward |
| 9214 | GGAGGAGGATCCAAAACAACACAGATAAATTAACACACA | BamHI | | *Pfqc* HR2 reverse |
| **Guide RNA oligos for generating Cas9 targeting plasmids (pLf0150 and pLf0151)** | | | | |
| 9251 | TATTGGACTCCAAAGAAGAAACAC | | | *Pfqc* gRNA062 forward |
| 9252 | AAACGTGTTTCTTCTTTGGAGTCC | | | *Pfqc* gRNA062 reverse |
| 9253 | TATTCAGAAATTATGGGCAACATC | | | *Pfqc* gRNA063 forward |
| 9254 | AAACGATGTTGCCCATAATTTCTG | | | *Pfqc* gRNA063 reverse |
| **Primers used for genotyping** | | | | |
| ***PbΔqc*1 and *PbΔqc*2 lines (2930cl1 and 2931cl1)** | | | | |
| 8989 | ACGAATACAAGAAATGAATGGAAAAGTTGATC | | 5419 | 5’ *Pbqc* integration forward |
| 4494 | GGGTTGGGTGACTTTGGTGACAG | | | 5’ *Pbqc* integration reverse |
| 7289 | TAAAGCACAATATCTAGGATACTAC | | 3866 | 3’ *Pbqc* integration forward |
| 8990 | CGCCGACATCGCTATCTGCTCCATCATGCC | | | 3’ *Pbqc* integration reverse |
| 4698 | GTTCGCTAAACTGCATCGTC | | 1108 | SM forward |
| 4699 | GTTTGAGGTAGCAAGTAGACG | | | SM reverse |
| 8991 | GGGTGGTGATCCTTGTGATAATCATTTG | | 624 | *Pbqc orf* confirmation forward |
| 8992 | GTGGAATCTCTAATTCGTATAAAAATTCATTTCC | | | *Pbqc orf* confirmation reverse |
| ***PbΔqc* GIMO line (3198cl1)** | | | | |
| 9585 | CCTATATTGTTTTAAAGGTGCGAATTGCCCT | | 1540 | 5’ *Pbqc* integration forward |
| 5696 | TTACTGGTGCTTTGAGGGGTG | | | 5’ *Pbqc* integration reverse |
| 7922 | GTCTCTTCAATGATTCATAAATAGTTGG | | 1398 | 3’ *Pbqc* integration forward |
| 9588 | GATCCAGAAAGTATTAATAGAAAGGTTTATTGG | | | 3’ *Pbqc* integration reverse |
| 8987 | GTGCCAAATAAGTGTAAGCCAATATCCC | | 1492 | *Pbqc orf* confirmation forward |
| 8992 | GTGGAATCTCTAATTCGTATAAAAATTCATTTCC | | | *Pbqc orf* confirmation reverse |
| ***Pfqc* complementation in *PbΔqc* GIMO line (3213cl1)** | | | | |
| 9598 | ATGTATGCATTAAAACATATGTATGTAGTAAATC | | 725 | *Pfqc orf* confirmation forward |
| 9487 | CAGGGATCCATGGAAAATGATAAATTAATAAATAA | | | *Pfqc orf* confirmation reverse |
| 9585 | CCTATATTGTTTTAAAGGTGCGAATTGCCCT | | 1481 | *5’ Pbqc* integration forward |
| 9589 | CAATTTTATCCAGTGTTTCTTCTTTGGAGTCC | | | *5’ Pfqc* integration reverse |
| 9590 | ACCTTGCTAGTTACCGGAAAATTGTGGTCC | | 1386 | *3’ Pfqc* integration forward |
| 9588 | GATCCAGAAAGTATTAATAGAAAGGTTTATTGG | | | *3’ Pbqc* integration reverse |

| | | | |
|---|---|---|---|
| 9233<br>8992 | GTAGCGGCCGCATGGCAAATAAATTAGTTGCGAG<br>GTGGAATCTCTAATTCGTATAAAAATTCATTTCC | 1424 | *Pbqc orf* confirmation forward<br>*Pbqc orf* confirmation reverse |
| ***Pbqc* complementation in *PbΔqc* GIMO line (3216cl1) and $Pbqc^{CD}$ *Pbqc* cyclase dead mutant line (3272)** | | | |
| 8987<br>8992 | GTGCCAAATAAGTGTAAGCCAATATCCC<br>GTGGAATCTCTAATTCGTATAAAAATTCATTTCC | 1492 | *Pbqc orf* confirmation forward<br>*Pbqc orf* confirmation reverse |
| 9585<br>9586 | CCTATATTGTTTTAAAGGTGCGAATTGCCCT<br>AAATGGTGAGATGCTCTATATTTTTTCTCTC | 1698 | *5' Pbqc* integration forward<br>*5' Pbqc* integration reverse |
| 9587<br>9588 | AGCAAAACATGGAAAGAAATAGGGAATCC<br>GATCCAGAAAGTATTAATAGAAAGGTTTATTGG | 1460 | *3' Pbqc* integration forward<br>*3' Pbqc* integration reverse |
| **To confirm point mutations in the $Pbqc^{CD}$ mutant line (3272) by Sanger's sequencing** | | | |
| 8991<br>8992 | GGGTGGTGATCCTTGTGATAATCATTTG<br>GTGGAATCTCTAATTCGTATAAAAATTCATTTCC | 624 | *Pbqc orf* forward<br>*Pbqc orf* reverse |
| pJET FP<br>pJET RP | CGACTCACTATAGGGAGAGCGGC<br>AAGAACATCGATTTTCCATGGCAG | | *pJET* forward<br>*pJET* reverse |
| ***Pfqc orf* Southern probe** | | | |
| 9597<br>9598 | CATGCAAGAAAATATTATTATGTAGAAATGTACC<br>ATGTATGCATTAAAACATATGTATGTAGTAAATC | 444 | *Pfqc* southern probe forward<br>*Pfqc* southern probe reverse |
| ***Pbqc::cmyc* (3272)** | | | |
| 8987<br>5394 | GTGCCAAATAAGTGTAAGCCAATATCCC<br>GTTACACGTATATTACGCATACAACG | 2300 | *5' Pbqc-cmyc* integration forward<br>*5' Pbqc-cmyc* integration reverse |
| 5225<br>5861 | CAGGGTTTTCCCAGTCACGACGTTG<br>CCTTCAATTTCGGATCCACTAGCTTTTCACATCTTCAGTGG | 2830 | *3' Pbqc-cmyc* integration forward<br>*3' Pbqc-cmyc* integration reverse |
| ***PfΔqc*1 (*Pf* Exp.189) and *PfΔqc*2 (*Pf* Exp. 250) lines** | | | |
| 9393<br>9397 | TATATTTATGAACACCTGAAATATAAGTAAATTATTA<br>GTTTGTATTTCACCAGATATTTATGCG | 1678 | 5' *Pfqc* integration forward<br>5' *Pfqc* integration reverse |
| 9175<br>9396 | ATGAACATAAAGTACAACATTAATATATAGC<br>TTACAGTGGTTAATGTACAATGAAAAAAGG | 1600 | 3' *Pfqc* integration forward<br>3' *Pfqc* integration reverse |
| 9182<br>9183 | TCCACCCTCATTGAAAGAGCAA<br>CAATTCACGAATCCCAACTGCC | 342 | BSD forward<br>BSD reverse |
| 9251<br>9208 | TATTGGACTCCAAAGAAGAAACAC<br>AAACACGCCTTACGTATAAACCTA | 983 | *Pfqc orf* confirmation forward<br>*Pfqc orf* confirmation reverse |
| ***Pfqc orf* Southern probes** | | | |
| 9597<br>9598 | CATGCAAGAAAATATTATTATGTAGAAATGTACC<br>ATGTATGCATTAAAACATATGTATGTAGTAAATC | 444 | *Pfqc* southern probe forward<br>*Pfqc* southern probe reverse |

| | | | |
|---|---|---|---|
| 9193 | TTTGTATGTGCATGCGTACAAGA | 622 | *Pfbtrap* southern probe forward |
| 9274 | ACGACGACGCGTTGATTCTTTTAATGAATCTTGAATACTATCTGG | | *Pftrap* southern probe reverse |

#### Primers used for dsRNA-mediated gene silencing in *Anopheles stephensi* mosquitoes

| Primer code | Sequence (5' to 3') | | Primer description |
|---|---|---|---|
| LRIM1_ Ast814ExF | CTCGAGGTGCTGAATGTGTC | 517 | *LRIM1* external PCR forward |
| LRIM1_Ast814ExR | CGAGCTTGTTGTTGCTGAGA | | *LRIM1* external PCR reverse |
| LRIM1_Ast814InF | TAATACGACTCACTATAGGGGTTGGATGTTCGATCGCTCT | 445 | *LRIM1* nested PCR forward |
| LRIM1_Ast814InR | TAATACGACTCACTATAGGGAGCTTGTTCGACGAGAGGTC | | *LRIM1* nested PCR reverse |
| F_dsRNA_Asteph009395b | TACGGTGAGTATCCGTGGGT | 508 | *CLIPA8* external PCR forward |
| R_dsRNA_Asteph009395b | GAAGCCGAAAGAAGGGACCA | | *CLIPA8* external PCR reverse |
| F_T7nest_dsRNA_As009395b | TAATACGACTCACTATAGGAGTGGGATTTGACACGCGAA | 358 | *CLIPA8* nested PCR forward |
| R_T7nest_dsRNA_As009395b | TAATACGACTCACTATAGGGGTGACCGCAAATAACTGCT | | *CLIPA8* nested PCR reverse |

#### qPCR primers used to assess gene silencing

| | | |
|---|---|---|
| Aste_LRIM1_F | GGGTCCGTACTGCTGTGAAAA | *LRIM1* qPCR forward |
| Aste_LRIM1_R | TCCTTTTCCGAACCGACGCG | *LRIM1* qPCR reverse |
| F_qPCR_As009395b | AGTCTGGTTCGTACGTGCTG | *CLIPA8* qPCR forward |
| R_qPCR_As009395b | CCCTGCAGGTGGCAACTTAG | *CLIPA8* qPCR reverse |
| S7F | AGAACCAGCAGACCACCATC | Ribosomal protein *S7* forward |
| S7R | GCTGCAAACTTCGGCTATTC | Ribosomal protein *S7* reverse |

HR – Homology region; *orf* - open reading frame; utr – untranslated region SM - selectable marker

**Table S2. Gametocyte, ookinete and oocyst production of wild type *P. berghei* parasites (WT) and parasites of *Δqc* lines (Δ*qc*1, 2, G, 1(-sm)), complemented lines (*Pf*qc(c); *Pbq*c(c)); cmyc-tagged line (*Pbqc::cmyc*) and catalytically-dead mutant line (*Pbqc*$^{CD}$)**

| Mutant | Gametocyte production (%)[1] mean (s.d.) | Male exflagellation rate (%)[2] mean (s.d.) | Ookinete production (%)[3] mean (s.d.) | Number of oocysts[4] mean (s.d.) | Number of melanized oocysts[5] mean (s.d.) | Number of salivary gland sporozoites[6] (x10$^3$) |
|---|---|---|---|---|---|---|
| Δ*qc*1 | 17 (1), n=3 | 81 (3), n=3 | 81 (4), n=3 | 163 (118), n>10 | 26 (19), n>10 | 11.2; 5.2; 13.2 |
| Δ*qc*2 | 18 (1), n=3 | 88 (2), n=3 | 78 (3), n=3 | 222 (143), n>10 | 20 (16), n>10 | 3.4; 6.9; 7.3, 5.5 |
| Δ*qc*-G | 19 (1), n=3 | 93 (2), n=3 | 81 (3), n=3 | 141 (83), n>10 | 19 (14), n>10 | 12; 3.6 |
| *Pfqc*(c) | n.d. | n.d. | n.d. | 159 (118), n>10 | 0, n>10 | 25.7, 50.3, 44.8 |
| *Pbqc*(c) | n.d. | n.d. | n.d. | 235 (106), n>10 | 0, n>10 | 24, n=1 |
| *qc::cmyc* | n.d. | n.d. | n.d. | 245 (134), n>10 | 0, n>10 | 25, 18, 24 |
| *qc*$^{CD}$ | n.d. | n.d. | n.d. | 235 (138), n>10 | 18 (13), n>10 | 7.8, 13.1 |
| *Δqc*1(-sm) | n.d. | n.d. | n.d. | 142 (58), n>10 | 22 (13), n>10 | 11, n=1 |
| WT[7] | 15-25, *n*>10 | 65-95, n>10 | 50-90, *n*>10 | 75-249, n>10 | 0, n>10 | 23-45 , n>10 |

[1] The mean percentage of blood stage parasites developing into gametocytes *in vivo*
[2] The mean percentage of males that exflagellate *in vitro* 12-15 minutes after activation
[3] The mean percentage of female gametes developing into mature ookinetes *in vitro*16-18 hours after activation of gametocytes
[4] The mean number of oocysts per mosquito (days 12–13)
[5] The mean number of melanised oocysts per mosquito (days 14, 17 and 21)
[6] The mean number of sporozoites per mosquito (days 21)
[7] Wild type: reference *P. berghei* ANKA reporter line 1868cl1
s.d.: standard deviation
N.D.: not determined

#### Table S3: p-values

| Details of figure / graph | Comparison | P value |
|---|---|---|
| Fig. 1C: *in vitro* QC activity | WT-*Pf*QC vs $PfQC^{CD}$ | <0.0001 (****) |
| Fig. 2A: Number of oocysts in WT and different *qc*-mutant infected mosquitoes | WT vs *Δqc1* | 0.1590 (ns) |
| | WT vs *qc2* | 0.1736 (ns) |
| | WT vs *Δqc-G* | 0.1876 (ns) |
| | WT vs *Pbqc(c )* | 0.3600 (ns) |
| | WT vs *Pfqc (c)* | 0.0798 (ns) |
| | WT vs $qc^{CD}$ | 0.0617 (ns) |
| Fig. 2B: Number of salivary gland sporozoites produced in WT and different *qc*-mutant infected mosquitoes | WT vs *Δqc1* | 0.0095 (**) |
| | WT vs *qc2* | 0.0043 (**) |
| | WT vs *Δqc-G* | 0.0238 (*) |
| | WT vs *Pbqc(c )* | - |
| | WT vs *Pfqc (c)* | 0.2619 (ns) |
| | WT vs *qc::cmyc* | 0.1905 (ns) |
| | WT vs $qc^{CD}$ | 0.0714 (ns) |
| | WT vs *Δqc1* | 0.0095 (**) |
| | WT vs *qc2* | 0.0043 (**) |
| Fig. 4A: Number of oocysts in *Pf*WT, *PfΔqc* mutant infected mosquitoes | WT vs *Δqc1* | 0.3333 (ns) |
| | WT vs *Δqc2* | 0.5312 (ns) |
| Fig. 4B: Number of salivary gland sporozoites produced in *Pf*WT, *PfΔqc* mutant infected mosquitoes | WT vs *Δqc1* | 0.1333 (ns) |
| | WT vs *Δqc2* | 0.1333 (ns) |
| Fig. 6A (left panel): Number of oocysts in WT, *Δqc* and different double knockout mutant infected mosquitoes | WT vs *ΔqcΔcsp* | 0.0026 (**) |
| | WT vs *ΔqcΔrom3* | 0.9675 (ns) |
| | WT vs *ΔqcΔecp1* | 0.1836 (ns) |
| | WT vs *ΔqcΔcrmp4* | 0.2157 (ns) |
| | WT vs *ΔqcΔtrap* | 0.8948 (ns) |
| | WT vs *Δqc1(-sm)* | 0.0748 (ns) |
| Fig. 6A (right panel): Number of salivary gland sporozoites produced in WT, *Δqc* and different double knockout mutant infected mosquitoes | WT vs Δqc*Δcsp* | - |
| | WT vs Δqc*Δrom3* | - |
| | WT vs Δqc*Δecp1* | - |
| | WT vs Δqc*Δcrmp4* | - |
| | WT vs Δqc*Δtrap* | - |
| | WT vs *Δqc1(-sm)* | 0.0714 (ns) |

| | | |
|---|---|---|
| Fig. 6D (right panel): Number of salivary gland sporozoites produced in WT and *csp*$^{mut}$ parasite infected mosquitoes | WT vs *CS*$^{mut1}$ | 0.0727 (ns) |
| | WT vs *CS*$^{mut2}$ | 0.0734 (ns) |
| Extended data Fig. 5a: Number of oocysts in *Pf*WT, *PfΔqc* mutant infected mosquitoes | WT vs *Δqc1* | 0.3333 (ns) |
| | WT vs *Δqc2* | 0.5312 (ns) |
| Fig. S4E (left panel): RLU values in the livers of mice at 44h infected with WT and *Pbqc::cmyc* parasites | WT vs *Pbqc::cmyc* | 0.4000 (ns) |
| Fig. S4E (right panel): Pre-patency periods of WT and *Pbqc::cmyc* parasites in mice infected with $10^4$ sporozoites | WT vs *Pbqc::cmyc* | >0.9999 (ns) |
| Fig. S8B: Velocity of WT and *Δqc* mutant sporozoites | WT vs *Δqc1* | >0.9999 (ns) |
| | WT vs *Δqc2* | >0.9999 (ns) |
| Fig. S9B: Sizes of WT and *Δqc* mutant EEFs in cultured Huh7 hepatocytes at 24 and 48 hpi | WT 24h vs. *PbΔqc*1 24h | 0.3028 (ns) |
| | WT 24h vs. *PbΔqc*2 24h | 0.3284 (ns) |
| | WT 48h vs. *PbΔqc*1 48h | 0.5832 (ns) |
| | WT 48h vs. *PbΔqc*2 48h | 0.5085 (ns) |
| Fig. S9C: Infectivity of WT and *Δqc* mutant sporozoites in cultured Huh7 hepatocytes *in vitro* | WT vs *Δqc1* | 0.6837 (ns) |
| | WT vs *Δqc2* | 0.2810 (ns) |
| Fig. S12D: Growth of asexual blood-stages of *PfΔqc1, PfΔqc2* and WT *Pf*NF54 | WT vs *Δqc1* | 0.8593 (ns) |
| | WT vs *Δqc2* | 0.9805 (ns) |
| Fig. S13A: Number of oocysts in WT, *Δqc* and different single gene knockout mutant infected mosquitoes | WT vs *Δcs* | 0.4747 (ns) |
| | WT vs *Δrom3* | 0.0068 (**) |
| | WT vs *Δcrmp4* | 0.9075 (ns) |
| | WT vs *Δtrap* | 0.2661 (ns) |
| Fig. S13B: Number of salivary gland sporozoites produced in WT *Δqc* and different single gene knockout mutant infected mosquitoes | WT vs *Δqc1* | 0.0095(**) |
| Fig. S17D: Number of oocysts in WT and *csp*$^{mut}$ parasites infected mosquitoes | WT vs *csp*$^{mut1}$ | 0.4390 (ns) |
| | WT vs *csp*$^{mut2}$ | 0.2639 (ns) |
| Fig. S17E (left panel): Infectivity of WT and *csp*$^{mut}$ sporozoites in cultured Huh7 hepatocytes *in vitro* | WT 24h vs. *csp*$^{mut1}$ 24h | 0.4215 (ns) |
| | WT 24h vs. *csp*$^{mut2}$ 24h | 0.5395 (ns) |
| Fig. S17E (right panel): Sizes of WT and *Δqc* mutant EEFs in cultured Huh7 hepatocytes at 24 and 48 hpi | WT vs *cs*$^{mut1}$ | 0.9026 (ns) |
| | WT vs *cs*$^{mut2}$ | 0.4960 (ns) |
